# Multicohort Analysis of Publicly-available Monocyte Expression Data Identifies Gene Signatures to Accurately Monitor Subset-specific Changes in Human Diseases

**DOI:** 10.1101/2020.12.21.423397

**Authors:** Francesco Vallania, Liron Zisman, Claudia Macaubas, Shu-Chen Hung, Narendiran Rajasekaran, Sonia Mason, Jonathan Graf, Mary Nakamura, Elizabeth D Mellins, Purvesh Khatri

**Author notes:** Contributed equally (Elizabeth D Mellins, Purvesh Khatri). Correspondence: Purvesh Khatri or Elizabeth Mellins.

## Abstract

Monocytes and monocyte-derived cells play important roles in the regulation of inflammation, both as precursors as well as effector cells. Monocytes are heterogeneous and characterized by three distinct subsets in humans. Classical and non-classical monocytes represent the most abundant subsets, each carrying out distinct biological functions. Consequently, altered frequencies of different subsets have been associated with inflammatory conditions, such as infections and autoimmune disorders including lupus, rheumatoid arthritis, inflammatory bowel disease, and, more recently, COVID-19. Dissecting the contribution of different monocyte subsets to disease is currently limited by samples and cohorts, often resulting in underpowered studies and, consequently, poor reproducibility. Public transcriptomes provide an alternative source of data characterized by high statistical power and real world heterogeneity. However, most transcriptome datasets profile bulk blood or tissue samples, requiring the use of *in silico* approaches to quantify changes in the levels of specific cell types.

Here, we integrated 853 publicly available microarray expression profiles of sorted human monocyte subsets from 45 independent studies to identify robust and parsimonious gene expression signatures, consisting of 10 genes specific to each subset. These signatures, although derived using only datasets profiling healthy individuals, maintain their accuracy independent of the disease state in an independent cohort profiled by RNA-sequencing (AUC = 1.0). Furthermore, we demonstrate that our signatures are specific to monocyte subsets compared to other immune cells such as B, T, dendritic cells (DCs) and natural killer (NK) cells (AUC = 0.87~0.88, p<2.2e-16). This increased specificity results in estimated monocyte subset levels that are strongly correlated with cytometry-based quantification of cellular subsets (r = 0.69, p = 6.7e-4). Consequently, we show that these monocyte subset-specific signatures can be used to quantify changes in monocyte subsets levels in expression profiles from patients in clinical trials. Finally, we show that proteins encoded by our signature genes can be used in cytometry-based assays to specifically sort monocyte subsets. Our results demonstrate the robustness, versatility, and utility of our computational approach and provide a framework for the discovery of new cellular markers.

## Introduction

Monocytes, together with macrophages and dendritic cells (DCs), are part of the mononuclear phagocyte system. Monocytes and monocyte-derived cells play important roles in the regulation of inflammation, both as precursors as well as effector cells (1,2,3). Monocytes are a heterogeneous group of cells, and since Passlick and colleagues showed that the combined use of CD16 (FcγRIII) and the LPS co-receptor CD14 identified distinct subsets of monocytes in humans (4), three major subsets have been defined, as well as their murine counterparts. These subsets are: a classical (CD14+CD16-in humans and Ly6C^hi^ in mice), a nonclassical (CD14-CD16+ in humans and Ly6C^lo^ in mice), and an intermediate subset (CD14+CD16+ in humans and Ly6C+Treml4+ in mice) (5,6, 10). Although plasticity is an important characteristic of monocytes (7,8), surface marker expression and functional studies have shown each monocyte subset to be functionally distinct. Consistent with this, transcriptome analyses of sorted monocyte subsets have revealed different gene expression profiles ascribed to each subset (9,10,11). Classical monocytes, around 90% of total monocytes in humans, are efficient phagocytic cells important for the initiation of inflammatory response, with high expression of scavenger and chemokine receptors, and elevated cytokine production (12,13). Earlier work on nonclassical monocytes, around 5% of total monocytes, emphasized their capacity to produce inflammatory cytokines, especially TNF (14). However, recent work has shown that nonclassical monocytes are also involved in immune surveillance of the vasculature and have pro- and antiinflammatory functions (9,15). Intermediate monocytes are considered efficient antigen presenting cells (Kapellos et al, 2019). Consequently, altered frequencies of different subsets have been associated with inflammatory conditions, such as infections and autoimmune disorders including lupus, rheumatoid arthritis, and inflammatory bowel disease (13,16-19), and more recently, COVID-19 (20,21).

Dissecting the contribution of different monocyte subsets to disease is currently limited by samples and cohorts that can be profiled experimentally using cytometry and cell-staining-based assays. These limitations often result in underpowered studies and, consequently, poor reproducibility (22). Public transcriptomes provide an alternative source of data characterized by high statistical power and real-world biological, clinical, and technical heterogeneity, resulting in increased reproducibility (23-30). However, most transcriptome datasets profile bulk blood or tissue samples, requiring the use of *in silico* approaches to quantify changes in the levels of specific cell types (31-34).

Here, we integrated 853 publicly available microarray expression profiles of sorted human monocyte subsets from 45 independent studies to identify robust and parsimonious gene expression signatures, consisting of 10 genes specific to each subset. These signatures, although derived using only datasets profiling healthy individuals, maintain their accuracy independent of the disease state in an independent cohort profiled by RNA-sequencing. Furthermore, we demonstrate that these signatures are specific to monocyte subsets compared to other immune cells such as B, T, dendritic cells (DCs) and natural killer (NK) cells. This increased specificity results in estimated monocyte subset levels that are strongly correlated with cytometry-based quantification of cellular subsets. Consequently, we show that these monocyte subset-specific signatures can be used to quantify changes in monocyte subsets levels in expression profiles from patients in clinical trials. Finally, we show that proteins encoded by our signature genes can be used in cytometry-based assays to specifically sort monocyte subsets. Our results demonstrate the robustness, versatility, and utility of our computational approach and provide a framework for the discovery of new cellular markers.

## Results

### Robust subset-specific monocyte signatures from heterogeneous gene expression datasets

We hypothesized that integrating heterogeneous transcriptome profiles of sorted human monocyte subsets across multiple cohorts would allow us to identify robust subset-specific gene expression signatures. To test this hypothesis, we collected and annotated 853 publicly available gene expression profiles of sorted human monocytes across 45 studies. Our datasets spanned 22 microarray platforms for transcriptome profiling of samples acquired from healthy donors (**Supplementary Table 1**). After sample annotation, we co-normalized and integrated all expression data as previously described (31) (**Figure 1**). For each gene, we calculated effect sizes as Hedge’s *g* between the samples from a cell subset of interest compared to all other samples. We then characterized the underlying biological functions represented within the transcriptional data for each monocyte subset by performing Gene Set Enrichment Analysis (GSEA) (see **Methods**). We identified 91 significantly enriched pathways in classical monocytes and 1737 for the nonclassical subset (FDR < 5%). Our analysis revealed pathways associated with known functions of classical monocytes, such as wound healing, cytoskeleton remodeling, and phagocytosis, positively enriched in the classical subset (**Supplementary Table 2**). In contrast, our analysis of the nonclassical subset revealed categories of disease-associated gene expression changes, cell cycle, and metabolism (**Supplementary Table 3**). Notably, our most significant enrichment consisted of a gene signature previously reported to be down-regulated in Alzheimer’s Disease (pathway: ‘BLALOCK_ALZHEIMERS_DISEASE_DN’, p = 9.9e-6). This is in agreement with previous reports showing that nonclassical monocytes are found to be reduced in patients affected by Alzheimer’s (35). These results suggest that our data integration strategy allowed us to preserve and capture previously described biological functions of monocyte subsets irrespective of technical and biological confounders within our collection of datasets. We then applied our multi-cohort analysis framework to identify robust cell-subset specific genes (see Methods) (23,31,36). We considered classical and nonclassical monocyte subsets for our analysis because of their functional importance and the number of available datasets for each subset that could be integrated into our analysis (n >=4). There were 30 genes significantly over-expressed in classical monocytes and 268 genes overexpressed in nonclassical monocytes. We created classical and nonclassical monocyte specific gene signatures, using the top 10 genes for each monocyte subset that were consistently elevated within the subset of interest across all our discovery cohorts (**Figure 2A-B, Supplementary Table 4**). Among the classical signature genes, five have been previously associated with the classical monocytes in a single-cell RNA-seq study of healthy monocytes, and *SIGLEC10* was identified in non-classical monocytes as well (37).

**Figure 1:**
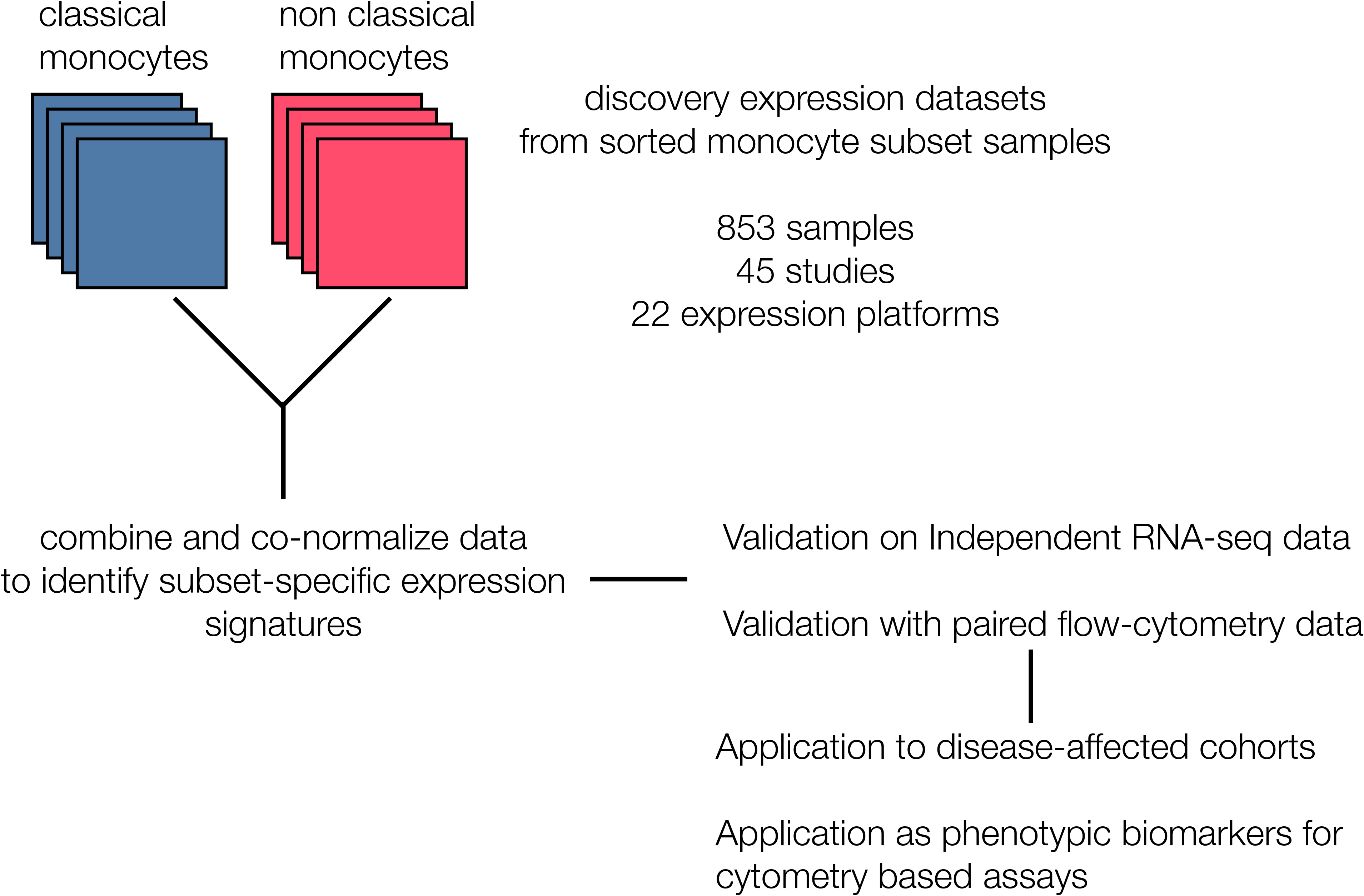
Generation of monocyte-specific signatures. Workflow depicting collection and annotation of publicly available discovery datasets from NCBI GEO profiling sorted human monocyte cell subsets (classical and non classical). Data was then combined and co-normalized to identify subset-specific signatures. Signatures were validated on an independent RNA-seq cohort, and on PBMCs expression profiles with paired flow data. After validation, signatures were applied on disease-affected cohorts and tested for their viability as phenotypic markers in cytometry-based assays.

**Figure 2:**
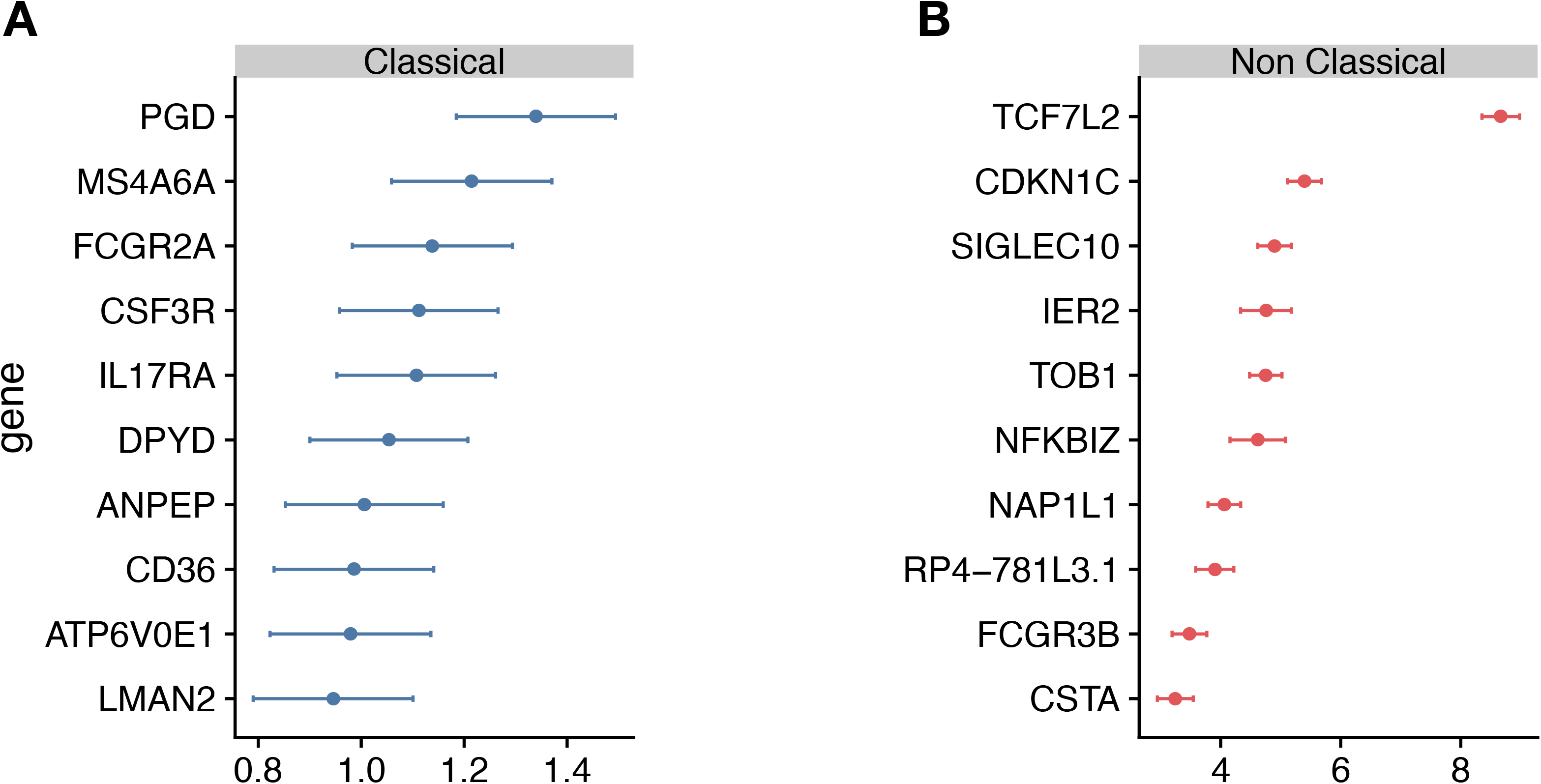
Identification of monocyte-subset specific gene expression signatures. (a) Forest plots displaying genes specific to the classical monocyte subset in the discovery cohort. Dots indicate *Hedges’ g* effect size values and bars correspond to their standard errors. (b) Same as (a) but for non classical subset.

### Monocyte signatures are robust in an independent validation cohort and independent of disease state

Although the gene signatures for classical and non-classical monocytes were derived by integrating independent datasets with substantial technical and biological heterogeneity, they only included healthy human subjects. We have previously shown that the accuracy of cell-type specific genes can be significantly affected by disease-induced changes in gene expression (31). Therefore, we investigated whether these monocyte subset-specific signatures are confounded by disease state in an independent cohort of healthy controls (n=3) and patients with rheumatoid arthritis (RA; n=3). We sorted monocyte subsets (classical, nonclassical, and intermediate) from peripheral blood samples and measured their transcriptomic profile using RNA-seq (see **Materials and Methods**).

Hierarchical clustering of the RNA-seq data using all genes in our monocyte subset-specific signatures accurately separated samples according to their cell subset identity, but not by their disease status (**Figure 3A**). Importantly, the signature genes showed variable expression levels in the intermediate monocytes, suggesting that the intermediate monocytes may represent a transitional cellular state rather than a stable state. All genes except two from the non-classical monocyte signature, (*IER2* and *CTSA*), were correctly over-expressed in their respective subset, and most (15 of 20) were independently confirmed by RT-PCR in sorted monocytes from healthy controls and patients with RA (**Supplementary Figure 1**). Next, we defined a classical monocyte subset score (cMSS) of a sample as a geometric mean of expression of genes in classical monocyte-specific signature, and nonclassical monocyte subset score (ncMSS) of a sample as a geometric mean of expression of genes in nonclassical monocyte-specific signature. We computed cMSS and ncMSS for each sample. In the independent cohort of healthy controls and patients with RA, we found that cMSS was higher in classical monocytes compared to nonclassical monocytes (p=3e-6), whereas ncMSS was higher in nonclassical monocytes compared to the classical subset (p=2.7e-6) (**Figure 3B,C**). Both scores identified their respective monocyte subset with high accuracy irrespective of the disease state of the sample (AUROC=1; **Supplementary Figure 2**), although *IER2* and *CTSA* were expressed in opposite directions. Importantly, cMSS and ncMSS for intermediate monocytes possessed scores between those of classical and non-classical monocytes (cMSS: intermediate vs classical p-value = 7.4e-4; intermediate vs non-classical p-value = 3.4e-4) (ncMSS: intermediate vs classical p-value = 1.5e-3; intermediate vs non-classical p-value = 9.9e-3) (**Figure 3B,C**). Overall, our results provide further independent evidence that our monocyte subset-specific signatures are consistently accurate across healthy and disease-affected samples.

**Figure 3:**
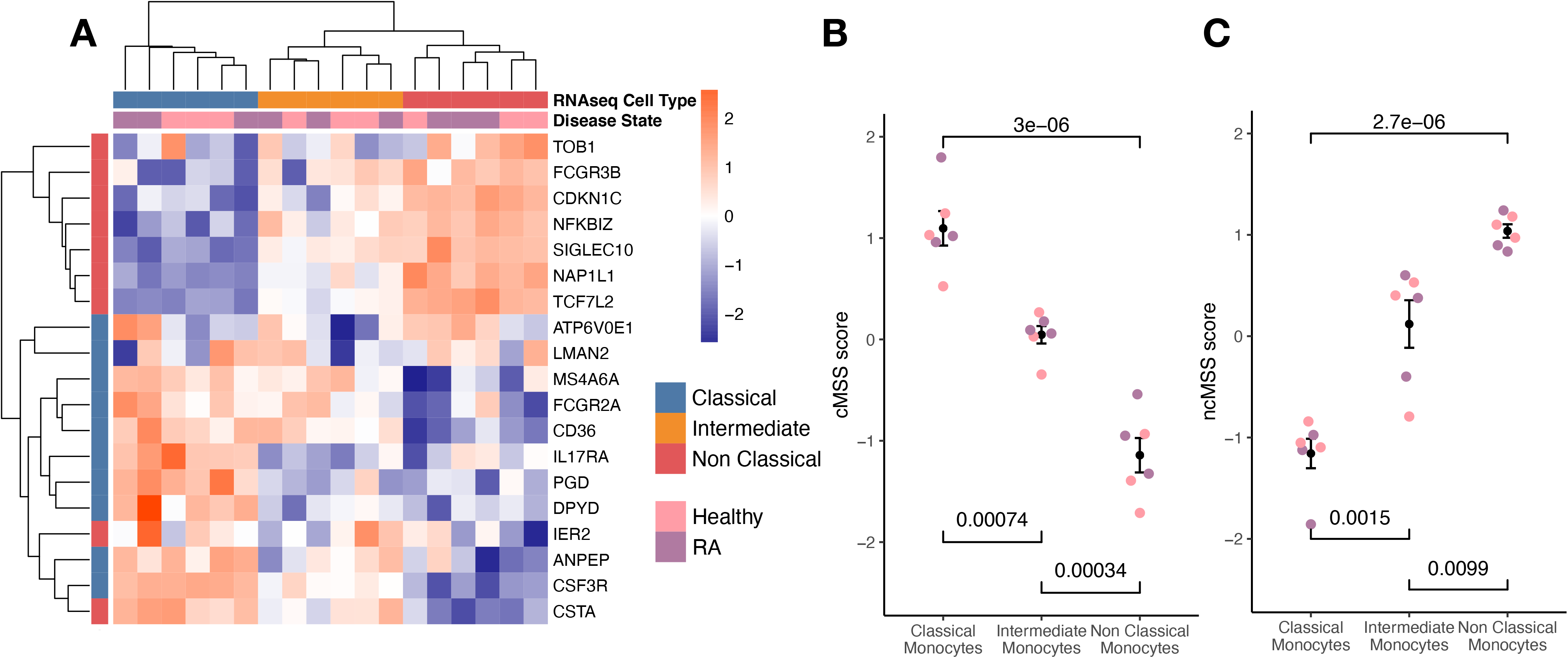
Subset-specific genes are independent of disease state. (a) Heat-map showing expression of subset signature genes (rows) on sorted human monocytes (columns) from independent RNA-seq validation. Samples are labeled by cell type and disease state. (b) Bee-swarm plots displaying classical monocyte subset signature (cMSS) scores across monocyte subsets and disease condition. Significance was computed by t-test. (c) Same as (b) for non-classical monocyte subset signature (ncMSS) scores.

### Monocyte signatures are highly specific across all immune cells

Although our transcriptional signatures are accurate and specific within monocyte subsets, irrespective of disease state and gene expression platform, they were obtained using gene expression data solely derived from monocytes. Therefore, we asked whether our signatures maintained their specificity and accuracy when compared across all immune cell lineages, including B cells, T cells, NK cells. This is important, as their direct application to blood or biopsy-derived expression profiles, which contain multiple and diverse cell populations, would otherwise produce confounded results (38). To answer this question, we compared effect sizes for each gene in our monocyte subset signatures across 20 sorted human immune cell types using 6160 transcriptomes from across 42 different microarray platforms described before (31). Hierarchical clustering of Hedge’s *g* effect sizes (see Methods) of the genes in monocyte subset-specific signatures distinguished myeloid and lymphoid lineages (**Supplementary Figure 3**). Further, within the myeloid cluster, both CD14+ and CD16+ subsets clustered separately from other myeloid cells (**Supplementary Figure 3**).

Next, we calculated cMSS and ncMSS scores for each sample and evaluated their ability to accurately distinguish each subset among all immune cells. As expected, cMSS scores were significantly higher in classical monocytes compared to nonclassical monocytes (t-test p=1.2e-7) and other immune cell types (t-test p<2.2e-16, **Figure 4A**). Similarly, ncMSS were significantly higher in nonclassical monocytes compared to classical monocytes (t-test p=4.6e-9) and other immune cell types (t-test p<2.2e-16, **Figure 4B**). Furthermore, we observed high classification accuracy across both signatures (cMSS AUROC = 0.88; 95% CI: 0.87-0.89; ncMSS AUROC = 0.87; 95% CI: 0.83-0.91; **Supplementary Figure 4**), indicative of their specificity across all immune cells.

**Figure 4:**
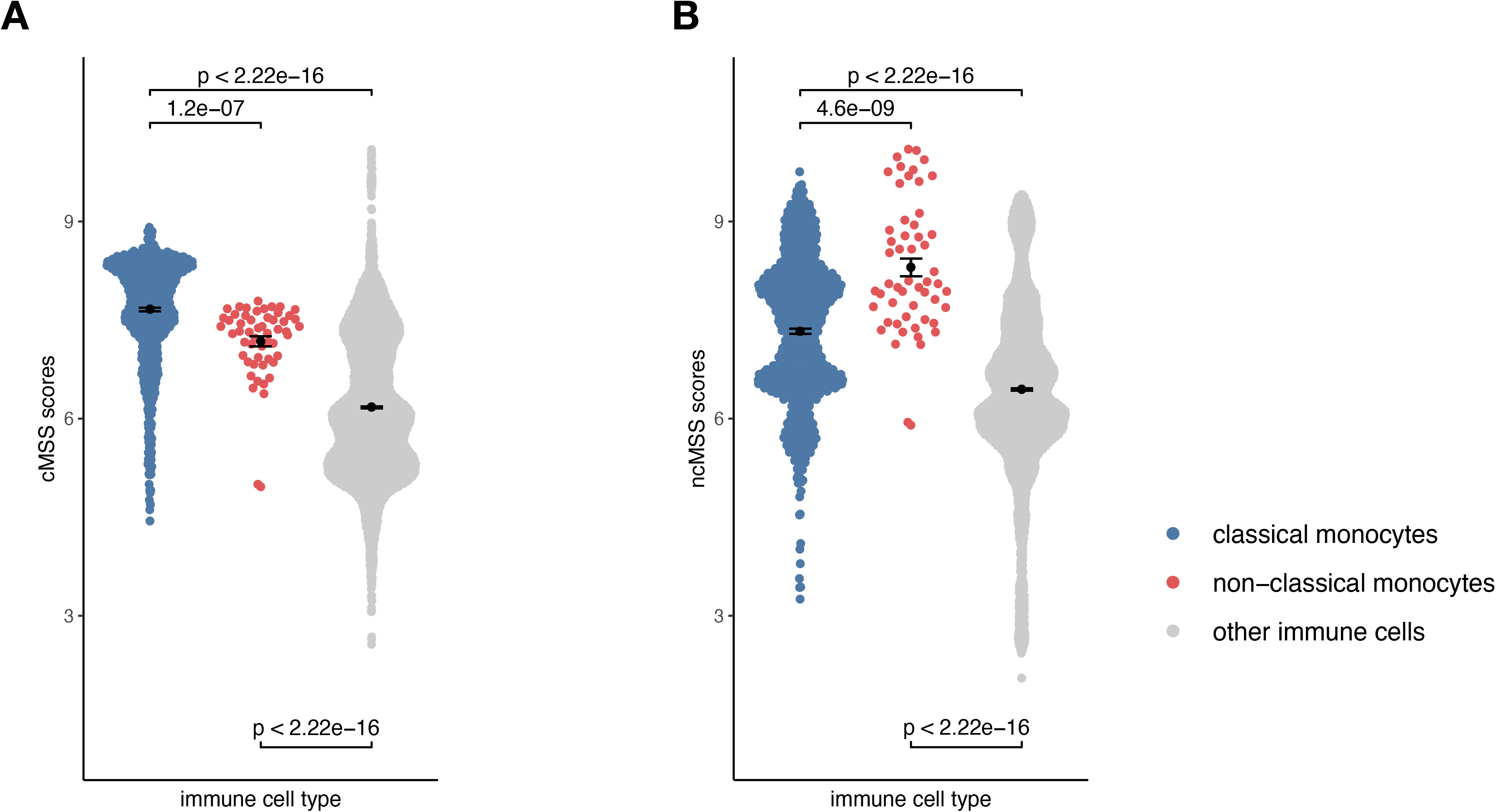
Monocyte signatures are specific across all immune cells. (a) Beeswarm plots displaying cMSS scores across 6160 transcriptomes profiling sorted human immune cells. Colors indicate whether a sample is a classical, non-classical monocyte, or any other immune cell. P-values were computed by t-test. (b) Same as (a) with respect to ncMSS.

### Monocyte signatures reveal changes associated with disease and treatment

Next, we hypothesized that monocyte subset-specific signatures and their corresponding scores, cMSS and ncMSS, could be used to monitor changes in proportions of monocyte subset levels associated with disease. To test this hypothesis, we analyzed transcriptome profiles of whole blood samples (GSE93272) from healthy controls (n=43) and patients with RA (n=232) (39). We computed cMSS and ncMSS for each sample. We observed a significant increase in both cMSS (p = 4.2e-4) and ncMSS (p = 4.4e-8) in RA-patients compared to healthy controls, which is in line with increased monocyte proportion in patients with RA that has been previously observed (**Figure 5A**) (40). Finally, we assessed whether our signatures could detect changes in cellular composition induced by treatment. To this end, we analyzed a longitudinal dataset, GSE80060, profiling whole blood samples of patients affected by sJIA before and after treatment with canakinumab, a monoclonal antibody against IL-1 beta. Changes in levels of circulating monocytes in sJIA have been described, with higher levels during active disease (41-42). When comparing pre- and post-treatment samples, we measured a significant decrease in both classical (p = 2.9e-12) and non-classical (p = 1.5e-6) signatures post-treatment irrespective of response (**Figure 5B**). Our results indicate that our signatures can be used to specifically monitor changes in monocyte subsets occurring during disease and treatment.

**Figure 5:**
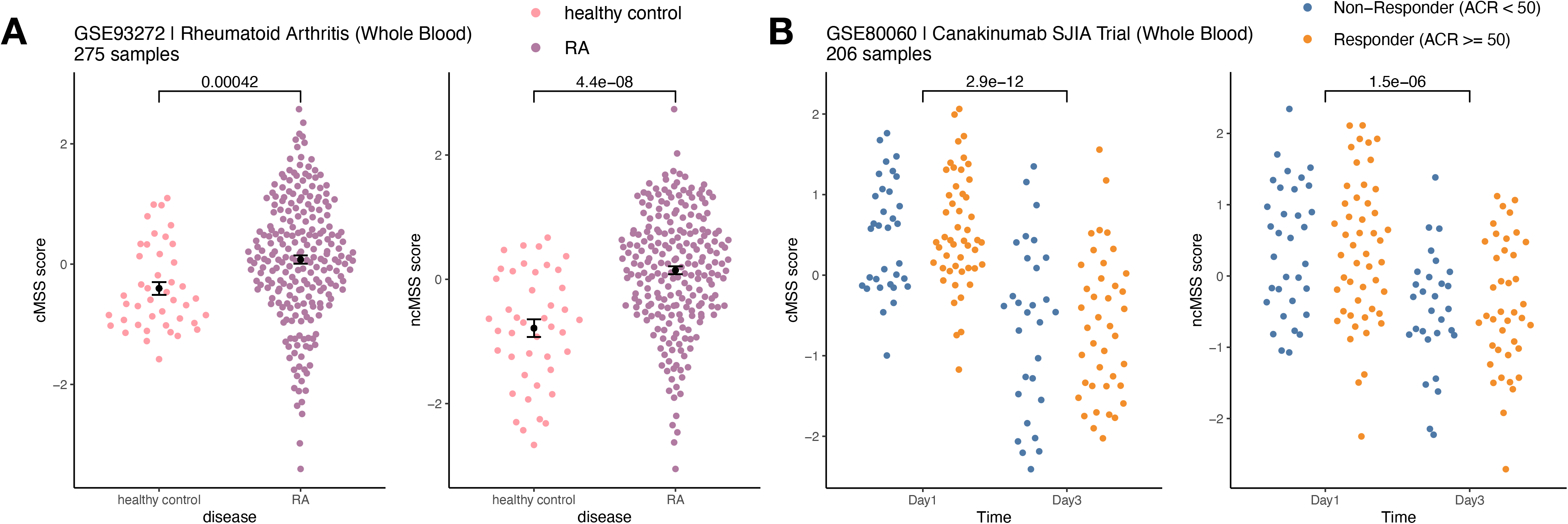
Monocyte subset signatures reveal specific changes in immune cell-composition occurring in disease and treatment. (a) Increase in monocyte levels in Rheumatoid Arthritis (RA) patients compared to healthy controls. Significance measured by Wilcoxon’s Rank Sum Test. (b) Reduction in monocyte levels after Canakinumab treatment of SJIA patients independently of response. Significance measured by Wilcoxon’s Rank Sum Test.

### Validation of monocyte signature genes as novel cell surface markers for subset quantification using cytometry

Cell type-specific gene signatures have been shown to enable accurate *in silico* estimation of corresponding cell types from expression data of mixed-cell samples, such as whole blood or peripheral blood mononuclear cells (PBMCs) (31). We therefore tested whether these signatures could be used to accurately quantify monocyte subsets within human samples by using publicly available expression profiles from healthy human PBMCs with paired flow-cytometry (GSE65316, 43). We indeed found the cMSS to be strongly and significantly correlated with cytometry-measured monocyte proportions across all samples (r = 0.69, p = 6.7e-4; **Figure 6**).

**Figure 6:**
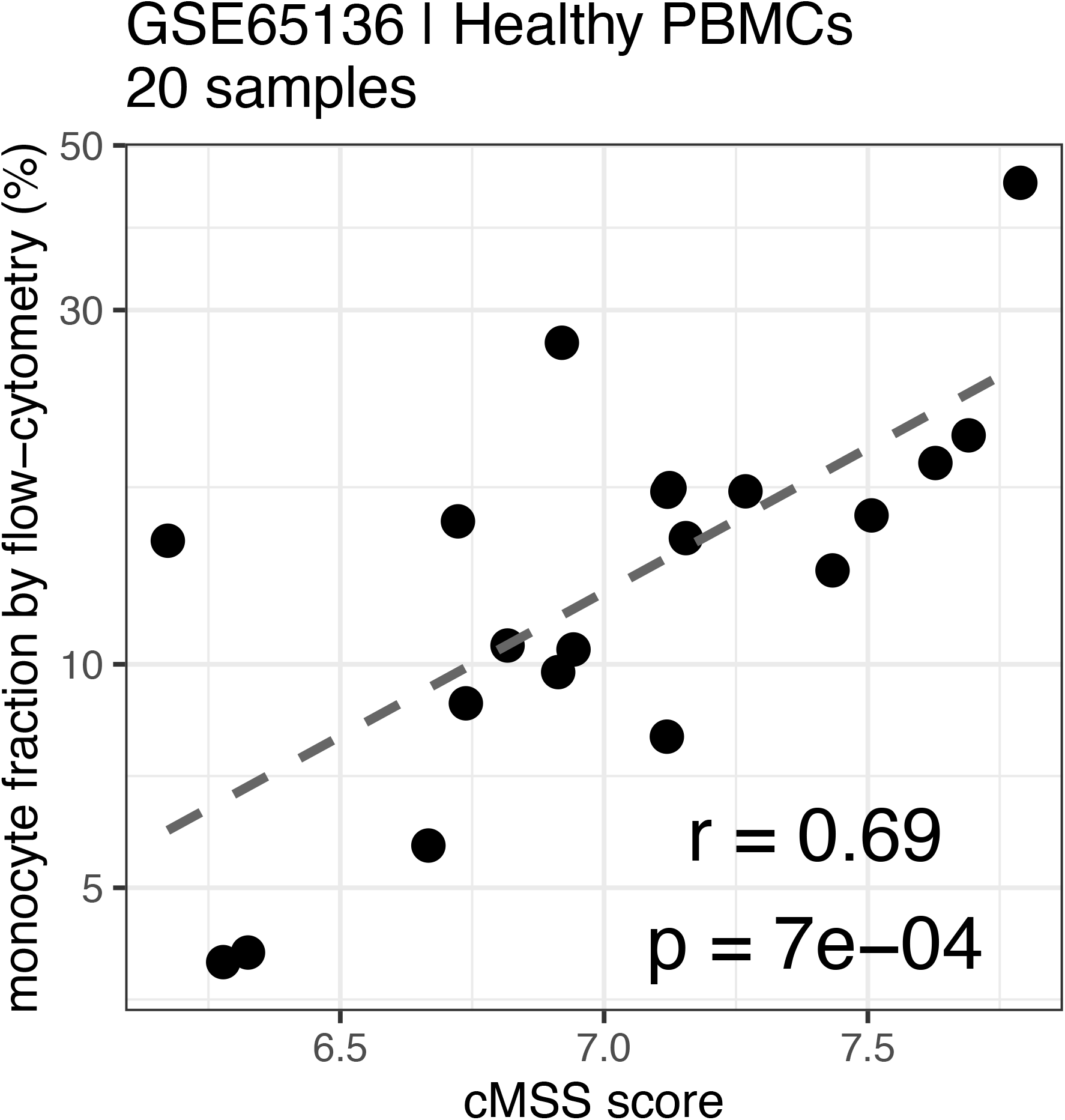
Monocyte signature quantifies monocytes in PBMC by flow-cytometry. Correlation between cMSS score and measured monocyte fraction by flow-cytometry from healthy PBCMs measured in GSE65136. Y-axis indicates measure cellular proportions of the monocyte

Next, we hypothesized that our large-scale transcriptome analysis would enable identification of cell surface markers to improve cellular phenotyping by cytometry using FACS or CyTOF. To test this hypothesis, we selected an extended set of genes that were significantly highly expressed in either classical or non classical monocytes in both discovery and validation samples with an absolute effect size ≥ 1, had documented surface expression, and for which an antibody for follow-up protein quantification by cytometry was commercially available (**Supplementary Table 5**). We selected *CD114* (gene name: *CSF3R*, ES=-1.28, p=2.84e-7), *CD32* (gene name: *FCGR2A*, ES=-1.21, p=9.48e-7), *CD36* (ES=-1.17, p=3.63e-6), and *IL17RA* (ES=-1.10, p=4.39e-6) as markers with higher expression in classical monocytes, and *SIGLEC10* (ES=4.93, p=2.2e-70) as a marker with higher expression in nonclassical monocytes compared to classical monocytes (**Figure 7A,B**).

**Figure 7:**
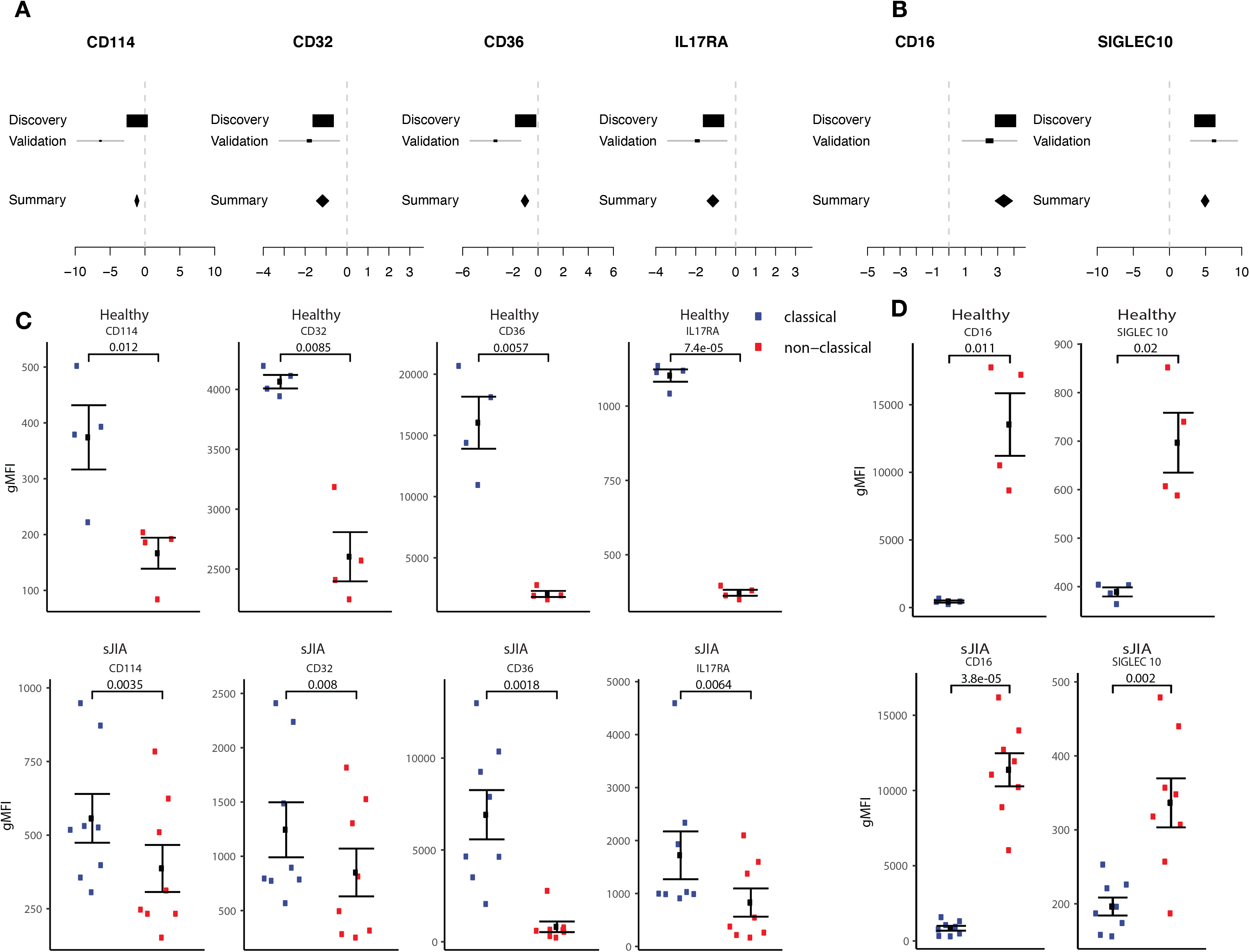
Monocyte signature genes distinguish monocyte subsets by flow-cytometry. (a) mRNA expression effect sizes comparing classical vs. non-classical subsets in both discovery and validation cohorts for each marker associated with classical monocytes (lower effect size values indicate higher expression in classical monocytes). (b) Same as (a) but for markers associated with non-classical monocytes (higher effect size values indicate higher expression in non-classical monocytes). (c) Monocyte subsets were manually gated using FlowJo software. Data shown are geometric mean fluorescence intensity (gmean) in both healthy (top row) and sJIA patients (bottom row). Comparisons made using T-test unpaired unless specified for markers associated with classical monocytes. (d) same as (cs) for markers associated with non-classical monocytes.

We profiled cell surface proteins corresponding to these differentially expressed genes in PBMCs from healthy adult donors (n=8) and pediatric patients with sJIA (n=12). In both healthy adults and sJIA subjects, expression of the selected markers was higher in the corresponding monocyte subset, as predicted by transcriptome analysis (**Figure 7C,D**) and **Supplementary Figure 5**). All of our predicted proteins had significantly different levels between classical and nonclassical monocytes on the cell surface in both healthy controls and sJIA patients (p<0.05).

In summary, we have developed monocyte subset-specific robust and parsimonious gene expression signatures. Our results highlight their specificity and accuracy irrespective of technical and biological confounders and show their utility in translational applications. More importantly, our approach demonstrates that genes differentially expressed between two groups despite biological and technical heterogeneity across multiple independent datasets can be robust differentiators of the two groups at the protein level as well.

## Discussion

Here, we describe the generation and application of robust and parsimonious gene expression signatures to accurately and specifically quantify changes in monocyte subset levels from existing publicly available datasets. By applying a computational framework that integrates existing heterogeneous public expression data from sorted human monocytes, we identified gene signatures for the classical and nonclassical monocyte subsets, each consisting of ten up-regulated genes for the subset of interest. Our analysis presented here builds upon an existing framework that was previously applied to create a new and unbiased basis matrix for cell-mixture deconvolution of gene expression data.

Our current work differs from previous efforts in several meaningful ways: First, our previous work on deconvolution aimed at building a basis matrix, immunoStates, that would account for multiple different immune cell types (n = 20). As a result, immunoStates is composed of over 300 genes, and is applied in its entirety to deconvolve a sample of interest. In contrast, here we focused on creating cell-type specific gene signatures consisting of a small set of genes to be used independent of other cell-specific signatures or any deconvolution framework, while retaining high specificity and accuracy. Second, our current gene selection strategy was chosen to prioritize genes that could be easily used as individual biomarkers for cytometry-based assays. Such strategies take into account the directionality of the markers and their expression difference, to increase the likelihood of validation by qPCR and flow-cytometry. Indeed, a number of genes in our signatures correspond to surface markers with commercially available antibodies. Using this set of markers, we confirmed the subset specificity of the markers in both healthy and disease samples at the protein level. Among the markers identified and validated by flow cytometry, only CD16 and CD36, in addition to CD14 and HLA-DR, have been commonly used to identify monocyte subsets (44-45). The additional markers we identified could thus be potentially useful in further probing the heterogeneity of monocyte subsets, as revealed by recent studies utilizing the high dimensionality of mass cytometry and single cell sequencing to tease out the heterogeneity of the human monocyte population (37-38, 46)

Finally, our analysis leveraged only samples profiled from healthy individuals, whereas our previous work included expression data from disease-affected samples as well. Our rationale for this decision was based on having on average 22 studies per targeted cell type in our discovery set, which triples both the statistical power and the amount of accounted heterogeneity in this study compared to our previous work (8 studies per cell type, 31). We hypothesized these increases would result in more robust signatures, and our validation cohort validated our signatures irrespective of disease state. Analysis of our cohort also revealed that the expression levels of our signature scores in intermediate monocytes were intermediate between the classical and non-classical subsets. It has long been debated whether intermediate monocytes exist as a stable subset or represent a transitional state between classical and non-classical monocytes (47). Another alternative, not necessarily mutually exclusive, is that intermediate monocytes may comprise a heterogeneous cell mix (37, 48). Our results are consistent with evidence that human intermediate monocytes, like similar cells in mice (49) are an intermediate, transitory subset between classical and non-classical monocytes rather than a fixed, independent population (50). To test this hypothesis, we generated an additional expression signature from our discovery data, identifying genes that could specifically distinguish intermediate monocytes compared to all other subsets. Using the same criteria applied for the other signatures, we identified 10 genes that accurately distinguished intermediate monocytes from all other subsets (*ATG2A*, *ATP50, DX39A, EVL, GPR183, LPCAT1, POU2F2, TSC222D4, ZNF14, ARHGAP27*, **Supplementary Figure 6**). Unlike our previous signatures, we did not observe a good distinction of intermediate monocyte samples, neither in our own discovery set, nor in our validation cohort. Similarly, although gene expression analysis, both from microarray as well as from single cell RNA-seq analysis, generally support the concept of genetically separate three monocyte subsets, the exact nature of the intermediate subset, and its relationship to other monocyte subsets, could not be fully determined (51).

Our work has several limitations. First, our discovery data sets consist exclusively of microarray datasets, which can limit the number and type of cell-type specific markers that can be discovered compared to sequencing-based transcriptome profiling. This is particularly relevant in the context of intermediate monocytes. For example, HLA-DM has been identified as a strong discriminating marker that separates the intermediate subset from classical and non-classical (52). However, this marker is not usually profiled on microarrays, limiting our discovery potential. Secondly, we identified our signature genes by simply selecting the top 10 from a ranked list. While this approach is simple and intuitive, it prevents the consideration of other high-ranking genes as potential biomarkers. This potential can be explored in future work, where additional gene set selection strategies can be applied to this data.

Finally, to increase robustness and power of our signatures, our work leverages solely transcriptomic data without accounting for differences occurring post-transcriptionally that may affect final protein levels (53). This concern is especially relevant when translating our results into cytometry/staining based assays that leverage protein expression of surface markers. To this date, high-throughput proteomics data is limited by technical constraints on the number of protein markers that can be simultaneously profiled on a single sample. Advances in mass-cytometry based techniques can in principle extend our ability to profile multiple markers expressed in a single-cell (54), as well as proteomics (55), but at scales substantially lower than transcriptomics-based assays.

In conclusion, we present a collection of robust and parsimonious gene expression signatures that can distinguish and quantify monocyte subsets across disease affected samples and can be used to identify cytometry biomarkers. Our work provides several applications and highlights the potential for our signatures and markers to be used in clinical and translational settings.

## Methods

### Public data collection, annotation, and analysis

Unless otherwise noted, we obtained all gene expression data used in this study from the Gene Expression Omnibus (GEO) database (www.ncbi.nlm.nih.gov/geo/) using the MetaIntegrator R package from CRAN (36) (Supplementary Table 1). All data was manually annotated using the available expression metadata. We normalized each expression dataset using quantile normalization and computed gene-level expression from probelevel data using the original probe annotation files available from GEO as described previously (31). We performed co-normalization, effect size calculation, and gene ranking as previously described (31). We performed gene set selection to identify parsimonious gene signatures using the following criteria: (a) we ranked genes based on effect size (b) we filtered genes that were up-regulated in the cell subset of interest (c) we filtered genes with a mean expression difference of 32 expression units or above (d) We selected the top 10 genes for each subset. We chose these criteria to increase the likelihood of successful independent experimental validation of each marker gene. Signature scores were computed by calculating the geometric mean of expression levels of the signature genes in the dataset of interest, as described previously (17-23). All follow-up analyses were performed using R (v. 3.4.1). Analysis scripts are included as Supplementary Materials.

### Pathway analysis

We performed gene set enrichment analysis (GSEA) as previously described (56). Briefly, for each monocyte subset we computed an effect size vector across all genes as described above. We then applied GSEA to the effect size vectors by comparing to MSigDB, a collection of molecular signatures derived from pathway analysis databases and published molecular data (http://software.broadinstitute.org/gsea/msigdb/index.jsp). We corrected for multiple hypothesis testing across all pathways using Benjamini-Hochberg’s FDR correction. We performed this analysis using the ‘fgsea’ and ‘MSigDB’ packages in R.

### Samples

For the monocyte sorting and expression profiling, de-identified blood samples from 3 healthy adult donors were obtained from the Stanford Blood Center (SBC) and from 3 Rheumatoid Arthritis (RA) patients at UCSF (Supplementary Table 6). For the flow cytometry, 8 de-identified blood samples from healthy adults from the SBC, and 12 samples from patients with Systemic Juvenile Idiopathic Arthritis from Lucille Packard Children’s Hospital were tested (Supplementary Table 7). The work was conducted with approval from the Administrative Panels on Human Subjects Research from Stanford University and UCSF.

### Fluorescent-activated cell sorting of monocytes

Venous blood from healthy controls and RA patients was collected in heparin tubes (BD Vacutainer, BD, Franklin Lakes, NJ); peripheral blood mononuclear cells (PBMCs) were isolated by density gradient centrifugation using LSM Lymphocyte Separation Medium (MP Biomedicals, Santa Ana, CA). PBMCs were enriched for total monocytes using the Pan Monocyte Isolation Kit (Miltenyi Biotech, San Diego, CA). Enriched monocytes were stained for surface antigens as previously described (52). Briefly, cells were stained with LIVE/DEAD Fixable Aqua Dead Cell Stain (Life Technologies, Eugene, OR). Antibodies against CD3, CD19, CD56, and CD66b, all labeled with PercpCy5.5, were used to exclude T cells, B cells, NK cells, and neutrophils respectively (‘dump’); antibodies against CD1c and CD141 were used to identify and exclude dendritic cells. Antibodies against HLA-DR-APC Cy7 (clone L243), CD14-Pacific Blue (clone M5E2), and CD16-PE Cy7 (clone 3G8) were used to identify monocytes and their three subsets. All antibodies are from Biolegend (San Diego, CA). Fluorescence minus one (FMO) were used as control for gating cell populations. Sterile flow cytometry sorting was performed using a BD FACSAria II (BD Biosciences) at the Stanford Shared FACS Facility (SSFF) using a 100uM nozzle, yielding monocyte subset purity of over 98% (verified using the classical subset). Sorted cells were collected into polypropylene tubes containing RPMI media (RPMI+10% Heat inactivated Fetal Bovine Serum +1% Penicillin Streptomycin), counted and spun down for total RNA extraction.

### Total RNA extraction and RNA seq

Total RNA extraction was performed using the Qiagen RNeasy Micro kit (Qiagen, Germantown, MD). Total RNA concentration and quality were determined using a NanoDrop 1000 Spectrophotometer (ThermoFisher Scientific, Waltham, MA) and a BioAnalyzer 2100 (Agilent Technologies, Santa Clara, CA) at the Stanford PAN Facility. RNA sequencing (RNA-seq) was performed by BGI Americas (Cambridge, MA). The total RNA was enriched for mRNA using oligo(dT), and the RNA was fragmented. cDNA synthesis was performed using random hexamers as primers. After purification and end reparation, the short fragments were connected with adapters. The suitable fragments were selected for PCR amplification. The library was then sequenced using Ilumina HiSeq 2000.

### Gene expression by real time RT-PCR

Total RNA from sorted monocytes was converted into cDNA using the iScript Reverse Transcription Supermix (Bio-Rad, Hercules, CA), and cDNA was amplified with SsoAdvanced™ SYBR® Green Supermix (Bio-Rad); reactions were performed in a CFX384 real time PCR instrument (Bio-Rad). Primers sequences were obtained from qPrimerDepot (http://primerdepot.nci.nih.gov/), synthesized at the Stanford University Protein and Nucleic Acid Facility (PAN) and validated in our laboratory as previously described (57) using unsorted total monocytes. Primers are listed on Supplementary Table 8. Relative starting amounts of each gene of interest were determined using the delta delta Cq method.

### Flow cytometry

PBMCs were isolated by density gradient centrifugation; healthy donors cells were treated with ACK lysing buffer (Thermo Fisher) to lyse red blood cells (RBCs). Cells were frozen in 10% DMSO/10% heat inactivated (HI) human AB serum (Corning) for later flow cytometry staining. Frozen PBMC were thawed in RPMI 1640 with 10% HI AB human serum. The cells were washed with DPBS before staining with LIVE/DEAD Aqua (1:1000 dilution in PBS, ThermoFisher Scientific) for 10 min at room temperature. Cells were then washed with flow buffer (DPBS supplemented with 1% BSA and 0.02% NaN3) before blocking for nonspecific binding with flow buffer containing 5% heat inactivated AB serum (Corning), 5% goat serum (ThermoFisher Scientific), 0.5% mouse serum, and 0.5% rat serum for 15 min on ice. After blocking, cells were stained on ice for 30 min with the following fluorochrome-conjugated antibodies in a 12-color staining combination: APC anti-IL17RA (clone W15177A, BIolegend) at 1:50; APC-Vio770anti-CD32 (clone 2E1, Miltenyi Biotec) at 1:100;FITC anti-CD16 (clone 3G8, Biolegend) at 1:20; PerCP/Cy5.5 anti-CD114 (cloneLMM741, Biolegend) at 1: 20; Brilliant Violet 605 anti-Siglec 10 (clone 5G6, BD) at 1:20; Brilliant Violet 785 anti-CD14 (clone M5E2, Biolegend)at 1:50; PE anti-GPR183 (clone SA313E4, Biolegend)at 1:20; PE-CF594 anti-CD13 (clone VM15, BD)at 1: 20; PE/Cy7 anti-HLA-DR (clone L243, Biolegend) at 1:50; BUV395 anti-CD36 (clone CB38, BD) at 1:50. Alexa 700 dump channel includes anti-CD3 (clone UCHT1, Biolegend) at 1:300, anti-CD19 (clone HIB19, Biolegend) at 1:300, anti-CD56 (clone HCD56, Biolegend)at 1:300, anti-CD1c (clone L161, Biolegend)at 1:150, anti-CD66b (clone G10F5, Biolegend) at 1:150, anti-CD141 (clone 501733, Novus Biologicals) at 1:150. After wash, cells were fixed with 200 μl fixation buffer (Cytofix, BD Biosciences) followed by 2 washes with flow buffer and resuspensionin 200 μl flow buffer. Cells were analyzed on a Becton Dickinson LSRII Analyzer at Stanford Shared FACS facility. Data were analyzed using FlowJo version 10 (FlowJo LLC).

## Supporting information

code for analyses and figures

---

**Supplementary Figure 1:**
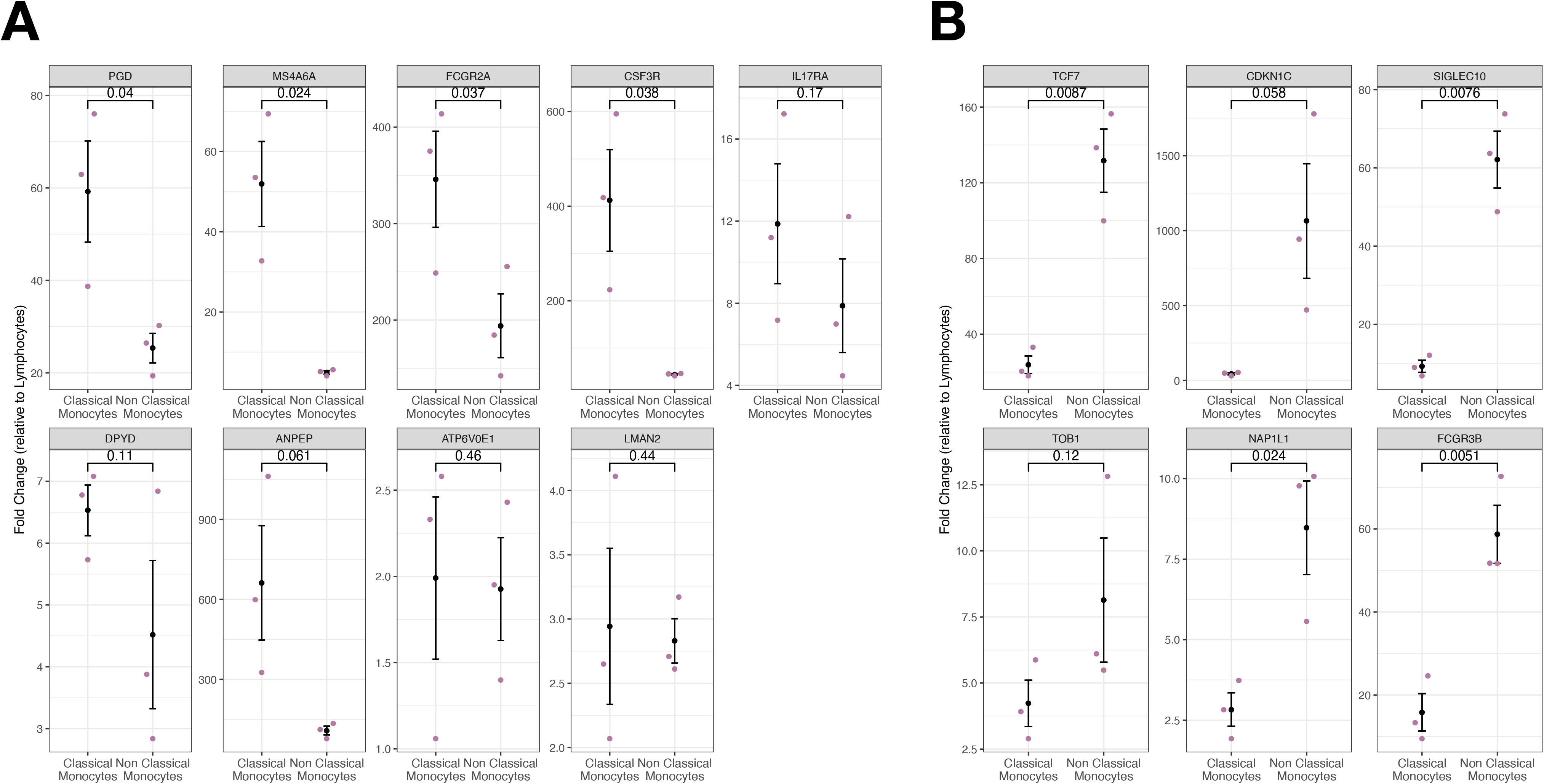
Independent qPCR validation of subset-specific signatures. (a) Dotplots representing classical signature genes fold change measured by qPCR on sorted classical and non classical monocytes from healthy human samples. Significance was estimated by t-test (b) Same as in (a) but for non classical signature genes.

**Supplementary Figure 2:**
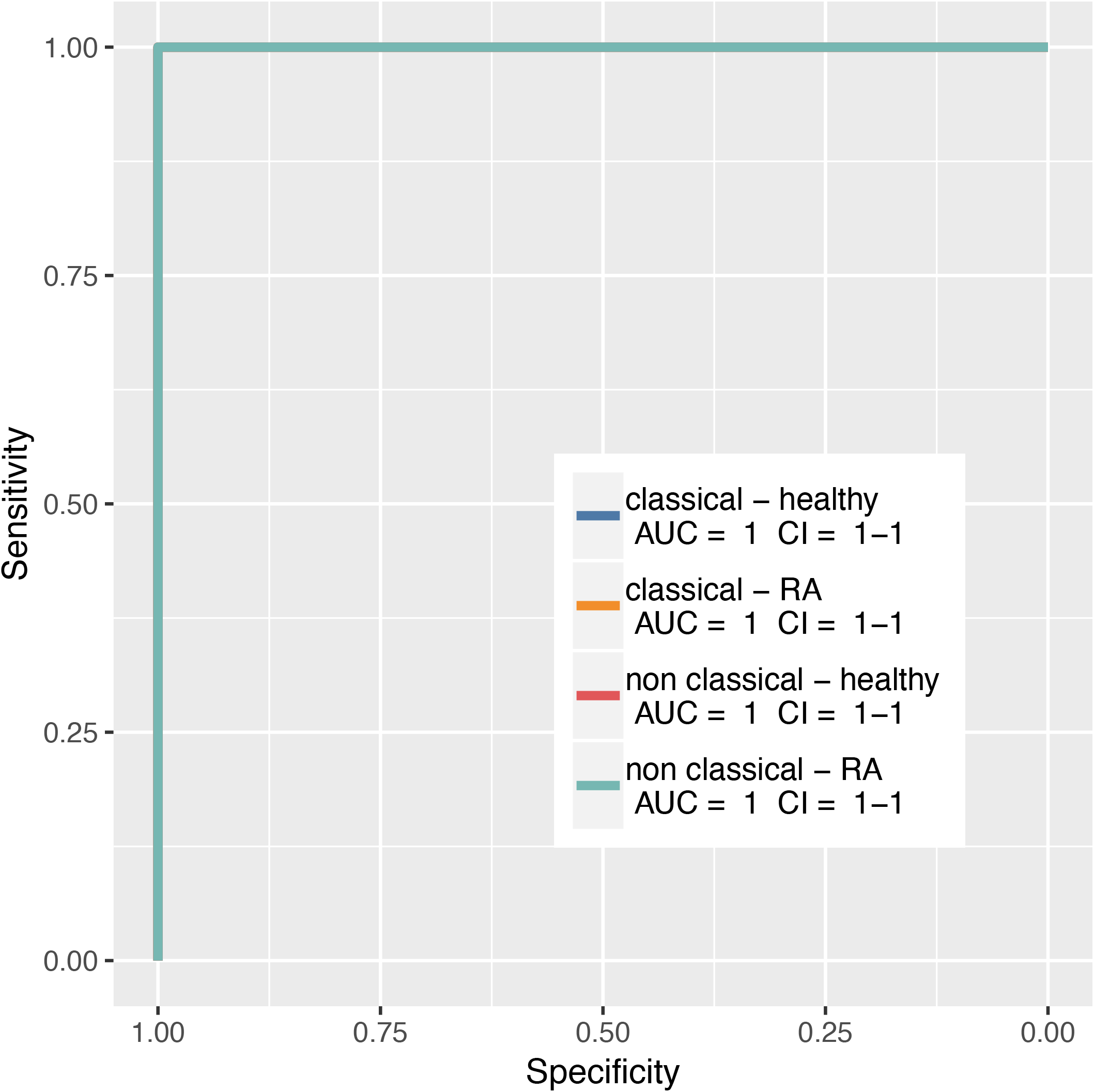
Accuracy in subset classification on independent validation cohort. ROC curves depicting accuracy of subset classification by corresponding gene expression signature across healthy and RA-affected samples.

**Supplementary Figure 3:**
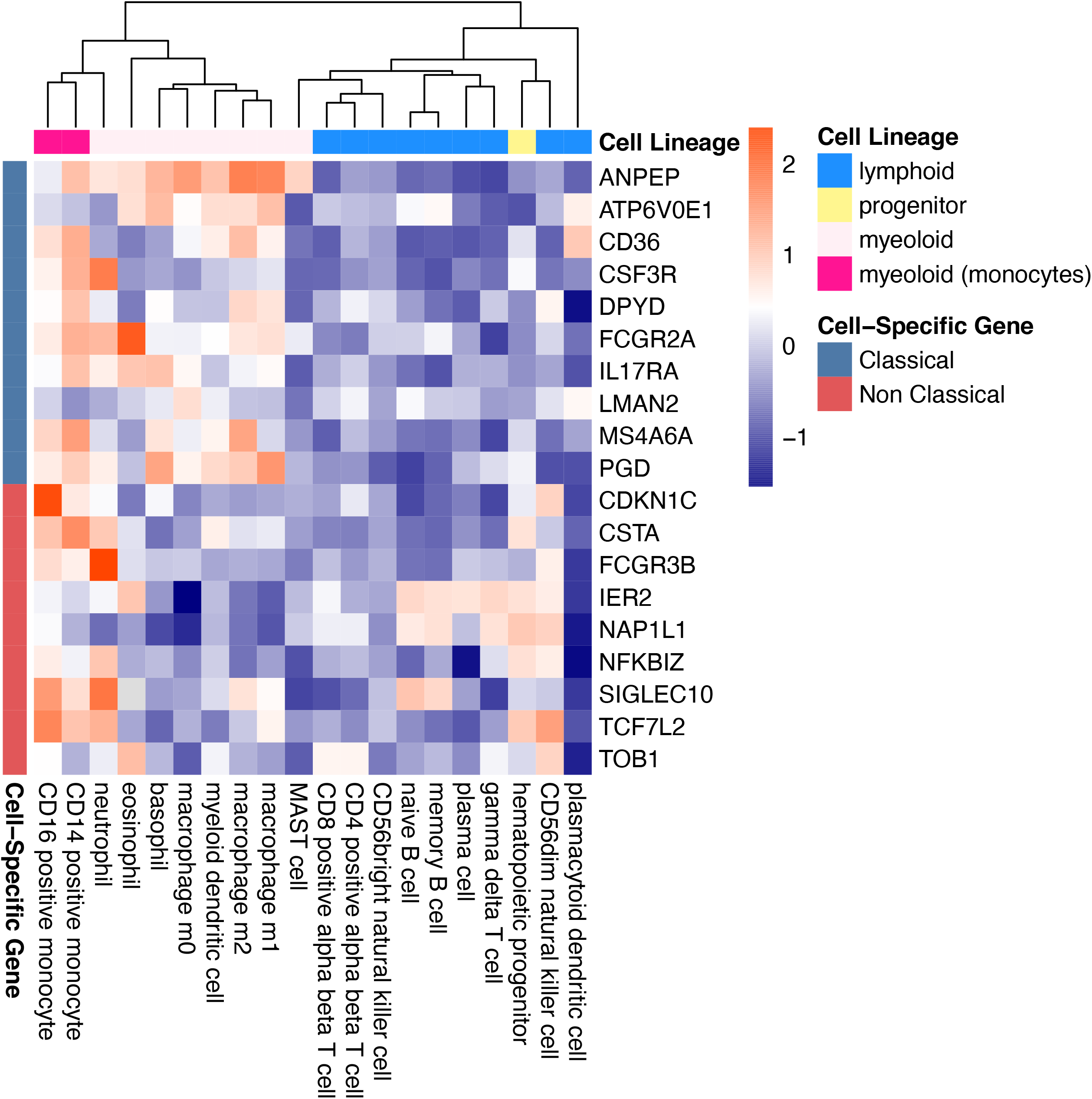
Monocyte signatures are specific across all immune cells. Heat-map displaying gene expression effect sizes of monocyte subset-specific signature genes across 20 human immune cells.

**Supplementary Figure 4:**
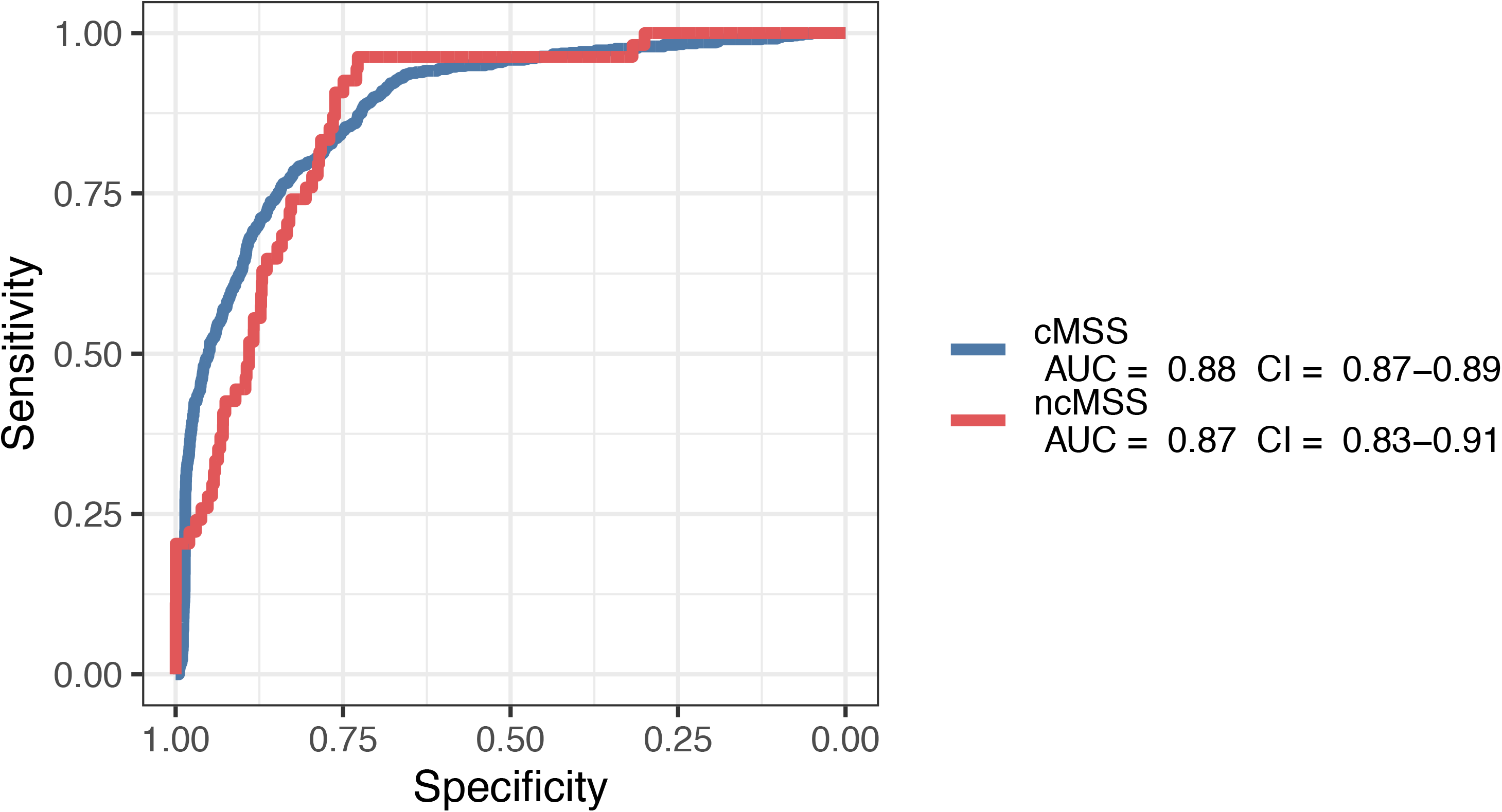
Monocyte signatures are specific across all immune cells. ROC curves depicting accuracy of subset classification by corresponding gene expression signature across 6160 transcriptomes profiling sorted human immune cells

**Supplementary Figure 5:**
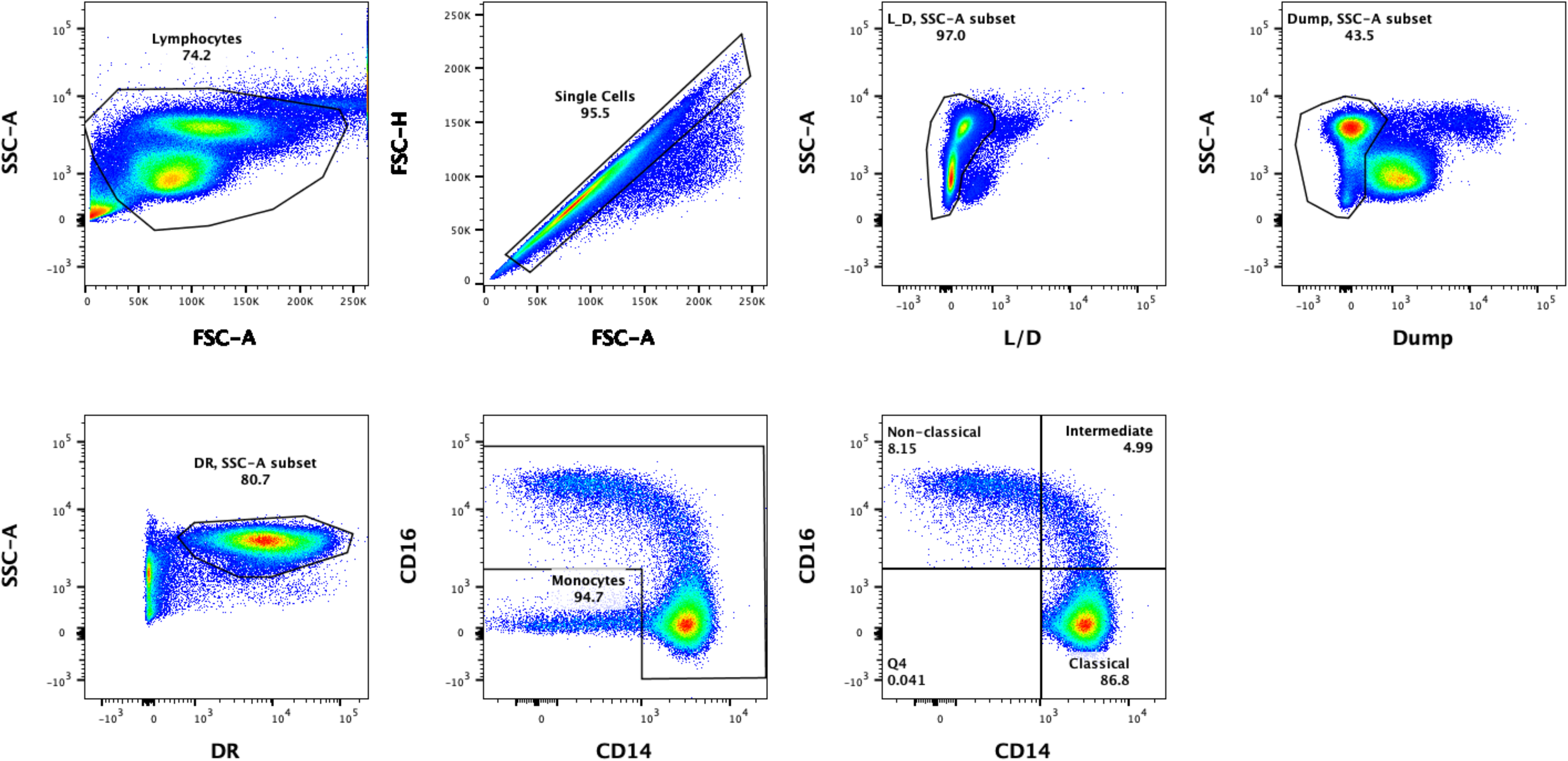
Flow cytometry gating strategy to identify monocyte subsets in peripheral mononuclear cells (PBMCs) PBMCs were gated on a side scatter (SSC) v. forward scatter (FSC) plot, followed by singlet gating based on FSC-H (Height) v. FSC-A (Area). Live cells were determined as negative for LIVE/DEAD (L/D) Aqua staining. T cells (CD3), B cells (CD19), NK cells (CD56), dendritic cells (CD1c and CD141) and neutrophils (CD66b) were excluded as a dump channel. HLA-DR positive cells were selected and plotted on a CD16 vs CD14 plot to identify total monocytes. Gates for classical (CD14+CD16-), intermediate (CD14+CD16+), and nonclassical (CD14-CD16+) subsets are based on Fluorescence minus one (FMO) controls.

**Supplementary Figure 6:**
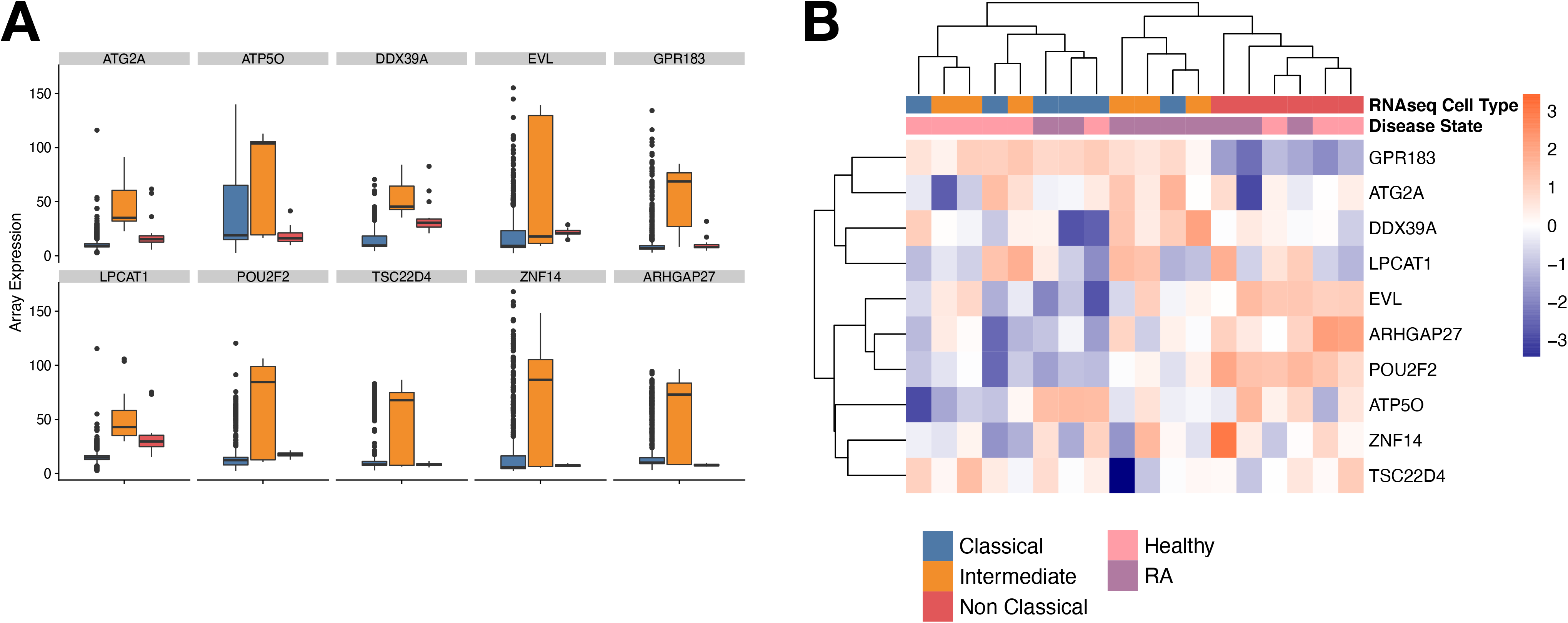
Intermediate monocytes are not distinguishable by our transcriptional signatures. (a) Selection of an intermediate subset gene signature in the discovery cohort. (b) Heat-map displaying the expression of the intermediate subset genes in the RNA-seq validation dataset.

## Notes

### Competing Interest Statement

The authors have declared no competing interest.

