## Supplementary material for "Multicohort Analysis of Publicly-available Monocyte Expression Data Identifies Gene Signatures to Accurately Monitor Subset-specific Changes in Human Diseases": code for analyses and figures: monocyte_paper_figure_3.R

```
#######################################################################################
#Monocyte Signatures Analysis
#by Francesco Vallania, January 30st 2016 @ 9:53pm
#@ Stanford
#~~~~~~~~~~~~~~~~~~~~~~~~~~~~~~~~~~~~~~~~~~~~~~~~~~~~~~~~~~~~~~~~~~~~~~~~~~~~~~~~~~~~~~
#the purpose of this code is generate plots for Figure 3
#######################################################################################

#######################################################################################
#R Functions
#~~~~~~~~~~~~~~~~~~~~~~~~~~~~~~~~~~~~~~~~~~~~~~~~~~~~~~~~~~~~~~~~~~~~~~~~~~~~~~~~~~~~~~
#######################################################################################

#~~~~~~~~~~~~~~~~~~~~~~~~~~~~~~~~~~~~~~~~~~~~~~~~~~~~~~~~~~~~~~~~~~~~~~~~~~~~~~~~~~~~~~
#garbage cleaning
#~~~~~~~~~~~~~~~~~~~~~~~~~~~~~~~~~~~~~~~~~~~~~~~~~~~~~~~~~~~~~~~~~~~~~~~~~~~~~~~~~~~~~~
rm(list=ls())

#~~~~~~~~~~~~~~~~~~~~~~~~~~~~~~~~~~~~~~~~~~~~~~~~~~~~~~~~~~~~~~~~~~~~~~~~~~~~~~~~~~~~~~
#~Load all the necessary libraries
#~~~~~~~~~~~~~~~~~~~~~~~~~~~~~~~~~~~~~~~~~~~~~~~~~~~~~~~~~~~~~~~~~~~~~~~~~~~~~~~~~~~~~~

library(data.table)
library(reshape2)
library(parallel)
library(ggplot2)
library(pROC)
library(BiocGenerics)
library(pheatmap)
library(ggthemes)
library(ggsignif)
library(ggbeeswarm)
library(readxl)
library(DESeq2)


#~~~~~~~~~~~~~~~~~~~~~~~~~~~~~~~~~~~~~~~~~~~~~~~~~~~~~~~~~~~~~~~~~~~~~~~~~~~~~~~~~~~~~~
#rocGGMultiRoc
#~This function computes ROC curves from an input list of pROC objects
#~~~~~~~~~~~~~~~~~~~~~~~~~~~~~~~~~~~~~~~~~~~~~~~~~~~~~~~~~~~~~~~~~~~~~~~~~~~~~~~~~~~~~~
rocGGMultiRoc <- function(validation_roc_list,
                          rocStyle      = 1,
                          rocSize       = 1.4,
                          newLineLegend = TRUE){
  
  #reformat in a data.table
  roc_data_table <- lapply(
    validation_roc_list,
    function(i){
      #
      final_name <- NULL#paste(i$name,"\n",paste("AUC = ",round(i$roc$auc[[1]],2),"%",collapse=""))
      if(newLineLegend == FALSE){
        final_name <- paste(i$name," ",paste("AUC = ",round(i$roc$auc[[1]],2),collapse=""))
      }else{
        final_name <- paste(i$name,"\n",paste("AUC = ",round(i$roc$auc[[1]],2),collapse=""),
                            paste(" CI = ",paste(c(round(ci(i$roc)[1],2),round(ci(i$roc)[3],2)),collapse = "-")))
      }
      
      #
      x <- data.table(name        = final_name,
                      AUC         = paste("AUC = ",round(i$roc$auc[[1]],2),"%",collapse=""),
                      Specificity = i$roc$specificities,
                      Sensitivity = i$roc$sensitivities,
                      FPR         = 100 - i$roc$specificities)
      x <- x[order(Sensitivity),]
      return(x)
    })
  roc_data_table <- rbindlist(roc_data_table)
  
  #generate plot
  p <- ggplot(roc_data_table,aes(x= Specificity,y = Sensitivity,colour = name)) + 
    geom_line(size = rocSize,linetype = rocStyle)                               +
    scale_x_reverse()                                                           + 
    #theme_bw()+
    geom_segment(x=-100, y=0, xend=0, yend=100, linetype=3, colour="grey75")    + 
    theme(legend.key.height=unit(2,"line"),
          legend.title    = element_blank(),
          legend.text     = element_text(size = 10),
          
          legend.position      = c(.45, .11+.11*length(validation_roc_list)), 
          legend.justification = c(0, 1))
  
  #return plot
  return(p)  
}

#~~~~~~~~~~~~~~~~~~~~~~~~~~~~~~~~~~~~~~~~~~~~~~~~~~~~~~~~~~~~~~~~~~~~~~~~~~~~~~~~~~~~~~
#fisher_enrichment_analysis
#~[by Francesco Vallania 05/03/2014]
#~this function computes enrichment analysis of labels in set vs tot
#######################################################################################
fisher_enrichment_analysis <- function(go_set,
                                       go_tot,
                                       unique_names=NULL){
  
  #build a contingency table
  go_tab           <- table(rbind(go_set,go_tot));
  
  #create dataframe
  go_df <- data.frame(label = row.names(go_tab),
                      set   = go_tab[,1],
                      total = go_tab[,2]);
  
  #create dataframe with names if it's available
  if(length(unique_names)>0){
    go_df <- data.frame(label = row.names(go_tab),
                        set   = go_tab[,1],
                        total = go_tab[,2],
                        name  = unique_names[c(match(row.names(go_tab),unique_names[,1])),2]);
  }  
  
  #p-values
  p_values_p <- sapply(1:nrow(go_df),function(i) phyper(go_df[i,2]-1,
                                                        go_df[i,3],
                                                        sum(go_df[-i,3]),
                                                        sum(go_df[,2]),
                                                        lower.tail=F));
  
  #compute bonferroni cut-off
  bonf <- rep(0,nrow(go_df));
  bonf[which(p_values_p<=(0.05/nrow(go_df)))] <- 1;
  
  #return go table with p-values
  go_df <- cbind(go_df,p_values_p,bonf);
  go_df <- go_df[order(go_df$p_values_p),];
  return(go_df);
}

#~~~~~~~~~~~~~~~~~~~~~~~~~~~~~~~~~~~~~~~~~~~~~~~~~~~~~~~~~~~~~~~~~~~~~~~~~~~~~~~~~~~~~~
#go_enrichment_analysis
#~[by Francesco Vallania 05/03/2014]
#~
#######################################################################################
go_enrichment_analysis <- function(gene_set,
                                   tot_gene = NULL,
                                   go_table = NULL){
  #load Hs go_table if it's by default
  if(is.null(go_table)){
    die("input GO table");  
  }
  
  #run code only if there is stuff
  if(length(gene_set)>0){
    #use all genes if it's by default
    if(length(tot_gene)==0){
      tot_gene <- unique(go_table$gene_symbol);
    }
    
    #collect GOs associated to gene set
    go_set       <- data.frame(go=go_table[which(go_table$gene_symbol %in% gene_set),2],set=1);
    go_tot       <- data.frame(go=go_table[which(go_table$gene_symbol %in% tot_gene),2],set=2);
    unique_names <- unique(go_table[,2:3])
    
    #generate and return fisher test table
    return(fisher_enrichment_analysis(go_set,go_tot,unique_names));
  }
  else{
    #if there is no gene return table with every gene with p-value of 1 
    unique_names <- unique(go_table[,2:3]);
    final_df     <- data.frame(label      = unique_names$go_id,
                               set        = 0,
                               total      = 0,
                               name       = unique_names$go_name,
                               p_values_p = 1,
                               bonf       = 0)
    return(final_df);
  }
}


#imported from MetaIntegrator package
getES <- function(v, g){
  ## function to calculate basic statistics for two-class comparison for a single gene
  ## v: vector of samples (one row from an expression matrix)
  ## g: vector of composed of 0,1 values for every sample in v; defines case or control
  ## 
  ## encoding: CASES should be designated with the value 1, and CONTROLS with the value 0 to preserve the 
  ## directionality of the equation (mean of cases - mean of controls)
  
  # check to make sure every sample in expression matrix is assigned case or control value
  if ( length(v) != length(g) ) {
    stop("number of samples in expression matrix is different from length of class vector -- exiting")
  } 
  
  .cleanNA <- function(x) return( x[!is.na(x) & is.finite(x) ] )
  
  x <- .cleanNA( v[ which(g==1) ] )
  y <- .cleanNA( v[ which(g==0) ] )
  
  n1 <- length(x); n2 <- length(y)
  if( n1 < 2 | n2 < 2 )
    return( c(n1=NA, m1=NA, sd1=NA,
              n2=NA, m2=NA, sd2=NA,
              diff=NA, pooled.sd=NA,
              g=NA, se.g=NA) )
  
  #compute mean difference
  m1 <- mean(x)
  m2 <- mean(y)
  diff <- m1 - m2
  
  #compute pooled standard deviation
  sd1  <- stats::sd(x)
  sd2  <- stats::sd(y)
  sp   <- sqrt( ( (n1-1)*sd1^2 + (n2-1)*sd2^2 )/( n1 + n2 - 2 ) )
  
  #correct for small sample size [from Cohen's D to Hedge's G]
  cf   <- 1 - 3/( 4*(n1 + n2) - 9 ) # small sample size correction
  g    <- cf * diff/sp
  se.g <- sqrt( (n1+n2)/(n1*n2) + 0.5*g^2 /(n1+n2-3.94) )
  
  return( c(n1=n1, m1=m1, sd1=sd1,
            n2=n2, m2=m2, sd2=sd2,
            diff=diff, pooled.sd=sp,
            g=g, se.g=se.g) ) 
}

#######################################################################################
#######################################################################################
#~~~~~~~~~~~~~~~~~~~~~~~~~~~~~~~~~~~~~~~~~~~~~~~~~~~~~~~~~~~~~~~~~~~~~~~~~~~~~~~~~~~~~~
#R Main
#~~~~~~~~~~~~~~~~~~~~~~~~~~~~~~~~~~~~~~~~~~~~~~~~~~~~~~~~~~~~~~~~~~~~~~~~~~~~~~~~~~~~~~
#######################################################################################
#######################################################################################

#~~~~~~~~~~~~~~~~~~~~~~~~~~~~~~~~~~~~~~~~~~~~~~~~~~~~~~~~~~~~~~~~~~~~~~~~~~~~~~~~~~~~~~
#~~~~~~~~~~~~~~~~~~~~~~~~~~~~~~~~~~~~~~~~~~~~~~~~~~~~~~~~~~~~~~~~~~~~~~~~~~~~~~~~~~~~~~
#RNA seq analysis
#~~~~~~~~~~~~~~~~~~~~~~~~~~~~~~~~~~~~~~~~~~~~~~~~~~~~~~~~~~~~~~~~~~~~~~~~~~~~~~~~~~~~~~
#~~~~~~~~~~~~~~~~~~~~~~~~~~~~~~~~~~~~~~~~~~~~~~~~~~~~~~~~~~~~~~~~~~~~~~~~~~~~~~~~~~~~~~

#~~~~~~~~~~~~~~~~~~~~~~~~~~~~~~~~~~~~~~~~~~~~~~~~~~~~~~~~~~~~~~~~~~~~~~~~~~~~~~~~~~~~~~
#
#~~~~~~~~~~~~~~~~~~~~~~~~~~~~~~~~~~~~~~~~~~~~~~~~~~~~~~~~~~~~~~~~~~~~~~~~~~~~~~~~~~~~~~

#get data
sampleAllRNAseq <- fread('expression_data/rna_seq_all_genes.csv')

#get pheno
phenoDT <- data.table('sample'=paste("EM",1:18,sep=''),'disease'='RA','celltype'='CD14+ monocytes')

phenoDT[grep("EM7|EM8|EM9|10|11|12|13|14|15",sample),disease:='healthy']
phenoDT[grep("EM2|EM5|EM8|11|14|17",sample),celltype:='CD16+ monocytes']
phenoDT[grep("EM3|EM6|EM9|12|15|18",sample),celltype:='intermediate monocytes']

#
expMat <- as.matrix(sampleAllRNAseq[,grep("EM",colnames(sampleAllRNAseq)),with=FALSE])
row.names(expMat) <- sampleAllRNAseq$Symbol

#log transform [use pseudocount]
logExpMat <- log2((expMat+0.01))
row.names(logExpMat) <- sampleAllRNAseq$Symbol
colnames(logExpMat)  <- gsub("_FPKM","",colnames(expMat))

#compute effect sizes
esComp <- function(inputMat,case = NULL){
  
  rnaSeqDTmolten <- as.data.table(melt(inputMat))
  setnames(rnaSeqDTmolten,c("gene",'sample','value'))
  
  rnaSeqDTmolten[,class:=0]
  rnaSeqDTmolten[sample %in% case,class:=1]
  
  esDT <- rnaSeqDTmolten[,getES(value,class),gene]
  namesO <- c("n1","m1","sd1",
              "n2","m2","sd2",
              "diff","pooled.sd",
              "g","se.g")
  esDT <- dcast.data.table(esDT[,names:=rep(namesO,esDT[,.N]/length(namesO))],gene~names,value.var='V1')
  
  esDT[,p.value:=2*(1-pnorm(abs(g/se.g)))]
  esDT[,fdr:=p.adjust(p.value,method = 'fdr')]
  
  esDT <- esDT[order(p.value)]
  
  return(esDT)
}

esCD14 <- esComp(logExpMat,case = phenoDT[grep("CD14",        celltype)]$sample)
esCD16 <- esComp(logExpMat,case = phenoDT[grep("CD16",        celltype)]$sample)
esINT  <- esComp(logExpMat,case = phenoDT[grep("intermediate",celltype)]$sample)


geneSetFDR5p <- rbind(esCD14[fdr<0.05],
                      esCD16[fdr<0.05],
                      esINT[fdr<0.05])$gene

#~~~~~~~~~~~~~~~~~~~~~~~~~~~~~~~~~~~~~~~~~~~~~~~~~~~~~~~~~~~~~~~~~~~~~~~~~~~~~~~~~~~~~~
#Plot heatmap of differentially expressed genes
#~~~~~~~~~~~~~~~~~~~~~~~~~~~~~~~~~~~~~~~~~~~~~~~~~~~~~~~~~~~~~~~~~~~~~~~~~~~~~~~~~~~~~~

#create annotation
colorDisDf <- as.data.frame(phenoDT[,c(2:3),with=F])
row.names(colorDisDf) <- phenoDT$sample
colnames(colorDisDf)  <- c("Disease State","RNAseq Sample")


#~~~~~~~~~~~~~~~~~~~~~~~~~~~~~~~~~~~~~~~~~~~~~~~~~~~~~~~~~~~~~~~~~~~~~~~~~~~~~~~~~~~~~~
#Compare with genes from Lyron
#~~~~~~~~~~~~~~~~~~~~~~~~~~~~~~~~~~~~~~~~~~~~~~~~~~~~~~~~~~~~~~~~~~~~~~~~~~~~~~~~~~~~~~

#combine effect sizes from RNA-seq data
esCD14[,cellType:='Classical']
esCD16[,cellType:='Non_Classical']
esINT[, cellType:='Intermediate']

rnaSeqCombined <- rbind(esCD14,esCD16,esINT)

#~~~~~~~~
geneListMonoSubs <- readRDS('expression_data/monocyte_subset_final_gene_list.rds')


colorDT <- unique(phenoDT[,c(2,3),with=F])
colorV  <- colorDT$color
names(colorV) <- colorDT$group

# list(disease=c("midnightblue","brickred1"),celltype = c(""))
#create annotation
geneDisDf <- as.data.frame(geneListMonoSubs[,c(2),with=F])
row.names(geneDisDf) <- geneListMonoSubs$gene
colnames(geneDisDf) <- 'Array Target Cell'

#~~~~~~~~~~~~~~~~~~~~~~~~~~~~~~~~~~~~~~~~~~~~~~~~~~~~~~~~~~~~~~~~~~~~~~~~~~~~~~~~~~~~~~
#
#~~~~~~~~~~~~~~~~~~~~~~~~~~~~~~~~~~~~~~~~~~~~~~~~~~~~~~~~~~~~~~~~~~~~~~~~~~~~~~~~~~~~~~

#~~~~~~~~~~
colorDisDf <- as.data.frame(phenoDT[,c(2:3),with=F])
row.names(colorDisDf) <- phenoDT$sample
colnames(colorDisDf)  <- c("Disease State","RNAseq Cell Type")

colorDisDf[,2] <- gsub("CD14\\+ monocytes","Classical",colorDisDf[,2])
colorDisDf[,2] <- gsub("CD16\\+ monocytes","Non Classical",colorDisDf[,2])
colorDisDf[,2] <- gsub("intermediate monocytes","Intermediate",colorDisDf[,2])

colorDisDf[,1] <- gsub("healthy","Healthy",colorDisDf[,1])


geneDisDf <- as.data.frame(geneListMonoSubs[,gsub("_"," ",cellType)])
row.names(geneDisDf) <- geneListMonoSubs$gene
colnames(geneDisDf) <- 'Cell-Specific Gene'

#fix this
annotation_colors = list("RNAseq Cell Type"    = c("Classical"            = "#4E79A7",
                                                   "Intermediate"         = "#F28E2B",
                                                   "Non Classical"        = "#E15759"),
                         "Disease State"       = c("Healthy" = '#FF9DA7',
                                                   "RA"      = '#B07AA1'),
                         'Cell-Specific Gene' = c('Classical'              = "#4E79A7",
                                                  'Non Classical'          = "#E15759"))

pdf('pdfs/20201201_figure3_A.pdf',
    height=5*1.3,width=6*1.3)
pheatmap(logExpMat[row.names(logExpMat) %in% geneListMonoSubs[cellType != 'Intermediate']$gene,],scale='row',
         show_rownames = T,
         show_colnames = F,
         border_color  = NA,
         annotation_col    = colorDisDf,
         annotation_row    = geneDisDf,
         annotation_colors = annotation_colors,
         color=grDevices::colorRampPalette(c("navyblue","white","orangered"))(1000))
dev.off()


pdf('pdfs/20201201_supplementary_figure6.pdf',
    height=4,width=8)
pheatmap(logExpMat[row.names(logExpMat) %in% geneListMonoSubs[cellType == 'Intermediate']$gene,],scale='row',
         show_rownames = T,
         show_colnames = F,
         border_color  = NA,
         annotation_col    = colorDisDf,
         annotation_colors = annotation_colors,
         color=grDevices::colorRampPalette(c("navyblue","white","orangered"))(1000))
dev.off()


#~~~~~~~~~~~~~~~~~~~~~~~~~~~~~~~~~~~~~~~~~~~~~~~~~~~~~~~~~~~~~~~~~~~~~~~~~~~~~~~~~~~~~~
#Compute classification accuracy of signature on RNA-seq dataset
#~~~~~~~~~~~~~~~~~~~~~~~~~~~~~~~~~~~~~~~~~~~~~~~~~~~~~~~~~~~~~~~~~~~~~~~~~~~~~~~~~~~~~~

#
#~~~~~~~~~~~~~~~~~~~~~~~~~~~~~~~~~~~~~~~~~~~~~~~~~~~~~~~~~~~~~~~~~~~~~~~~~~~~~~~~~~~~~~
rnaSeqValid <- as.data.table(melt(logExpMat[row.names(logExpMat) %in% geneListMonoSubs[cellType != 'Intermediate']$gene,]))

rnaSeqValid[,cellType:='Classical\nMonocytes']
rnaSeqValid[grep("EM2|EM5|EM8|EM11|EM14|EM17",Var2),cellType:='Non Classical\nMonocytes']
rnaSeqValid[grep("EM3|EM6|EM9|EM12|EM15|EM18",Var2),cellType:='Intermediate\nMonocytes']

rnaSeqValid[,diseaseState:='RA']
rnaSeqValid[grep("EM7|EM8|EM9|EM10|EM11|EM12|EM13|EM14|EM15",Var2),diseaseState:='healthy']

#
#~~~~~~~~~~~~~~~~~~~~~~~~~~~~~~~~~~~~~~~~~~~~~~~~~~~~~~~~~~~~~~~~~~~~~~~~~~~~~~~~~~~~~~
classSign <- rnaSeqValid[Var1 %in% geneListMonoSubs[cellType=='Classical']$gene,mean(value),by=.(cellType,diseaseState,Var2)]
classSign[,class:=0]
classSign[cellType=='Classical\nMonocytes',class:=1]

NclassSign <- rnaSeqValid[Var1 %in% geneListMonoSubs[cellType=='Non_Classical']$gene,mean(value),by=.(cellType,diseaseState,Var2)]
NclassSign[,class:=0]
NclassSign[cellType=='Non Classical\nMonocytes',class:=1]

IclassSign <- rnaSeqValid[Var1 %in% geneListMonoSubs[cellType=='Intermediate']$gene,mean(value),by=.(cellType,diseaseState,Var2)]
IclassSign[,class:=0]
IclassSign[cellType=='Intermediate\nMonocytes',class:=1]


#violin/beeswarm plots
#~~~~~~~~~~~~~~~~~~~~~~~~~~~~~~~~~~~~~~~~~~~~~~~~~~~~~~~~~~~~~~~~~~~~~~~~~~~~~~~~~~~~~~

#ggplot with standard error -- trying new things out just to visualize
serr <- function(inV){sd(inV)/sqrt(length(inV))}

classSign$cellType  <- factor(classSign$cellType,c('Classical\nMonocytes',
                                                   'Intermediate\nMonocytes',
                                                   'Non Classical\nMonocytes'),
                              ordered = T)
NclassSign$cellType <- factor(NclassSign$cellType,c('Classical\nMonocytes',
                                                    'Intermediate\nMonocytes',
                                                    'Non Classical\nMonocytes'),
                              ordered = T)


#update on 2020/01/29
p <- ggplot(
  rbind(classSign[,V2:=scale(V1)][
    ,mean:=mean(V2),
    .(cellType)][,serr:=serr(V2),
                 .(cellType)][,sign:='Classical Monocyte\nSignature']),
  aes(x=cellType,y=V2)) +
  geom_point(aes(y=mean,x=cellType)) +
  geom_errorbar(aes(ymax=mean+serr,ymin=mean-serr),width=.1) +
  geom_quasirandom(aes(color=gsub("hea","Hea",diseaseState)),width=.2,size=2) +
  scale_color_manual(values=c('#FF9DA7','#B07AA1')) +
  labs(color="")     +
  theme_tufte(base_family='Helvetica') +
  geom_signif(comparisons = list(c('Classical\nMonocytes','Non Classical\nMonocytes'),
                                 c('Intermediate\nMonocytes','Non Classical\nMonocytes'),
                                 c('Classical\nMonocytes','Intermediate\nMonocytes')),
              test = 't.test',
              tip_length = c(.03,-.03,-.03),
              y_position = c(2,-2.5,-2)) +
  theme(axis.line.x = element_line(),
        axis.line.y = element_line()) +
  ylab('cMSS score') + xlab("")

pdf('pdfS/20201201_figure3_B_classical.pdf',
    width=4,height=4.5)
plot(p)
dev.off()

q <- ggplot(
  rbind(
    NclassSign[, V2:=scale(V1)][
      ,mean:=mean(V2),
      .(cellType)][,serr:=serr(V2),
                   .(cellType)][,sign:='Non Classical Monocyte\nSignature']),
  aes(x=cellType,y=V2)) +
  geom_point(aes(y=mean,x=cellType)) +
  geom_errorbar(aes(ymax=mean+serr,ymin=mean-serr),width=.1) +
  geom_quasirandom(aes(color=gsub("hea","Hea",diseaseState)),width=.2,size=2) +
  scale_color_manual(values=c('#FF9DA7','#B07AA1')) +
  labs(color="")     +
  theme_tufte(base_family='Helvetica') +
  geom_signif(comparisons = list(c('Classical\nMonocytes','Non Classical\nMonocytes'),
                                 c('Intermediate\nMonocytes','Non Classical\nMonocytes'),
                                 c('Classical\nMonocytes','Intermediate\nMonocytes')),
              test = 't.test',
              tip_length = c(.03,-.03,-.03),
              y_position = c(2,-2.5,-2)) +
  theme(axis.line.x = element_line(),
        axis.line.y = element_line()) +
  ylab('ncMSS score') + xlab("")

pdf('pdfS/20201201_figure3_C_nonclassical.pdf',
    width=4,height=4.5)
plot(q)
dev.off()

##############

#remake AUC plot
rocCH <- roc(classSign[diseaseState=='healthy']$class,classSign[diseaseState=='healthy']$V1)
rocCR <- roc(classSign[diseaseState=='RA']$class,classSign[diseaseState=='RA']$V1)

rocNR <- roc(NclassSign[diseaseState=='RA']$class,NclassSign[diseaseState=='RA']$V1)
rocNH <- roc(NclassSign[diseaseState=='healthy']$class,NclassSign[diseaseState=='healthy']$V1)


rocC  <- roc(classSign$class,classSign$V1)
rocN  <- roc(NclassSign$class,NclassSign$V1)


pdf('pdfs/20201201_supplementary_figure2.pdf',
    height=5,width=7)
plot(rocGGMultiRoc(list(list(name='classical signature',    roc = rocC),
                        list(name='non-classical signature',roc = rocN))) + 
       theme_bw() + scale_color_manual(values=c('#4E79A7','#E15759')) + labs(color=''))
dev.off()
```
