## Supplementary material for "Multicohort Analysis of Publicly-available Monocyte Expression Data Identifies Gene Signatures to Accurately Monitor Subset-specific Changes in Human Diseases": code for analyses and figures: monocyte_paper_figure_4.R

```
#######################################################################################
#Monocyte Signatures Analysis
#by Francesco Vallania, January 30st 2016 @ 9:53pm
#@ Stanford
#~~~~~~~~~~~~~~~~~~~~~~~~~~~~~~~~~~~~~~~~~~~~~~~~~~~~~~~~~~~~~~~~~~~~~~~~~~~~~~~~~~~~~~
#the purpose of this code is generate plots for Figure 4
#######################################################################################

#######################################################################################
#R Functions
#~~~~~~~~~~~~~~~~~~~~~~~~~~~~~~~~~~~~~~~~~~~~~~~~~~~~~~~~~~~~~~~~~~~~~~~~~~~~~~~~~~~~~~
#######################################################################################

#~~~~~~~~~~~~~~~~~~~~~~~~~~~~~~~~~~~~~~~~~~~~~~~~~~~~~~~~~~~~~~~~~~~~~~~~~~~~~~~~~~~~~~
#garbage cleaning
#~~~~~~~~~~~~~~~~~~~~~~~~~~~~~~~~~~~~~~~~~~~~~~~~~~~~~~~~~~~~~~~~~~~~~~~~~~~~~~~~~~~~~~
rm(list=ls())

#~~~~~~~~~~~~~~~~~~~~~~~~~~~~~~~~~~~~~~~~~~~~~~~~~~~~~~~~~~~~~~~~~~~~~~~~~~~~~~~~~~~~~~
#~Load all the necessary libraries
#~~~~~~~~~~~~~~~~~~~~~~~~~~~~~~~~~~~~~~~~~~~~~~~~~~~~~~~~~~~~~~~~~~~~~~~~~~~~~~~~~~~~~~

library(data.table)
library(reshape2)
library(parallel)
library(ggplot2)
library(pROC)
library(BiocGenerics)
library(pheatmap)
library(ggthemes)
library(ggsignif)
library(ggbeeswarm)
library(readxl)
library(DESeq2)


#~~~~~~~~~~~~~~~~~~~~~~~~~~~~~~~~~~~~~~~~~~~~~~~~~~~~~~~~~~~~~~~~~~~~~~~~~~~~~~~~~~~~~~
#rocGGMultiRoc
#~This function computes ROC curves from an input list of pROC objects
#~~~~~~~~~~~~~~~~~~~~~~~~~~~~~~~~~~~~~~~~~~~~~~~~~~~~~~~~~~~~~~~~~~~~~~~~~~~~~~~~~~~~~~
rocGGMultiRoc <- function(validation_roc_list,
                          rocStyle      = 1,
                          rocSize       = 1.4,
                          newLineLegend = TRUE){
  
  #reformat in a data.table
  roc_data_table <- lapply(
    validation_roc_list,
    function(i){
      final_name <- NULL
      
      if(newLineLegend == FALSE){
        final_name <- paste(i$name," ",paste("AUC = ",round(i$roc$auc[[1]],2),collapse=""))
      }else{
        final_name <- paste(i$name,"\n",paste("AUC = ",round(i$roc$auc[[1]],2),collapse=""),
                            paste(" CI = ",paste(c(round(ci(i$roc)[1],2),round(ci(i$roc)[3],2)),collapse = "-")))
      }
      
      #
      x <- data.table(name        = final_name,
                      AUC         = paste("AUC = ",
                                          round(i$roc$auc[[1]],2),"%",
                                          collapse=""),
                      Specificity = i$roc$specificities,
                      Sensitivity = i$roc$sensitivities,
                      FPR         = 100 - i$roc$specificities)
      x <- x[order(Sensitivity),]
      return(x)
    })
  roc_data_table <- rbindlist(roc_data_table)
  
  #generate plot
  p <- ggplot(roc_data_table,aes(x= Specificity,y = Sensitivity,colour = name)) + 
    geom_line(size = rocSize,linetype = rocStyle)                               +
    scale_x_reverse()                                                           + 
    #theme_bw()+
    geom_segment(x=-100, y=0, xend=0, yend=100, linetype=3, colour="grey75")    + 
    theme(legend.key.height=unit(2,"line"),
          legend.title    = element_blank(),
          legend.text     = element_text(size = 10),
          
          legend.position      = c(.45, .11+.11*length(validation_roc_list)), 
          legend.justification = c(0, 1))
  
  #return plot
  return(p)  
}

#~~~~~~~~~~~~~~~~~~~~~~~~~~~~~~~~~~~~~~~~~~~~~~~~~~~~~~~~~~~~~~~~~~~~~~~~~~~~~~~~~~~~~~
#fisher_enrichment_analysis
#~[by Francesco Vallania 05/03/2014]
#~this function computes enrichment analysis of labels in set vs tot
#######################################################################################
fisher_enrichment_analysis <- function(go_set,
                                       go_tot,
                                       unique_names=NULL){
  
  #build a contingency table
  go_tab           <- table(rbind(go_set,go_tot));
  
  #create dataframe
  go_df <- data.frame(label = row.names(go_tab),
                      set   = go_tab[,1],
                      total = go_tab[,2]);
  
  #create dataframe with names if it's available
  if(length(unique_names)>0){
    go_df <- data.frame(label = row.names(go_tab),
                        set   = go_tab[,1],
                        total = go_tab[,2],
                        name  = unique_names[c(match(row.names(go_tab),unique_names[,1])),2]);
  }  
  
  #p-values
  p_values_p <- sapply(1:nrow(go_df),function(i) phyper(go_df[i,2]-1,
                                                        go_df[i,3],
                                                        sum(go_df[-i,3]),
                                                        sum(go_df[,2]),
                                                        lower.tail=F));
  
  #compute bonferroni cut-off
  bonf <- rep(0,nrow(go_df));
  bonf[which(p_values_p<=(0.05/nrow(go_df)))] <- 1;
  
  #return go table with p-values
  go_df <- cbind(go_df,p_values_p,bonf);
  go_df <- go_df[order(go_df$p_values_p),];
  return(go_df);
}

#~~~~~~~~~~~~~~~~~~~~~~~~~~~~~~~~~~~~~~~~~~~~~~~~~~~~~~~~~~~~~~~~~~~~~~~~~~~~~~~~~~~~~~
#go_enrichment_analysis
#~[by Francesco Vallania 05/03/2014]
#~
#######################################################################################
go_enrichment_analysis <- function(gene_set,
                                   tot_gene = NULL,
                                   go_table = NULL){
  #load Hs go_table if it's by default
  if(is.null(go_table)){
    die("input GO table");  
  }
  
  #run code only if there is stuff
  if(length(gene_set)>0){
    #use all genes if it's by default
    if(length(tot_gene)==0){
      tot_gene <- unique(go_table$gene_symbol);
    }
    
    #collect GOs associated to gene set
    go_set       <- data.frame(go=go_table[which(go_table$gene_symbol %in% gene_set),2],set=1);
    go_tot       <- data.frame(go=go_table[which(go_table$gene_symbol %in% tot_gene),2],set=2);
    unique_names <- unique(go_table[,2:3])
    
    #generate and return fisher test table
    return(fisher_enrichment_analysis(go_set,go_tot,unique_names));
  }
  else{
    #if there is no gene return table with every gene with p-value of 1 
    unique_names <- unique(go_table[,2:3]);
    final_df     <- data.frame(label      = unique_names$go_id,
                               set        = 0,
                               total      = 0,
                               name       = unique_names$go_name,
                               p_values_p = 1,
                               bonf       = 0)
    return(final_df);
  }
}


#######################################################################################
#######################################################################################
#~~~~~~~~~~~~~~~~~~~~~~~~~~~~~~~~~~~~~~~~~~~~~~~~~~~~~~~~~~~~~~~~~~~~~~~~~~~~~~~~~~~~~~
#R Main
#~~~~~~~~~~~~~~~~~~~~~~~~~~~~~~~~~~~~~~~~~~~~~~~~~~~~~~~~~~~~~~~~~~~~~~~~~~~~~~~~~~~~~~
#######################################################################################
#######################################################################################

#~~~~~~~~~~~~~~~~~~~~~~~~~~~~~~~~~~~~~~~~~~~~~~~~~~~~~~~~~~~~~~~~~~~~~~~~~~~~~~~~~~~~~~
#~~~~~~~~~~~~~~~~~~~~~~~~~~~~~~~~~~~~~~~~~~~~~~~~~~~~~~~~~~~~~~~~~~~~~~~~~~~~~~~~~~~~~~
#Load data for Figure 4
#~~~~~~~~~~~~~~~~~~~~~~~~~~~~~~~~~~~~~~~~~~~~~~~~~~~~~~~~~~~~~~~~~~~~~~~~~~~~~~~~~~~~~~
#~~~~~~~~~~~~~~~~~~~~~~~~~~~~~~~~~~~~~~~~~~~~~~~~~~~~~~~~~~~~~~~~~~~~~~~~~~~~~~~~~~~~~~

#~~~~~~~~~~~~~~~~~~~~~~~~~~~~~~~~~~~~~~~~~~~~~~~~~~~~~~~~~~~~~~~~~~~~~~~~~~~~~~~~~~~~~~
#Load effect sizes from immunoStates datasets for monocyte gene signatures
#~~~~~~~~~~~~~~~~~~~~~~~~~~~~~~~~~~~~~~~~~~~~~~~~~~~~~~~~~~~~~~~~~~~~~~~~~~~~~~~~~~~~~~

#load effect size
outF <- 'expression_data/iS_effect_sizes_monocyte_genes.RData'
load(outF)

#load gene list
geneListMonoSub <- readRDS('expression_data/monocyte_subset_final_gene_list.rds')

#~~~~~~~~~~~~~~~~~~~~~~~~~~~~~~~~~~~~~~~~~~~~~~~~~~~~~~~~~~~~~~~~~~~~~~~~~~~~~~~~~~~~~~
#Supplementary Figure 3
#~~~~~~~~~~~~~~~~~~~~~~~~~~~~~~~~~~~~~~~~~~~~~~~~~~~~~~~~~~~~~~~~~~~~~~~~~~~~~~~~~~~~~~

#annotate rows
geneDisDf <- as.data.frame(geneListMonoSub[,gsub("_"," ",cellType)])
row.names(geneDisDf) <- geneListMonoSub$gene
colnames(geneDisDf) <- 'Cell-Specific Gene'

#annotate columns
cellLinDf <- as.data.frame(c(rep("myeoloid (monocytes)",2),
                             rep("lymphoid",4),
                             rep("myeoloid",3),
                             rep("lymphoid",1),
                             rep("progenitor",1),
                             rep("myeoloid",3),
                             rep("lymphoid",1),
                             rep("myeoloid",1),
                             rep("lymphoid",1),
                             rep("myeoloid",1),
                             rep("lymphoid",2)))
row.names(cellLinDf) <- colnames(iSgeneC)
colnames(cellLinDf) <- 'Cell Lineage'

annotation_colors = list('Cell-Specific Gene' = c('Classical'     = "#4E79A7",
                                                  'Non Classical' = "#E15759"),
                         'Cell Lineage'       = c('lymphoid'             = "dodgerblue",
                                                  'progenitor'           = 'khaki1',
                                                  'myeoloid'             = "lavenderblush",
                                                  'myeoloid (monocytes)' = "deeppink1"))
pdf('pdfs/20201201_supplementary_figure3.pdf',
    height=5*1.3,width=6*1.1)
pH_sf3 <- pheatmap(rbind(iSgeneC,iSgeneN),
                   show_rownames = T,
                   show_colnames = T,
                   border_color  = NA,
                   cluster_rows  = F, 
                   annotation_row    = geneDisDf,
                   annotation_col    = cellLinDf, 
                   annotation_colors = annotation_colors,
                   color=grDevices::colorRampPalette(c("navyblue","white","orangered"))(1000))

#rotate dendrogram
orderedL <- row.names(cellLinDf)[order(cellLinDf$`Cell Lineage`)][c(19:18,11,10,13,12,17:14,20,8:1,9)]
col_dend <- pH_sf3[[2]]
col_dend <- dendextend::rotate(col_dend,
                               order = orderedL)


pheatmap(rbind(iSgeneC,iSgeneN),
              show_rownames = T,
              show_colnames = T,
              border_color  = NA,
              cluster_rows  = F,
              cluster_cols=as.hclust(col_dend),
              annotation_row    = geneDisDf,
              annotation_col    = cellLinDf,
              annotation_colors = annotation_colors,
              color=grDevices::colorRampPalette(c("navyblue",
                                                  "white",
                                                  "orangered"))(1000))
dev.off()


#~~~~~~~~~~~~~~~~~~~~~~~~~~~~~~~~~~~~~~~~~~~~~~~~~~~~~~~~~~~~~~~~~~~~~~~~~~~~~~~~~~~~~~
#Load processed raw expression data for immunoStates
#~~~~~~~~~~~~~~~~~~~~~~~~~~~~~~~~~~~~~~~~~~~~~~~~~~~~~~~~~~~~~~~~~~~~~~~~~~~~~~~~~~~~~~

#compute scores 
#load IS data.table with quantile normalized expression data
load(file = 'expression_data/iS_raw_data_monocyte_genes.RData')

#~~~~~~~~~~~
iSclassDT[,meanV:=mean(V1,na.rm=T),midCell]
iSnclassDT[,meanV:=mean(V1,na.rm=T),midCell]

#~~~~~~~~~~~
iSclassDT_cmb <- merge(iSclassDT,iSnclassDT,by=c('gsm',"midCell"))
setnames(iSclassDT_cmb,c("gsm","midCell","Classical",
                         "meanC","Non-Classical","meanN"))

#combine signatures by taking the ration
iSclassDT_cmb[,mix:=`Non-Classical`/Classical]
iSclassDT_cmb[,meanM:=mean(mix,na.rm=T),midCell]


#~~~~~~~~~~~~~~~~~~~~~~~~~~~~~~~~~~~~~~~~~~~~~~~~~~~~~~~~~~~~~~~~~~~~~~~~~~~~~~~~~~~~~~
#Create Figure 4 plots
#~~~~~~~~~~~~~~~~~~~~~~~~~~~~~~~~~~~~~~~~~~~~~~~~~~~~~~~~~~~~~~~~~~~~~~~~~~~~~~~~~~~~~~

iSclassDT_cmb[,newlabel:='other immune cells']
iSclassDT_cmb[grep("^CD14_positive_mono",midCell),newlabel:='classical monocytes']
iSclassDT_cmb[grep("^CD16_positive_mono",midCell),newlabel:='non-classical monocytes']

iSclassDT_cmbM <- melt.data.table(iSclassDT_cmb[grep("global",midCell,invert = T)],
                                  id.vars = c("gsm","midCell","meanC","meanN",'meanM',"newlabel"))

serr <- function(inputV){sd(inputV)/sqrt(length(inputV))}
iSclassDT_cmbM <- iSclassDT_cmbM[!is.na(value)]
iSclassDT_cmbM[,mean:=mean(value,na.rm=T),.(newlabel,variable)]
iSclassDT_cmbM[,serr:=serr(value),.(newlabel,variable)]

pC <- ggplot(iSclassDT_cmbM[variable=='Classical'],
             aes(x     = newlabel,
                 color = newlabel,
                 ymax  = mean+serr,
                 ymin  = mean-serr,
                 y     = value))   + 
  geom_quasirandom()       +
  geom_point(aes(y=mean),color='black') +
  geom_errorbar(color='black',width=0.2) +
  theme_tufte(base_family = 'Helvetica') +
  xlab("immune cell type") +
  labs(color="") +
  scale_color_manual(values = c("#4E79A7","#E15759",'grey80')) +
  theme(axis.text.x = element_blank(),
        axis.line.x = element_line(color='black'),
        axis.ticks.x = element_blank(),
        axis.line.y = element_line(color='black'))+
  ylab("cMSS scores") + 
  geom_signif(comparisons = list(c('classical monocytes',
                                   'non-classical monocytes'),
                                 c('classical monocytes',
                                   'other immune cells'),
                                 c('other immune cells',
                                   'non-classical monocytes')),
              y_position = c(10.5,11,1),
              tip_length = c(.01,.01,-.01),
              test       = 't.test',
              color      = 'black')

pNC <- ggplot(iSclassDT_cmbM[variable=='Non-Classical'],
              aes(x     = newlabel,
                  color = newlabel,
                  ymax  = mean+serr,
                  ymin  = mean-serr,
                  y     = value))   + 
  geom_quasirandom()       +
  geom_point(aes(y=mean),color='black') +
  geom_errorbar(color='black',width=0.2) +
  theme_tufte(base_family = 'Helvetica') +
  xlab("immune cell type") +
  labs(color="") +
  scale_color_manual(values = c("#4E79A7","#E15759",'grey80')) +
  theme(axis.text.x = element_blank(),
        axis.line.x = element_line(color='black'),
        axis.ticks.x = element_blank(),
        axis.line.y = element_line(color='black'))+
  ylab("ncMSS scores") + 
  geom_signif(comparisons = list(c('classical monocytes',
                                   'non-classical monocytes'),
                                 c('classical monocytes',
                                   'other immune cells'),
                                 c('other immune cells',
                                   'non-classical monocytes')),
              y_position = c(10.5,11,1),
              tip_length = c(.01,.01,-.01),
              test       = 't.test',
              color      = 'black')


pdf('pdfs/20201201_figure4_A.pdf',
    width=5,height=5.5)
plot(pC)
dev.off()

pdf('pdfs/20201201_figure4_B.pdf',
    width=5,height=5.5)
plot(pNC)
dev.off()

#~~~~~~~~~~~~~~~~~~~~~~~~~~~~~~~~~~~~~~~~~~~~~~~~~~~~~~~~~~~~~~~~~~~~~~~~~~~~~~~~~~~~~~
#Supplementary Figure 4: Compute ROC curves
#~~~~~~~~~~~~~~~~~~~~~~~~~~~~~~~~~~~~~~~~~~~~~~~~~~~~~~~~~~~~~~~~~~~~~~~~~~~~~~~~~~~~~~

iSclassDT_cmb[,class:=0]
iSclassDT_cmb[newlabel=='classical monocytes',class:=1]

rocC_CvsAC <- roc(iSclassDT_cmb$class,
                  iSclassDT_cmb$Classical)

rocC_CvsNC <- roc(iSclassDT_cmb[grep("monocytes",newlabel)]$class,
                  iSclassDT_cmb[grep("monocytes",newlabel)]$Classical)

rocC_CvsOI <- roc(iSclassDT_cmb[grep("non-classical",newlabel,invert = T)]$class,
                  iSclassDT_cmb[grep("non-classical",newlabel,invert = T)]$Classical)

rocC_NCvsC <- roc(iSclassDT_cmb[grep("monocytes",newlabel)]$class,
                  iSclassDT_cmb[grep("monocytes",newlabel)]$`Non-Classical`)

iSclassDT_cmb[,class:=0]
iSclassDT_cmb[newlabel=='non-classical monocytes',class:=1]

rocC_NCvsAC <- roc(iSclassDT_cmb$class,
                   iSclassDT_cmb$`Non-Classical`)

rocC_NCvsOI <- roc(iSclassDT_cmb[grep("^classical",newlabel,invert = T)]$class,
                   iSclassDT_cmb[grep("^classical",newlabel,invert = T)]$`Non-Classical`)

#~~~~~~~~~~~~~~~~~~
pdf('pdfs/20201201_supplementary_figure4.pdf',
    height=3,width=5.5)
rocGGMultiRoc(list(list(name='cMSS', roc = rocC_CvsAC),
                   list(name='ncMSS',roc = rocC_NCvsAC))) + 
  theme_bw() + #geom_abline(slope=1,linetype=2,intercept = 1) +
  scale_color_manual(values=c('#4E79A7','#E15759')) + labs(color='')
dev.off()
```
