## Supplementary material for "Multicohort Analysis of Publicly-available Monocyte Expression Data Identifies Gene Signatures to Accurately Monitor Subset-specific Changes in Human Diseases": code for analyses and figures: monocyte_paper_figure_5.R

```
#######################################################################################
#Monocyte Signatures Analysis
#by Francesco Vallania, January 30st 2016 @ 9:53pm
#@ Stanford
#~~~~~~~~~~~~~~~~~~~~~~~~~~~~~~~~~~~~~~~~~~~~~~~~~~~~~~~~~~~~~~~~~~~~~~~~~~~~~~~~~~~~~~
#the purpose of this code is generate plots for Figure 5
#######################################################################################

#######################################################################################
#R Functions
#~~~~~~~~~~~~~~~~~~~~~~~~~~~~~~~~~~~~~~~~~~~~~~~~~~~~~~~~~~~~~~~~~~~~~~~~~~~~~~~~~~~~~~
#######################################################################################

#~~~~~~~~~~~~~~~~~~~~~~~~~~~~~~~~~~~~~~~~~~~~~~~~~~~~~~~~~~~~~~~~~~~~~~~~~~~~~~~~~~~~~~
#garbage cleaning
#~~~~~~~~~~~~~~~~~~~~~~~~~~~~~~~~~~~~~~~~~~~~~~~~~~~~~~~~~~~~~~~~~~~~~~~~~~~~~~~~~~~~~~
rm(list=ls())

#~~~~~~~~~~~~~~~~~~~~~~~~~~~~~~~~~~~~~~~~~~~~~~~~~~~~~~~~~~~~~~~~~~~~~~~~~~~~~~~~~~~~~~
#~Load all the necessary libraries
#~~~~~~~~~~~~~~~~~~~~~~~~~~~~~~~~~~~~~~~~~~~~~~~~~~~~~~~~~~~~~~~~~~~~~~~~~~~~~~~~~~~~~~

library(data.table)
library(reshape2)
library(parallel)
library(ggplot2)
library(pROC)
library(BiocGenerics)
library(pheatmap)
library(ggthemes)
library(ggsignif)
library(ggbeeswarm)
library(readxl)
library(DESeq2)
library(AnnotationDbi)

library(MetaIntegrator)


#~~~~~~~~~~~~~~~~~~~~~~~~~~~~~~~~~~~~~~~~~~~~~~~~~~~~~~~~~~~~~~~~~~~~~~~~~~~~~~~~~~~~~~
#extractDataForGenesDT
#######################################################################################
#Quickly create gene matrix for a given subset of genes or all
#[note: much faster than before because through data.table]
#######################################################################################
extractDataForGenesDT <- function(GEM,geneNames = NULL,promiscProbes = TRUE){
  
  #run stuff 
  if(GEM$key_comment!= "Annotation absent" && GEM$exp_comment != "Expression data is missing"){
    if(length(which(!is.na(GEM$keys)))>0){
      
      #update-> split keys that map to multiple genes
      #[note: use unlist2 from AnnotationDbi so that names are fine]
      #[note: make it into data.table]
      keyVector  <- NULL
      
      #choose to either include/exclude probes that map to more than one gene
      if(promiscProbes==TRUE){
        keyVector  <- unlist2(sapply(GEM$keys,function(i)strsplit(i,",")))
      }else{
        keyVector  <- GEM$keys[grep(",",GEM$keys,invert = T)]
      }
      
      #create keyVector DT and remove NA cases and empty cases as well
      keyVectorDT<- data.table(probe = names(keyVector),gene = keyVector)
      keyVectorDT<- keyVectorDT[!is.na(gene)]
      keyVectorDT<- keyVectorDT[gene!='']
      
      #expression DataTable
      exprDT    <- as.data.table(GEM$expr,
                                 keep.rownames = T)
      
      #merge them and remove probes [since they are now @ a gene leve]
      mergedDT  <- merge(exprDT,
                         keyVectorDT,
                         by.x = 'rn',
                         by.y = 'probe')
      
      if(nrow(mergedDT)>0){
        
        mergedDT  <- mergedDT[,rn:=NULL]
        
        #filter genes
        if(!is.null(geneNames)){
          mergedDT  <- mergedDT[gene %in% geneNames]
        }
        
        #compute mean and cast
        meanDT    <- melt.data.table(
          mergedDT,
          id.vars = 'gene')[,mean(value),.(gene,variable)]
        meanDT    <- dcast.data.table(
          meanDT,
          formula = gene~variable,value.var = 'V1')
        
        #covert data.table to matrix
        tRn    <- meanDT$gene
        outMat <- as.matrix(meanDT[,gene:=NULL])
        row.names(outMat) <- tRn
        
        #return matrix
        return(outMat)
      }else{
        return(NA)
      }
    }else{
      return(NA)
    }
  }else{
    return(NA)
  }
}

#######################################################################################
#######################################################################################
#~~~~~~~~~~~~~~~~~~~~~~~~~~~~~~~~~~~~~~~~~~~~~~~~~~~~~~~~~~~~~~~~~~~~~~~~~~~~~~~~~~~~~~
#R Main
#~~~~~~~~~~~~~~~~~~~~~~~~~~~~~~~~~~~~~~~~~~~~~~~~~~~~~~~~~~~~~~~~~~~~~~~~~~~~~~~~~~~~~~
#######################################################################################
#######################################################################################

#~~~~~~~~~~~~~~~~~~~~~~~~~~~~~~~~~~~~~~~~~~~~~~~~~~~~~~~~~~~~~~~~~~~~~~~~~~~~~~~~~~~~~~
#Load Monocyte genes
#~~~~~~~~~~~~~~~~~~~~~~~~~~~~~~~~~~~~~~~~~~~~~~~~~~~~~~~~~~~~~~~~~~~~~~~~~~~~~~~~~~~~~~

#load gene list
geneListMonoSub <- readRDS('expression_data/monocyte_subset_final_gene_list.rds')

#~~~~~~~~~~~~~~~~~~~~~~~~~~~~~~~~~~~~~~~~~~~~~~~~~~~~~~~~~~~~~~~~~~~~~~~~~~~~~~~~~~~~~~
#Figure 5A: RA
#~~~~~~~~~~~~~~~~~~~~~~~~~~~~~~~~~~~~~~~~~~~~~~~~~~~~~~~~~~~~~~~~~~~~~~~~~~~~~~~~~~~~~~

#whole blood RA
outputFile   <- 'expression_data/GSE93272_gem.rds'
if(!file.exists(outputFile)){
  GSE93272     <- getGEOData('GSE93272')
  saveRDS(GSE93272,outputFile)
}
GSE93272 <- readRDS(outputFile)

GSE93272phDT <- as.data.table(GSE93272$originalData$GSE93272$pheno,keep.rownames = T)

#extract expression matrix
GSE93272DT  <- extractDataForGenesDT(GSE93272$originalData$GSE93272)
GSE93272DTm <- setnames(as.data.table(melt(GSE93272DT)),c("gene","gsm",'value'))

GSE93272_sign    <- rbindlist(
  lapply(geneListMonoSub[,.N,cellType]$cellType,
         function(cellID)
           GSE93272DTm[gene %in% geneListMonoSub[cellType==cellID]$gene,
                       mean(value),gsm][,cellType:=cellID]))

GSE93272_sign <- merge(
  GSE93272_sign,
  GSE93272phDT[,c('rn','title','disease state:ch1',
                  'characteristics_ch1.5',
                  'sjc28:ch1',
                  'characteristics_ch1',
                  'sdai:ch1',
                  'pain.vas:ch1',
                  'symptom.vas:ch1')],
  by.x='gsm',by.y='rn')

serr <- function(inputV){{sd(inputV)/sqrt(length(inputV))}}
GSE93272_sign[,scaleZ:=scale(V1),cellType]
GSE93272_sign[,mean:=mean(scaleZ),.(cellType,`disease state:ch1`)]
GSE93272_sign[,serr:=serr(scaleZ),.(cellType,`disease state:ch1`)]

#create figure panels
p_GSE93272mC <- ggplot(GSE93272_sign[cellType=="Classical"],
                       aes(x     = `disease state:ch1`,#gsub("_BL","",`visit:ch1`),
                           y     =  scaleZ,
                           ymax  = mean+serr,
                           ymin  = mean-serr,
                           color = `disease state:ch1`)) + 
  geom_quasirandom() + 
  scale_color_manual(values=c('#FF9DA7','#B07AA1')) +
  labs(color='')+
  geom_errorbar(color='black',width=0.1) + 
  geom_point(aes(y =mean),color='black') +
  theme_tufte(base_family = 'Helvetica') +
  theme(axis.line.x = element_line(),
        axis.line.y = element_line()) +
  xlab("disease") + 
  ylab("cMSS score") + 
  ggtitle(paste("GSE93272 | Rheumatoid Arthritis (Whole Blood)\n",
                GSE93272_sign[,.N,gsm][,.N],
                " samples",sep='')) +
  geom_signif(comparisons = list(c("healthy control", "RA")),color='black')

p_GSE93272mNC <- ggplot(GSE93272_sign[cellType=='Non_Classical'],
                        aes(x     = `disease state:ch1`,#gsub("_BL","",`visit:ch1`),
                            y     =  scaleZ,
                            ymax  = mean+serr,
                            ymin  = mean-serr,
                            color = `disease state:ch1`)) + 
  geom_quasirandom() + 
  scale_color_manual(values=c('#FF9DA7','#B07AA1')) +
  labs(color='')+
  geom_errorbar(color='black',width=0.1) + 
  geom_point(aes(y =mean),color='black') +
  theme_tufte(base_family = 'Helvetica') +
  theme(axis.line.x = element_line(),
        axis.line.y = element_line()) +
  xlab("disease") + 
  ylab("ncMSS score") + 
  ggtitle(paste("GSE93272 | Rheumatoid Arthritis (Whole Blood)\n",
                GSE93272_sign[,.N,gsm][,.N],
                " samples",sep='')) +
  geom_signif(comparisons = list(c("healthy control", "RA")),color='black')


pdf('pdfs/20201201_figure5_A_classical.pdf',
    width = 5,height = 5)
plot(p_GSE93272mC)
dev.off()

pdf('pdfs/20201201_figure5_A_nonclassical.pdf',
    width = 5,height = 5)
plot(p_GSE93272mNC)
dev.off()


#~~~~~~~~~~~~~~~~~~~~~~~~~~~~~~~~~~~~~~~~~~~~~~~~~~~~~~~~~~~~~~~~~~~~~~~~~~~~~~~~~~~~~~
#Figure 5B: SJIA 
#~~~~~~~~~~~~~~~~~~~~~~~~~~~~~~~~~~~~~~~~~~~~~~~~~~~~~~~~~~~~~~~~~~~~~~~~~~~~~~~~~~~~~~

#donwload data
outputFile   <- 'expression_data/GSE80060_gem.rds'
if(!file.exists(outputFile)){
  trial_sjia     <- getGEOData('GSE80060')
  saveRDS(trial_sjia,outputFile)
}
trial_sjia <- readRDS(outputFile)

#extract expression matrix
trial_sjiaDT  <- extractDataForGenesDT(trial_sjia$originalData$GSE80060)
trial_sjiaDTm <- setnames(as.data.table(melt(trial_sjiaDT)),c("gene","gsm",'value'))


sign_trial    <- rbindlist(
  lapply(geneListMonoSub[,.N,cellType]$cellType,
         function(cellID)
           trial_sjiaDTm[gene %in% geneListMonoSub[cellType==cellID]$gene,
                         mean(value),
                         gsm][,cellType:=cellID]))

#look @ pheno data
sign_trial_pheno <- merge(as.data.table(trial_sjia$originalData$GSE80060$pheno),
                          sign_trial,
                          by.x='geo_accession',
                          by.y='gsm')
sign_trial_pheno[,response:='Non-Responder (ACR < 50)']
sign_trial_pheno[`acr response at day 15:ch1`>=50,response:='Responder (ACR >= 50)']

#make better plot
serr <- function(inputV){{sd(inputV)/sqrt(length(inputV))}}
sign_trial_pheno[,mean:=mean(V1),.(cellType,`treatment:ch1`,`visit:ch1`,response)]
sign_trial_pheno[,serr:=serr(V1),.(cellType,`treatment:ch1`,`visit:ch1`,response)]

sign_trial_pheno[,label:="Baseline\nResponder"]
sign_trial_pheno[response=='Non-Responder (ACR < 50)' & `visit:ch1` == 'Day1_BL',
                 label:="Baseline\nNon-Responder"]
sign_trial_pheno[response!='Non-Responder (ACR < 50)' & `visit:ch1` != 'Day1_BL',
                 label:="Day 3\nResponder"]
sign_trial_pheno[response=='Non-Responder (ACR < 50)' & `visit:ch1` != 'Day1_BL',
                 label:="Day 3\nNon-Responder"]


#Split plot of Figure 5 B into two separate panels
p_trial_C <- ggplot(
  sign_trial_pheno[
    grep("^Classical",
         cellType)][!`treatment:ch1`%in% c('none',
                                           "Placebo")][,scaleZ:=scale(V1),cellType],
  aes(x     = gsub("_BL","",`visit:ch1`),
      y     = scaleZ,
      color = response)) + 
  geom_quasirandom(dodge.width = 1) + 
  scale_color_tableau() +
  labs(color='')+
  theme_tufte(base_family = 'Helvetica') +
  theme(axis.line.x = element_line(),
        axis.line.y = element_line()) +
  xlab("Time") + 
  ylab("cMSS score") + #ggtitle("GSE80060 | Canakinumab SJIA Trial") +
  ggtitle(paste("GSE80060 | Canakinumab SJIA Trial (Whole Blood)\n",
                sign_trial_pheno[,.N,geo_accession][,.N],
                " samples",sep='')) +
  geom_signif(comparisons = list(c("Day1", "Day3")),color='black')


p_trial_NC <- ggplot(
  sign_trial_pheno[grep("^Non_Classical",
                        cellType)][
                          !`treatment:ch1`%in% c('none',
                                                 "Placebo")][,scaleZ:=scale(V1),cellType],
  aes(x     = gsub("_BL","",`visit:ch1`),
      y     = scaleZ,
      color = response)) + 
  geom_quasirandom(dodge.width = 1) + 
  scale_color_tableau() +
  labs(color='')+
  theme_tufte(base_family = 'Helvetica') +
  theme(axis.line.x = element_line(),
        axis.line.y = element_line()) +
  xlab("Time") + 
  ylab("ncMSS score") +
  ggtitle(paste("GSE80060 | Canakinumab SJIA Trial (Whole Blood)\n",
                sign_trial_pheno[,.N,geo_accession][,.N],
                " samples",sep='')) +
  geom_signif(comparisons = list(c("Day1", "Day3")),color='black')


pdf('pdfs/20201201_figure5_B_classical.pdf',
    width = 6,height = 5)
plot(p_trial_C)
dev.off()


pdf('pdfs/20201201_figure5_B_nonclassical.pdf',
    width = 6,height = 5)
plot(p_trial_NC)
dev.off()
```
