## Supplementary material for "Multicohort Analysis of Publicly-available Monocyte Expression Data Identifies Gene Signatures to Accurately Monitor Subset-specific Changes in Human Diseases": code for analyses and figures: monocyte_paper_figure_6.R

```
#######################################################################################
#Monocyte Signatures Analysis
#by Francesco Vallania, January 30st 2016 @ 9:53pm
#@ Stanford
#~~~~~~~~~~~~~~~~~~~~~~~~~~~~~~~~~~~~~~~~~~~~~~~~~~~~~~~~~~~~~~~~~~~~~~~~~~~~~~~~~~~~~~
#the purpose of this code is generate plots for Figure 5
#######################################################################################

#######################################################################################
#R Functions
#~~~~~~~~~~~~~~~~~~~~~~~~~~~~~~~~~~~~~~~~~~~~~~~~~~~~~~~~~~~~~~~~~~~~~~~~~~~~~~~~~~~~~~
#######################################################################################

#~~~~~~~~~~~~~~~~~~~~~~~~~~~~~~~~~~~~~~~~~~~~~~~~~~~~~~~~~~~~~~~~~~~~~~~~~~~~~~~~~~~~~~
#garbage cleaning
#~~~~~~~~~~~~~~~~~~~~~~~~~~~~~~~~~~~~~~~~~~~~~~~~~~~~~~~~~~~~~~~~~~~~~~~~~~~~~~~~~~~~~~
rm(list=ls())

#~~~~~~~~~~~~~~~~~~~~~~~~~~~~~~~~~~~~~~~~~~~~~~~~~~~~~~~~~~~~~~~~~~~~~~~~~~~~~~~~~~~~~~
#~Load all the necessary libraries
#~~~~~~~~~~~~~~~~~~~~~~~~~~~~~~~~~~~~~~~~~~~~~~~~~~~~~~~~~~~~~~~~~~~~~~~~~~~~~~~~~~~~~~

library(data.table)
library(reshape2)
library(parallel)
library(ggplot2)
library(pROC)
library(BiocGenerics)
library(pheatmap)
library(ggthemes)
library(ggsignif)
library(ggbeeswarm)
library(readxl)
library(DESeq2)
library(AnnotationDbi)

library(MetaIntegrator)


#~~~~~~~~~~~~~~~~~~~~~~~~~~~~~~~~~~~~~~~~~~~~~~~~~~~~~~~~~~~~~~~~~~~~~~~~~~~~~~~~~~~~~~
#extractDataForGenesDT
#######################################################################################
#Quickly create gene matrix for a given subset of genes or all
#[note: much faster than before because through data.table]
#######################################################################################
extractDataForGenesDT <- function(GEM,geneNames = NULL,promiscProbes = TRUE){
  
  #run stuff 
  if(GEM$key_comment!= "Annotation absent" && GEM$exp_comment != "Expression data is missing"){
    if(length(which(!is.na(GEM$keys)))>0){
      
      #update-> split keys that map to multiple genes
      #[note: use unlist2 from AnnotationDbi so that names are fine]
      #[note: make it into data.table]
      keyVector  <- NULL
      
      #choose to either include/exclude probes that map to more than one gene
      if(promiscProbes==TRUE){
        keyVector  <- unlist2(sapply(GEM$keys,function(i)strsplit(i,",")))
      }else{
        keyVector  <- GEM$keys[grep(",",GEM$keys,invert = T)]
      }
      
      #create keyVector DT and remove NA cases and empty cases as well
      keyVectorDT<- data.table(probe = names(keyVector),gene = keyVector)
      keyVectorDT<- keyVectorDT[!is.na(gene)]
      keyVectorDT<- keyVectorDT[gene!='']
      
      #expression DataTable
      exprDT    <- as.data.table(GEM$expr,keep.rownames = T)
      
      #merge them and remove probes [since they are now @ a gene leve]
      mergedDT  <- merge(exprDT,keyVectorDT,by.x='rn',by.y='probe')
      
      if(nrow(mergedDT)>0){
        
        mergedDT  <- mergedDT[,rn:=NULL]
        
        #filter genes
        if(!is.null(geneNames)){
          mergedDT  <- mergedDT[gene %in% geneNames]
        }
        
        #compute mean and cast
        meanDT    <- melt.data.table(mergedDT,id.vars = 'gene')[,mean(value),.(gene,variable)]
        meanDT    <- dcast.data.table(meanDT,formula = gene~variable,value.var = 'V1')
        
        #covert data.table to matrix
        tRn    <- meanDT$gene
        outMat <- as.matrix(meanDT[,gene:=NULL])
        row.names(outMat) <- tRn
        
        #return matrix
        return(outMat)
      }else{
        return(NA)
      }
    }else{
      return(NA)
    }
  }else{
    return(NA)
  }
}

#######################################################################################
#######################################################################################
#~~~~~~~~~~~~~~~~~~~~~~~~~~~~~~~~~~~~~~~~~~~~~~~~~~~~~~~~~~~~~~~~~~~~~~~~~~~~~~~~~~~~~~
#R Main
#~~~~~~~~~~~~~~~~~~~~~~~~~~~~~~~~~~~~~~~~~~~~~~~~~~~~~~~~~~~~~~~~~~~~~~~~~~~~~~~~~~~~~~
#######################################################################################
#######################################################################################

#~~~~~~~~~~~~~~~~~~~~~~~~~~~~~~~~~~~~~~~~~~~~~~~~~~~~~~~~~~~~~~~~~~~~~~~~~~~~~~~~~~~~~~
#Load data
#~~~~~~~~~~~~~~~~~~~~~~~~~~~~~~~~~~~~~~~~~~~~~~~~~~~~~~~~~~~~~~~~~~~~~~~~~~~~~~~~~~~~~~

#load gene list
geneListMonoSub <- readRDS('expression_data/monocyte_subset_final_gene_list.rds')


newman_Flow_DT  <- fread('expression_data/PBMCs-Fig3a-Flow-Cytometry.txt')
newman_GExp_DT  <- fread('expression_data/PBMCs-Fig3a-HumanHT-12-V4.txt')

newman_GExp_DTm <- melt.data.table(newman_GExp_DT,id.vars = 'GeneSym')
newman_Flow_DTm <- melt.data.table(newman_Flow_DT,id.vars = 'ID')
setnames(newman_Flow_DTm,c('ID',"cellType","proportion"))
setnames(newman_GExp_DTm,c('Gene',"ID","value"))

#compute signature scores [do it in log scale]
sign_newman    <- rbindlist(
  lapply(geneListMonoSub[,.N,cellType]$cellType,
         function(cellID)
           newman_GExp_DTm[Gene %in% geneListMonoSub[cellType==cellID]$gene,
                           mean(log2(value)),ID][,cellType:=cellID]))

sign_newman_flow <- merge(sign_newman,
                          newman_Flow_DTm[cellType=='Monos'],
                          by='ID')

#~~~~~~~~~~~~~~~~~~~~~~~~~~~~~~~~~~~~~~~~~~~~~~~~~~~~~~~~~~~~~~~~~~~~~~~~~~~~~~~~~~~~~~
#Create Figure
#~~~~~~~~~~~~~~~~~~~~~~~~~~~~~~~~~~~~~~~~~~~~~~~~~~~~~~~~~~~~~~~~~~~~~~~~~~~~~~~~~~~~~~

p_newman <- ggplot(sign_newman_flow[cellType.x=='Classical'],
                   aes(x=V1,y=proportion)) + 
  geom_point(size=3) + 
  theme_bw() +
  scale_y_log10() + 
  geom_smooth(method   = 'lm',
              linetype = 2,
              color    = 'grey40',
              fill     = NA) +
  xlab("cMSS score") +
  ylab("monocyte fraction by flow-cytometry (%)") + 
  ggtitle(paste("GSE65136 | Healthy PBMCs\n",
                sign_newman_flow[,.N,ID][,.N],
                " samples",sep='')) +
  annotate(
    'text',
    label= paste(
      paste("r =",
            round(sign_newman_flow[
              cellType.x=='Classical'][,cor(V1,log2(proportion))],2)),
      paste("p =",
            round(sign_newman_flow[
              cellType.x=='Classical'][,cor.test(V1,
                                                 log2(proportion))]$p.value,4)),
      sep='\n'),
    size = 6,
    x    = 7.4,
    y    = 5)


pdf('pdfs/20201201_figure6.pdf',
    width = 3.5,height = 3.7)
plot(p_newman)
dev.off()
```
