## Supplementary material for "Multicohort Analysis of Publicly-available Monocyte Expression Data Identifies Gene Signatures to Accurately Monitor Subset-specific Changes in Human Diseases": code for analyses and figures: monocyte_paper_figure_7.R

```
#######################################################################################
#Monocyte Signatures Analysis
#by Francesco Vallania, January 30st 2016 @ 9:53pm
#@ Stanford
#~~~~~~~~~~~~~~~~~~~~~~~~~~~~~~~~~~~~~~~~~~~~~~~~~~~~~~~~~~~~~~~~~~~~~~~~~~~~~~~~~~~~~~
#the purpose of this code is generate plots for Figure 7
#######################################################################################

#######################################################################################
#R Functions
#~~~~~~~~~~~~~~~~~~~~~~~~~~~~~~~~~~~~~~~~~~~~~~~~~~~~~~~~~~~~~~~~~~~~~~~~~~~~~~~~~~~~~~
#######################################################################################

#~~~~~~~~~~~~~~~~~~~~~~~~~~~~~~~~~~~~~~~~~~~~~~~~~~~~~~~~~~~~~~~~~~~~~~~~~~~~~~~~~~~~~~
#garbage cleaning
#~~~~~~~~~~~~~~~~~~~~~~~~~~~~~~~~~~~~~~~~~~~~~~~~~~~~~~~~~~~~~~~~~~~~~~~~~~~~~~~~~~~~~~
rm(list=ls())

#~~~~~~~~~~~~~~~~~~~~~~~~~~~~~~~~~~~~~~~~~~~~~~~~~~~~~~~~~~~~~~~~~~~~~~~~~~~~~~~~~~~~~~
#~Load all the necessary libraries
#~~~~~~~~~~~~~~~~~~~~~~~~~~~~~~~~~~~~~~~~~~~~~~~~~~~~~~~~~~~~~~~~~~~~~~~~~~~~~~~~~~~~~~

library(data.table)
library(reshape2)
library(parallel)
library(ggplot2)
library(pROC)
library(BiocGenerics)
library(pheatmap)
library(ggthemes)
library(ggsignif)
library(ggbeeswarm)
library(readxl)
library(MetaIntegrator)
library(rmeta)

#######################################################################################
#R Functions
#~~~~~~~~~~~~~~~~~~~~~~~~~~~~~~~~~~~~~~~~~~~~~~~~~~~~~~~~~~~~~~~~~~~~~~~~~~~~~~~~~~~~~~
#######################################################################################

#imported from MetaIntegrator package
getES <- function(v, g){
  ## function to calculate basic statistics for two-class comparison for a single gene
  ## v: vector of samples (one row from an expression matrix)
  ## g: vector of composed of 0,1 values for every sample in v; defines case or control
  ## 
  ## encoding: CASES should be designated with the value 1, and CONTROLS with the value 0 to preserve the 
  ## directionality of the equation (mean of cases - mean of controls)
  
  # check to make sure every sample in expression matrix is assigned case or control value
  if ( length(v) != length(g) ) {
    stop("number of samples in expression matrix is different from length of class vector -- exiting")
  } 
  
  .cleanNA <- function(x) return( x[!is.na(x) & is.finite(x) ] )
  
  x <- .cleanNA( v[ which(g==1) ] )
  y <- .cleanNA( v[ which(g==0) ] )
  
  n1 <- length(x); n2 <- length(y)
  if( n1 < 2 | n2 < 2 )
    return( c(n1=NA, m1=NA, sd1=NA,
              n2=NA, m2=NA, sd2=NA,
              diff=NA, pooled.sd=NA,
              g=NA, se.g=NA) )
  
  #compute mean difference
  m1 <- mean(x)
  m2 <- mean(y)
  diff <- m1 - m2
  
  #compute pooled standard deviation
  sd1  <- stats::sd(x)
  sd2  <- stats::sd(y)
  sp   <- sqrt( ( (n1-1)*sd1^2 + (n2-1)*sd2^2 )/( n1 + n2 - 2 ) )
  
  #correct for small sample size [from Cohen's D to Hedge's G]
  cf   <- 1 - 3/( 4*(n1 + n2) - 9 ) # small sample size correction
  g    <- cf * diff/sp
  se.g <- sqrt( (n1+n2)/(n1*n2) + 0.5*g^2 /(n1+n2-3.94) )
  
  return( c(n1=n1, m1=m1, sd1=sd1,
            n2=n2, m2=m2, sd2=sd2,
            diff=diff, pooled.sd=sp,
            g=g, se.g=se.g) ) 
}

#######################################################################################
#######################################################################################
#~~~~~~~~~~~~~~~~~~~~~~~~~~~~~~~~~~~~~~~~~~~~~~~~~~~~~~~~~~~~~~~~~~~~~~~~~~~~~~~~~~~~~~
#R Main
#~~~~~~~~~~~~~~~~~~~~~~~~~~~~~~~~~~~~~~~~~~~~~~~~~~~~~~~~~~~~~~~~~~~~~~~~~~~~~~~~~~~~~~
#######################################################################################
#######################################################################################

#~~~~~~~~~~~~~~~~~~~~~~~~~~~~~~~~~~~~~~~~~~~~~~~~~~~~~~~~~~~~~~~~~~~~~~~~~~~~~~~~~~~~~~
#Load up results from discovery
#~~~~~~~~~~~~~~~~~~~~~~~~~~~~~~~~~~~~~~~~~~~~~~~~~~~~~~~~~~~~~~~~~~~~~~~~~~~~~~~~~~~~~~

#load genes selected for signatures
geneListMonoSubset <- readRDS('expression_data/monocyte_subset_final_gene_list.rds')

#~~~~~~~~~~~~~~~~~~~~~~~~~~~~~~~~~~~~~~~~~~~~~~~~~~~~~~~~~~~~~~~~~~~~~~~~~~~~~~~~~~~~~~
#Load up results from validation (RNA-seq)
#~~~~~~~~~~~~~~~~~~~~~~~~~~~~~~~~~~~~~~~~~~~~~~~~~~~~~~~~~~~~~~~~~~~~~~~~~~~~~~~~~~~~~~

#get data
sampleAllRNAseq <- fread('expression_data/rna_seq_all_genes.csv')

#get pheno
phenoDT <- data.table('sample'=paste("EM",1:18,sep=''),
                      'disease'='RA',
                      'celltype'='CD14+ monocytes')

phenoDT[grep("EM7|EM8|EM9|10|11|12|13|14|15",sample),disease:='healthy']
phenoDT[grep("EM2|EM5|EM8|11|14|17",sample),celltype:='CD16+ monocytes']
phenoDT[grep("EM3|EM6|EM9|12|15|18",sample),celltype:='intermediate monocytes']

#~~~~~~~~~~~
expMat <- as.matrix(sampleAllRNAseq[,grep("EM",colnames(sampleAllRNAseq)),with=FALSE])
row.names(expMat) <- sampleAllRNAseq$Symbol

#log transform [use pseudocount]
logExpMat <- log2((expMat+0.01))
row.names(logExpMat) <- sampleAllRNAseq$Symbol
colnames(logExpMat)  <- gsub("_FPKM","",colnames(expMat))

#format as data.table
expDT <- as.data.table(melt(logExpMat))
setnames(expDT,c("gene","sample","value"))
aovInputDT  <- merge(expDT,phenoDT,by='sample')

#~~~~~~~~~~~~~~~~~~~~~~~~~~~~~~~~~~~~~~~~~~~~~~~~~~~~~~~~~~~~~~~~~~~~~~~~~~~~~~~~~~~~~~
#Compute effect sizes [Classical / Non-classical]
#~~~~~~~~~~~~~~~~~~~~~~~~~~~~~~~~~~~~~~~~~~~~~~~~~~~~~~~~~~~~~~~~~~~~~~~~~~~~~~~~~~~~~~

#gene vector
mono_genes <- c("CSF3R",#CD114
                "FCGR2A",#CD32
                "CD36",
                "IL17RA",
                "FCGR3B",#CD16
                "SIGLEC10")

#discovery ES
discoveryES <- geneListMonoSubset[gene %in% mono_genes]#[grep("Non_Classical",cellType)]
discoveryES[cellType=='Classical',g:=-g]

#validation ES
validationDT <- aovInputDT[celltype!='intermediate monocytes']
validationDT[,class:=1]
validationDT[celltype=='CD14+ monocytes',class:=0]
esDT <- validationDT[,getES(value,class),gene]
esDT[,names:=rep(c("n1","m1","sd1","n2","m2","sd2","diff","pooled.sd","g",'se.g'),.N/10)]
esDT <- dcast.data.table(esDT[gene %in% mono_genes],gene~names,value.var = 'V1')

#summary ES
finalDT <- rbind(discoveryES[,c('gene','g','se.g')][,set:='Discovery'],
                 esDT[,c('gene','g','se.g')][,set:='Validation'])

wrapMeta <- function(g,se.g){
  aaa <- meta.summaries(g,se.g)
  return(c(aaa$summary,aaa$se.summary))
}

summaryDT <- dcast.data.table(finalDT[,wrapMeta(g,se.g),gene][,names:=rep(c("g","se.g"),.N/2)],
                              gene~names,value.var = 'V1')
summaryDT[,set:='summary']
summaryDT[,zs:=g/se.g]
summaryDT[,pv:=2*pnorm(abs(zs),lower.tail = F)]

#~~~~~~~~~~~~~~~~~~~~~~~~~~~~~~~~~~~~~~~~~~~~~~~~~~~~~~~~~~~~~~~~~~~~~~~~~~~~~~~~~~~~~~
#Make forest plots
#~~~~~~~~~~~~~~~~~~~~~~~~~~~~~~~~~~~~~~~~~~~~~~~~~~~~~~~~~~~~~~~~~~~~~~~~~~~~~~~~~~~~~~


pdf('pdfs/20201201_figure7_A.pdf',
   width = 4,height =4)
lapply(mono_genes,
       function(i){
         
         titleName <- i
         if(i=='CSF3R'){titleName='CD114'}
         if(i=='TNFRSF9'){titleName='CD13'}
         if(i=='FCGR2A'){titleName='CD32'}
         if(i=='FCGR3B'){titleName='CD16'}
         
         metaplot(finalDT[gene==i]$g,
                  finalDT[gene==i]$se.g,
                  nn = finalDT[gene==i]$se.g^-2,
                  finalDT[gene==i]$set,
                  summn = summaryDT[gene==i]$g,
                  sumse = summaryDT[gene==i]$se.g,
                  sumnn = summaryDT[gene==i]$se.g^-2,
                  main=titleName,
                  xlim=c(-max(abs(finalDT[gene==i]$g) + abs(finalDT[gene==i]$se.g)*2),
                         max(abs(finalDT[gene==i]$g) + abs(finalDT[gene==i]$se.g)*2)),
                  xlab="",ylab="")
         
       })
dev.off()
```
