## Supplementary figures and images for "Multicohort Analysis of Publicly-available Monocyte Expression Data Identifies Gene Signatures to Accurately Monitor Subset-specific Changes in Human Diseases"

### 20201201_figure2.pdf

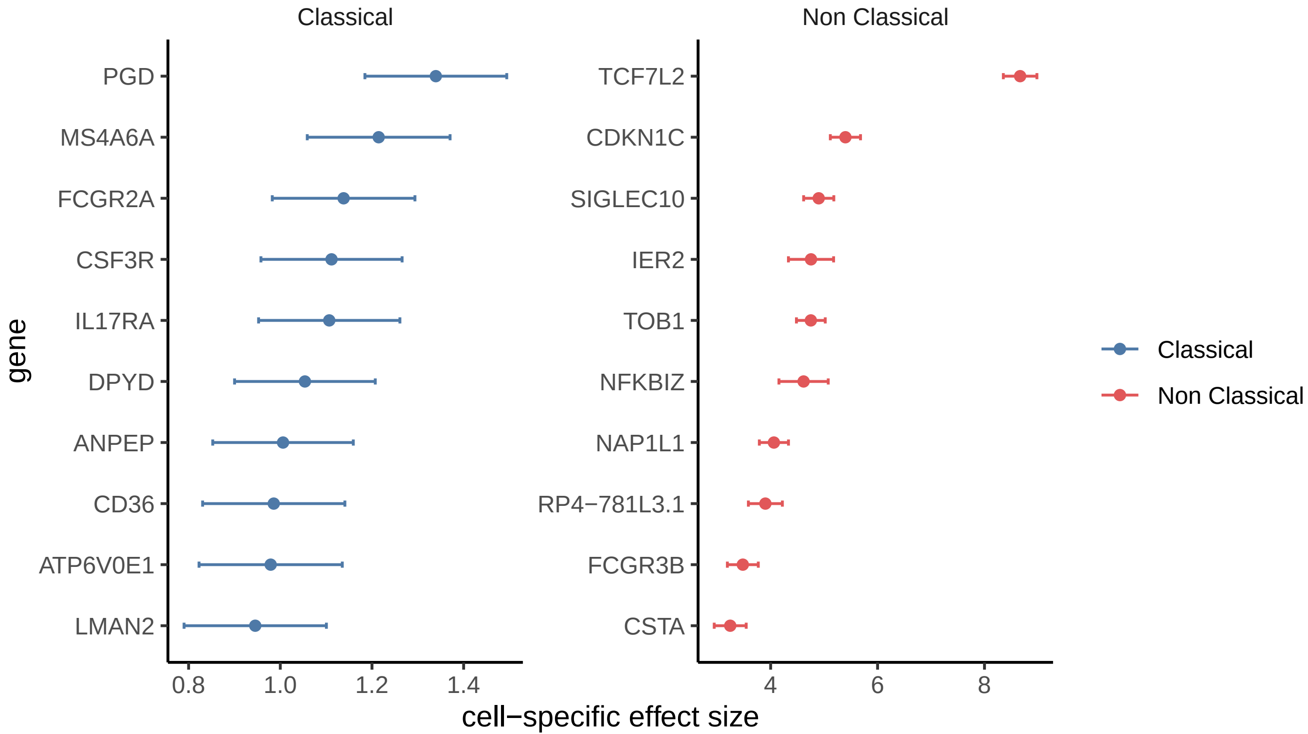

### 20201201_figure3_A.pdf

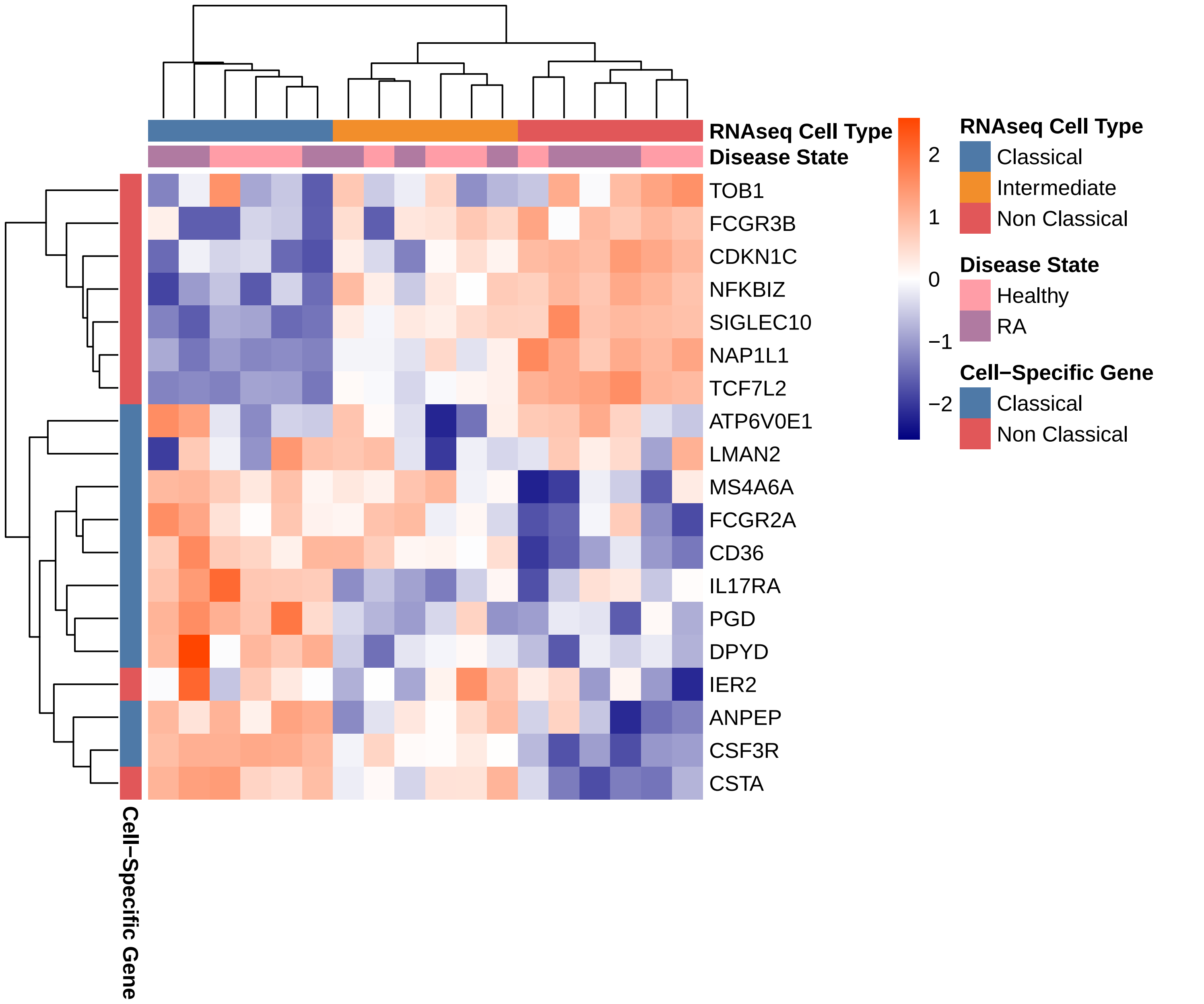

### 20201201_figure3_B_classical.pdf

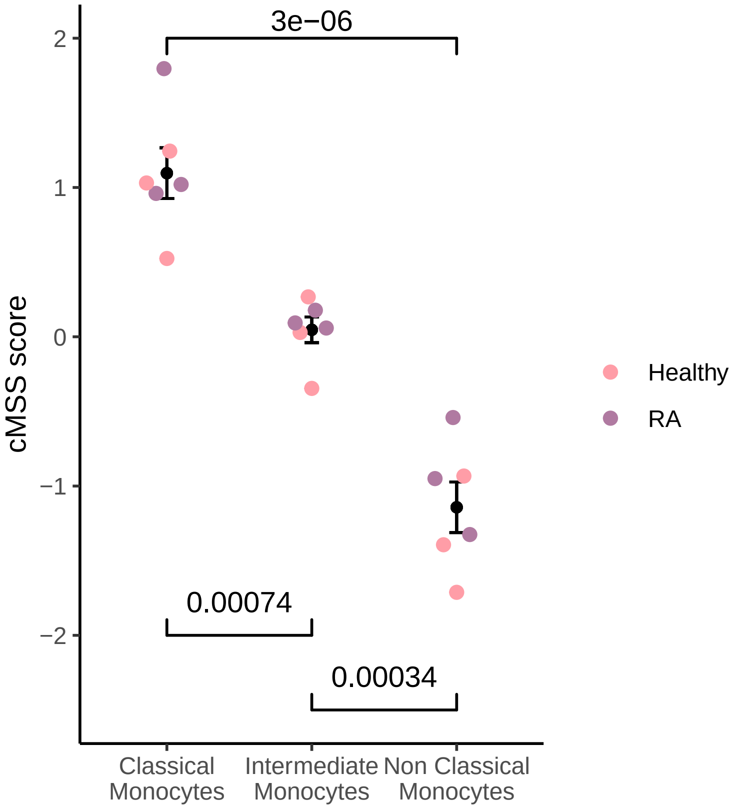

### 20201201_figure3_C_nonclassical.pdf

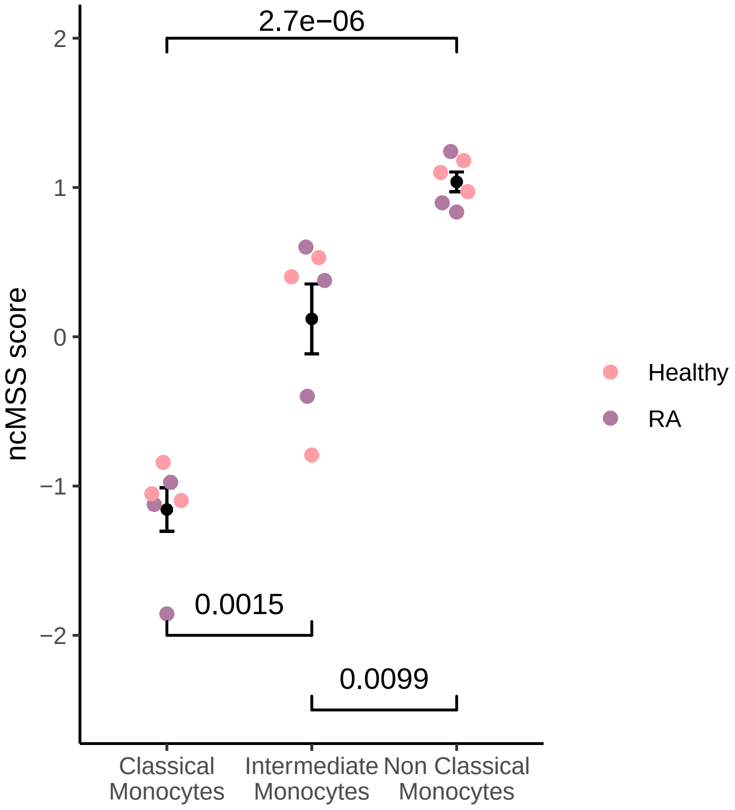

### 20201201_figure4_A.pdf

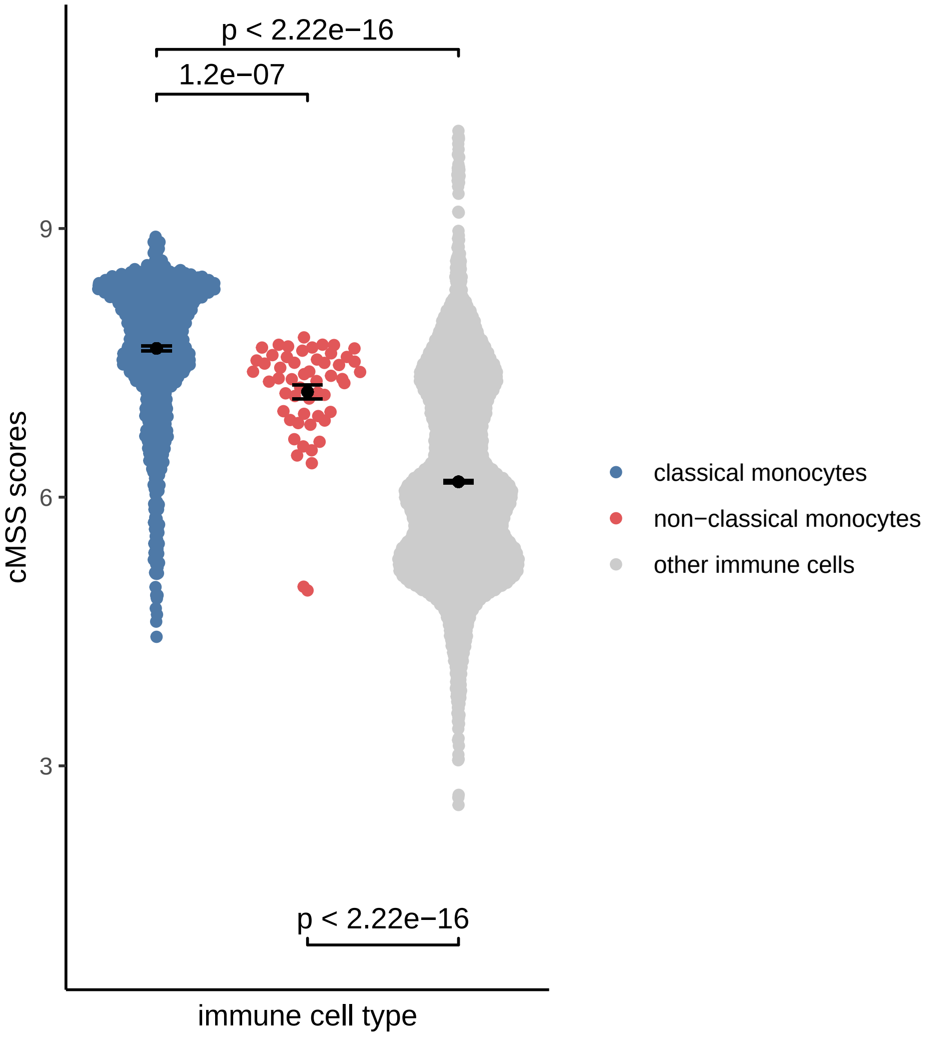

### 20201201_figure4_B.pdf

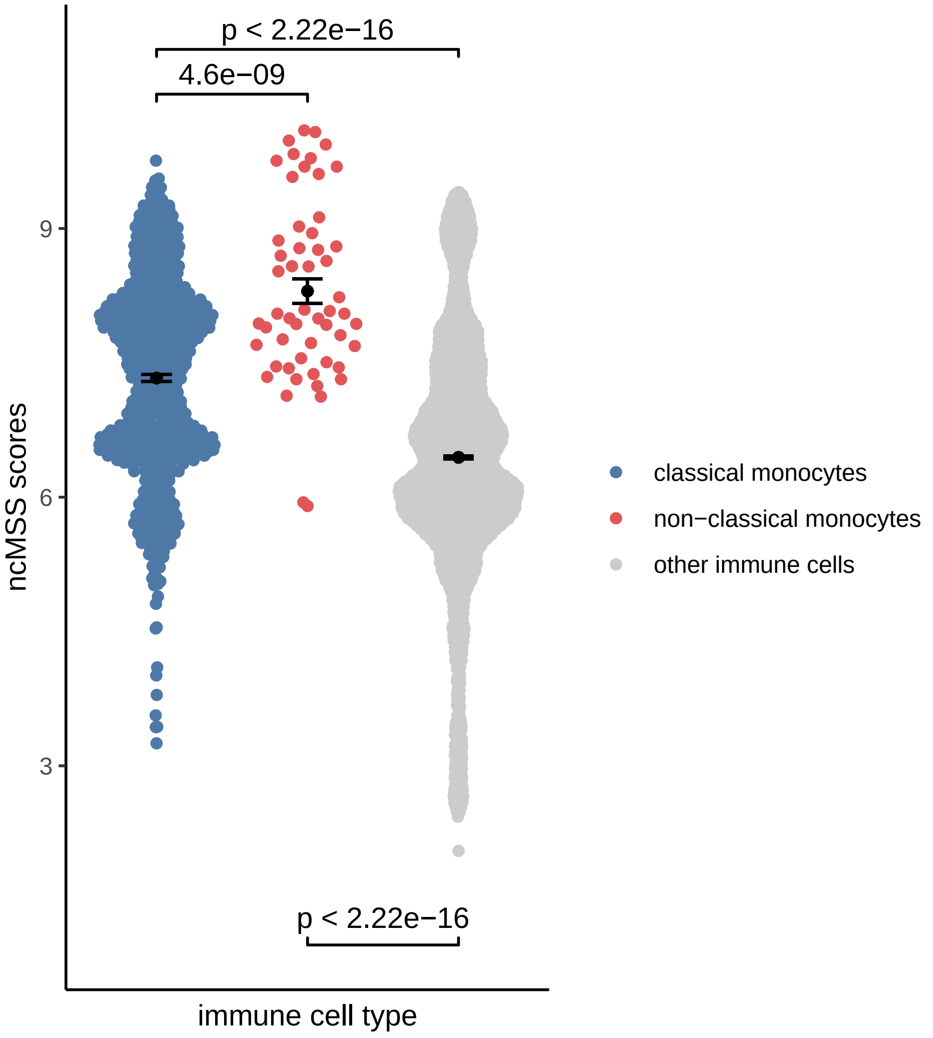

### 20201201_figure5_A_classical.pdf

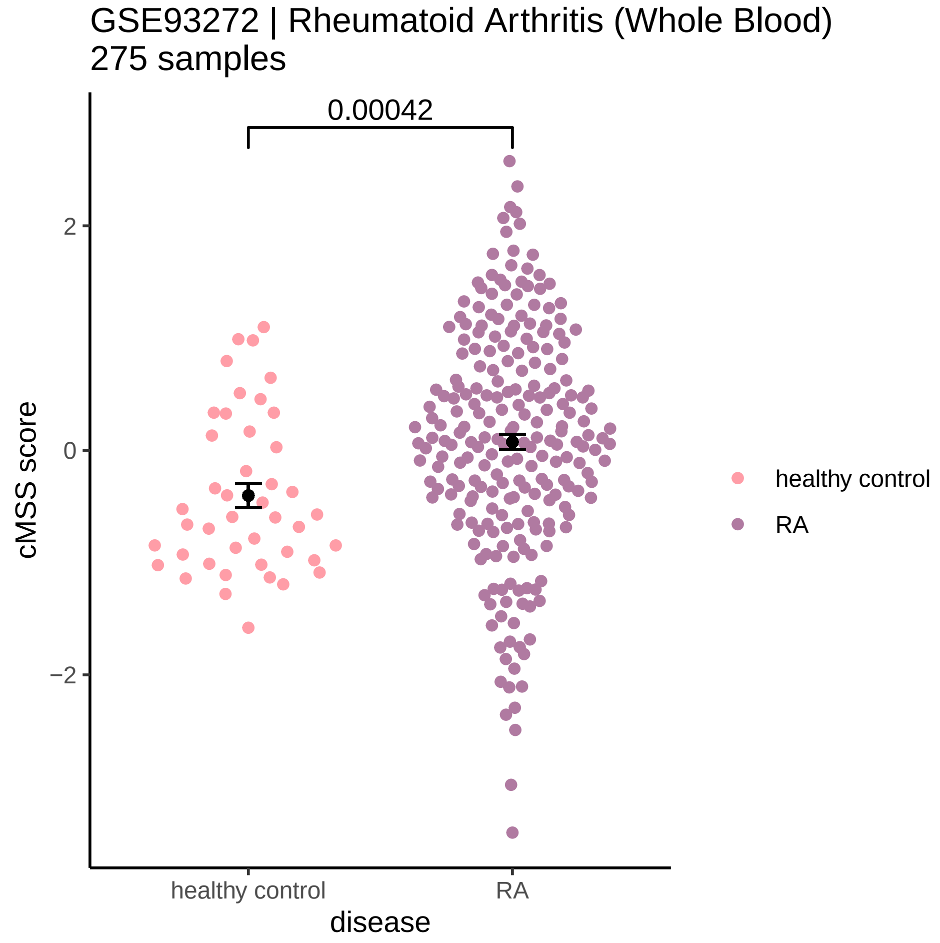

### 20201201_figure5_A_nonclassical.pdf

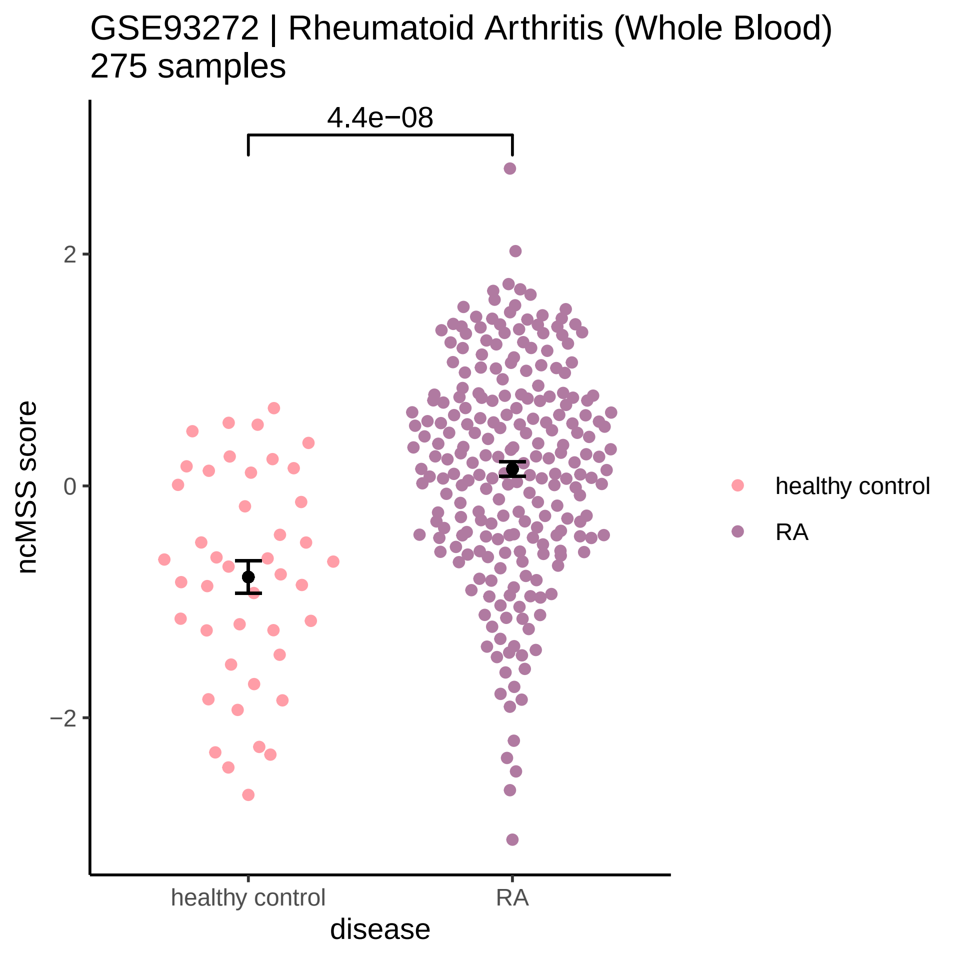

### 20201201_figure5_B_classical.pdf

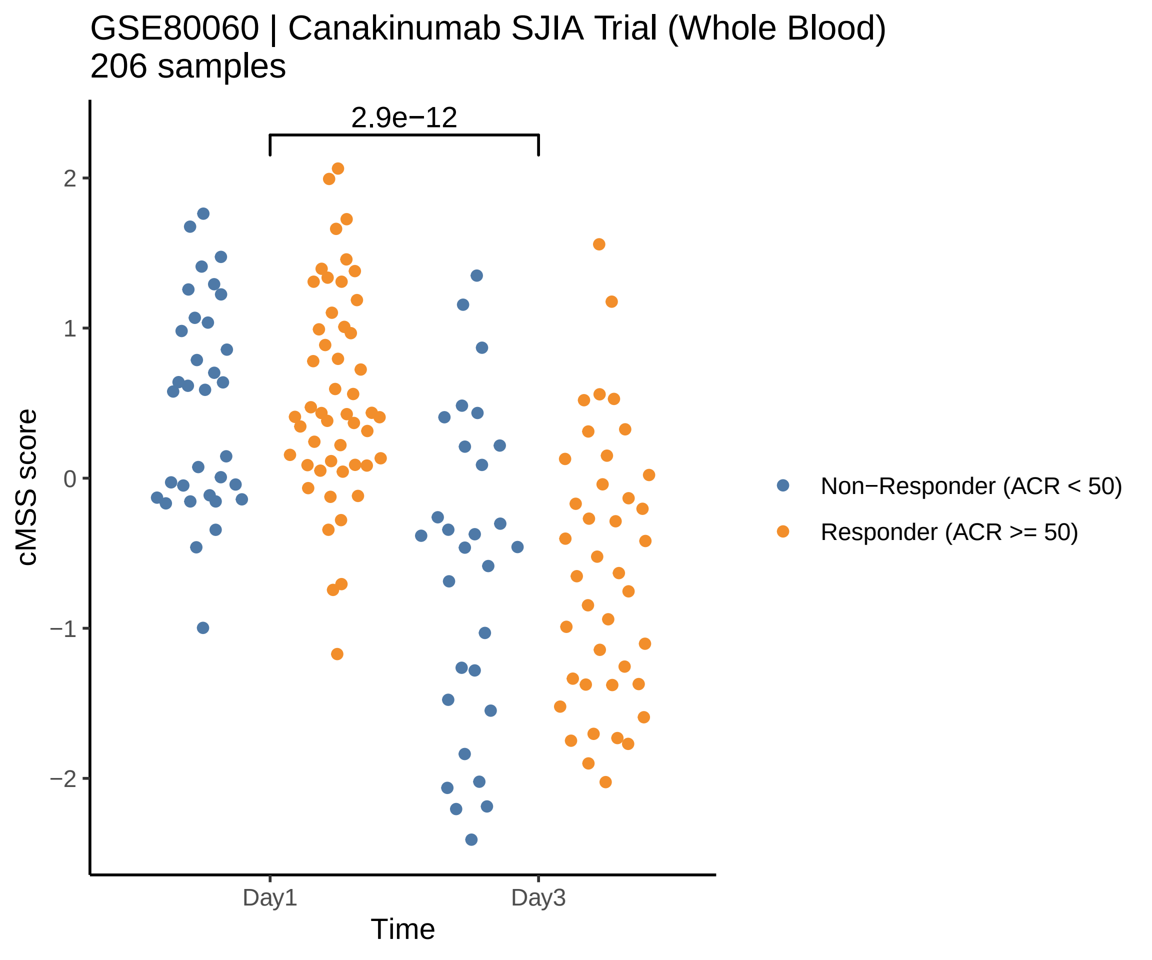

### 20201201_figure5_B_nonclassical.pdf

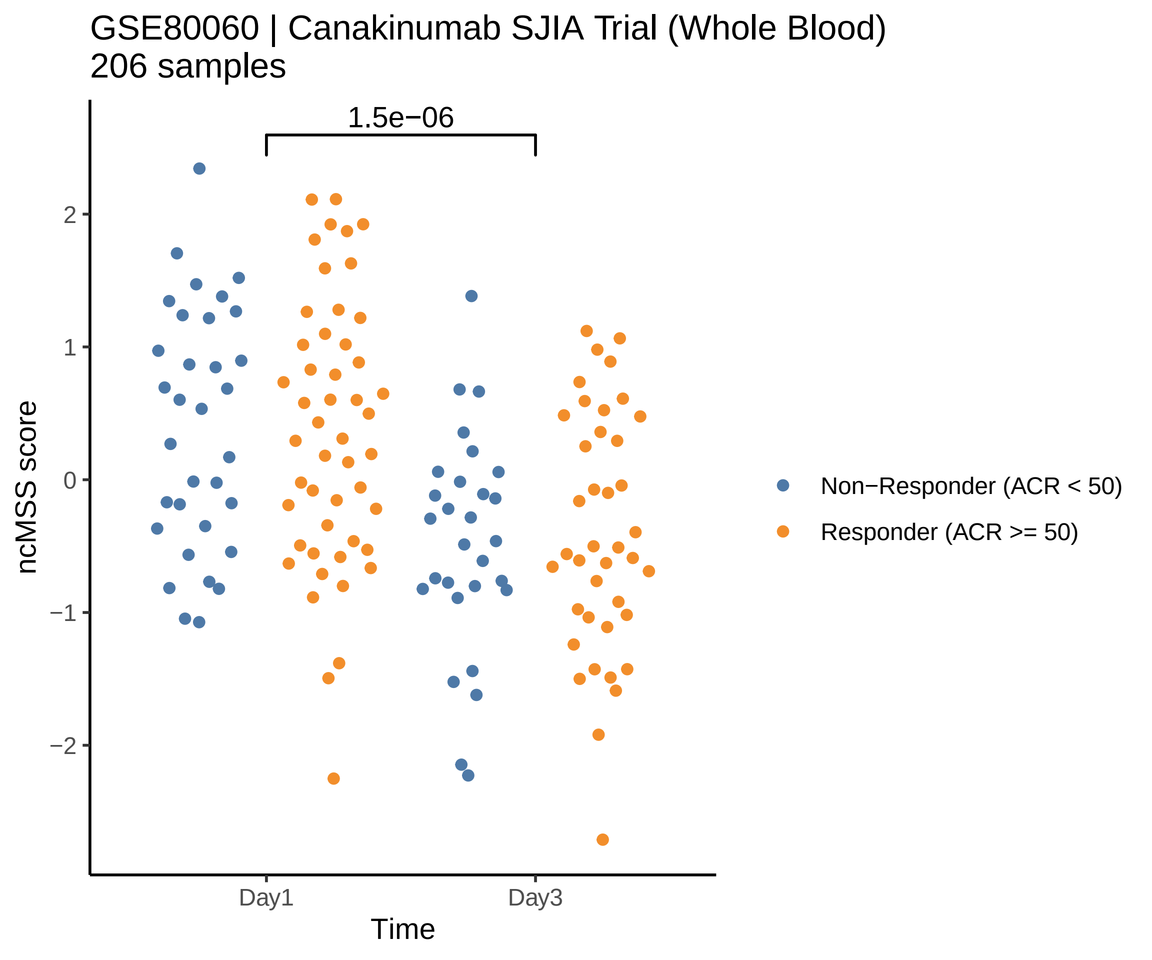

### 20201201_figure6.pdf

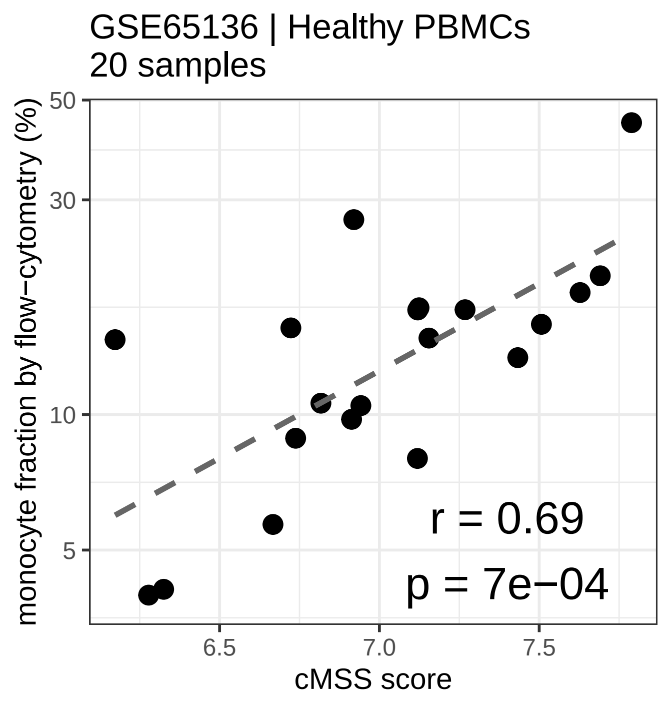

### 20201201_figure7_A.pdf

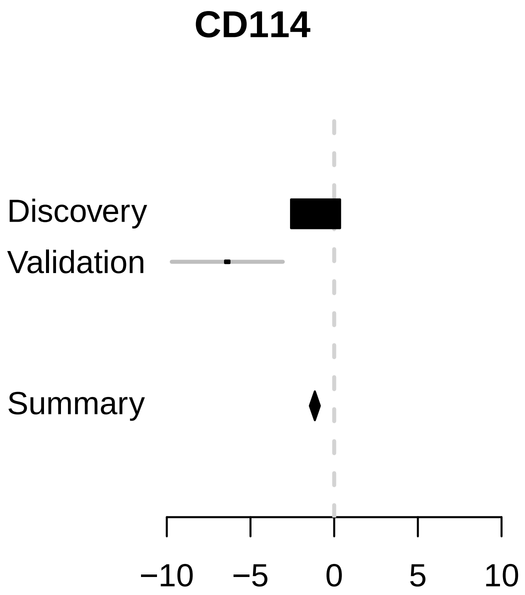

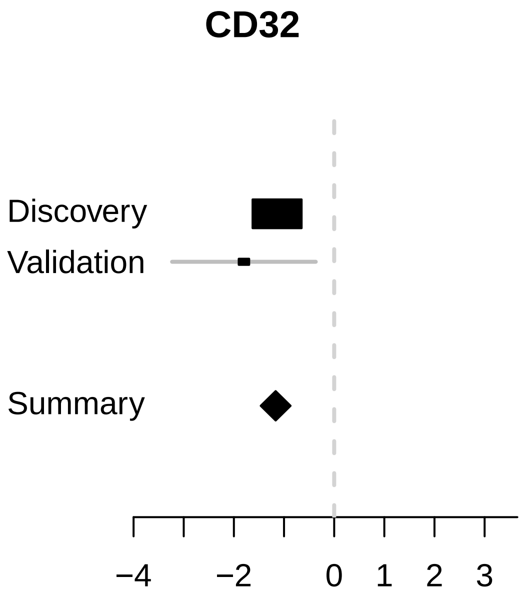

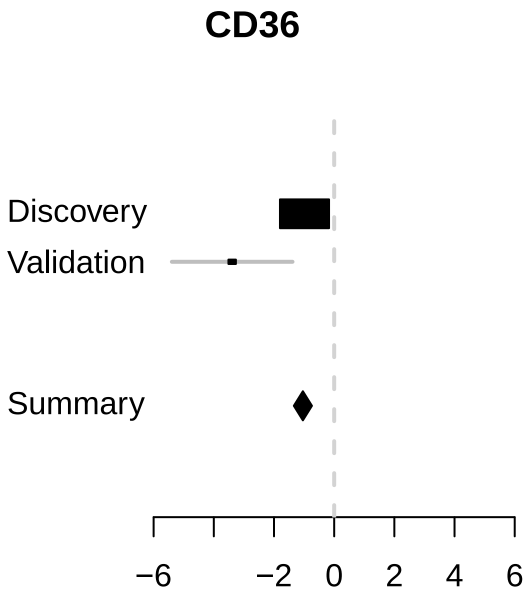

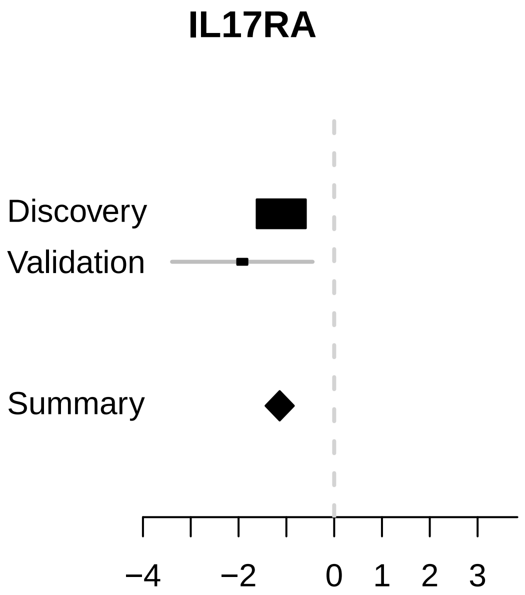

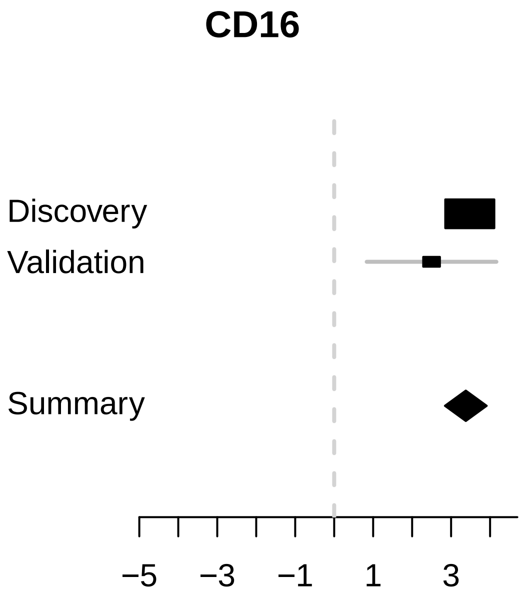

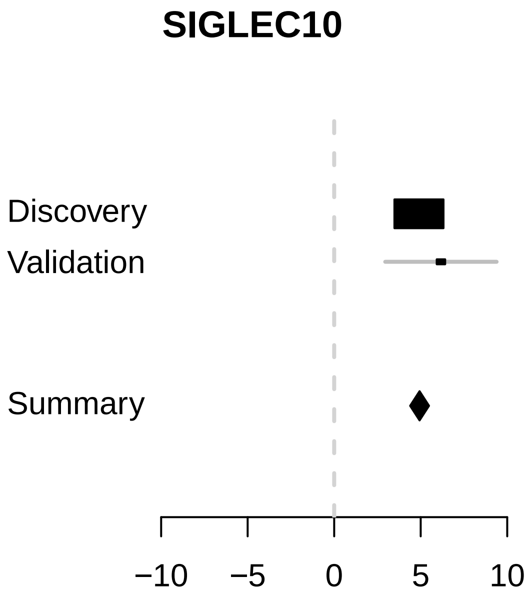

### 20201201_supplementary_figure2.pdf

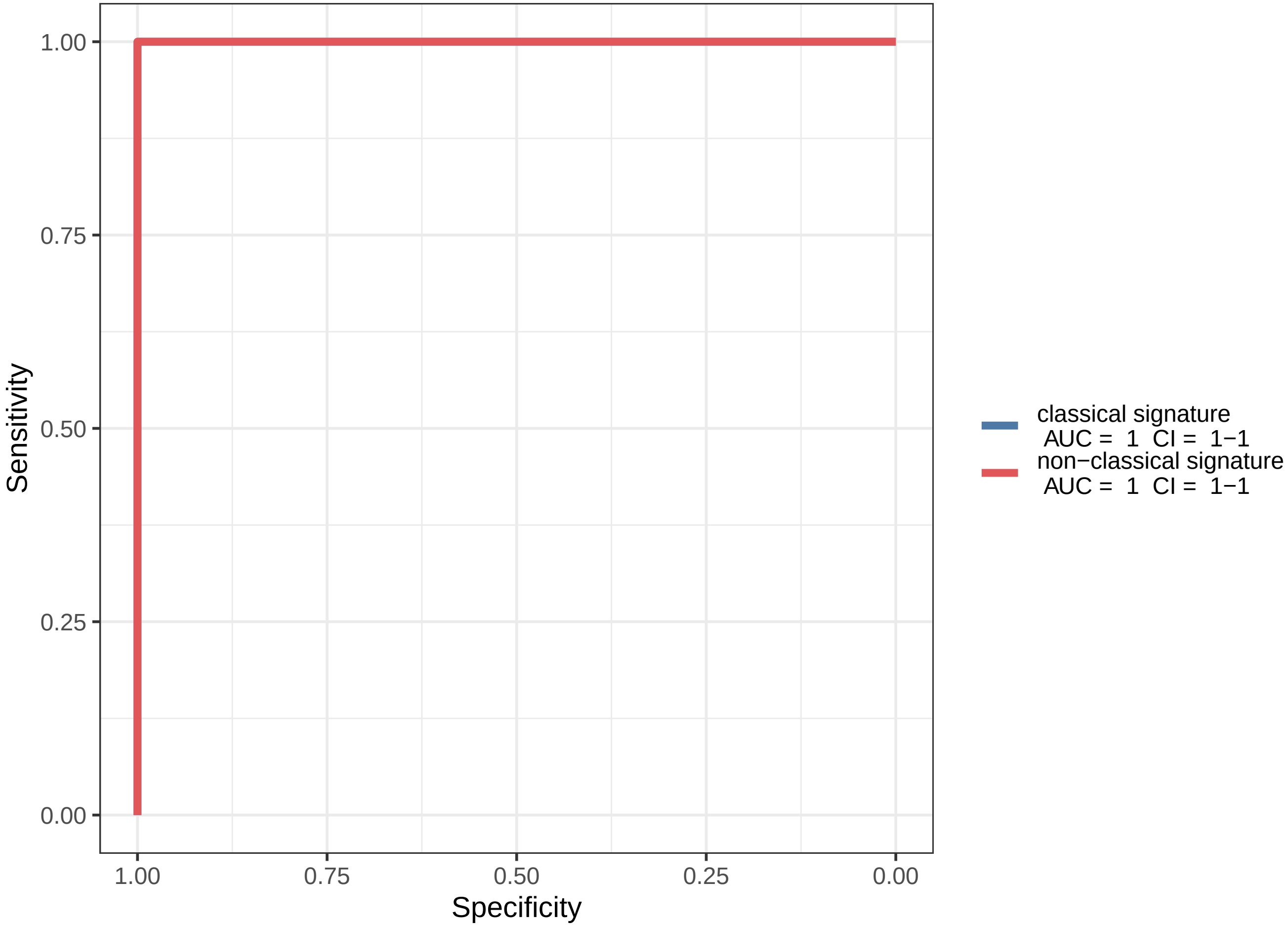

### 20201201_supplementary_figure3.pdf

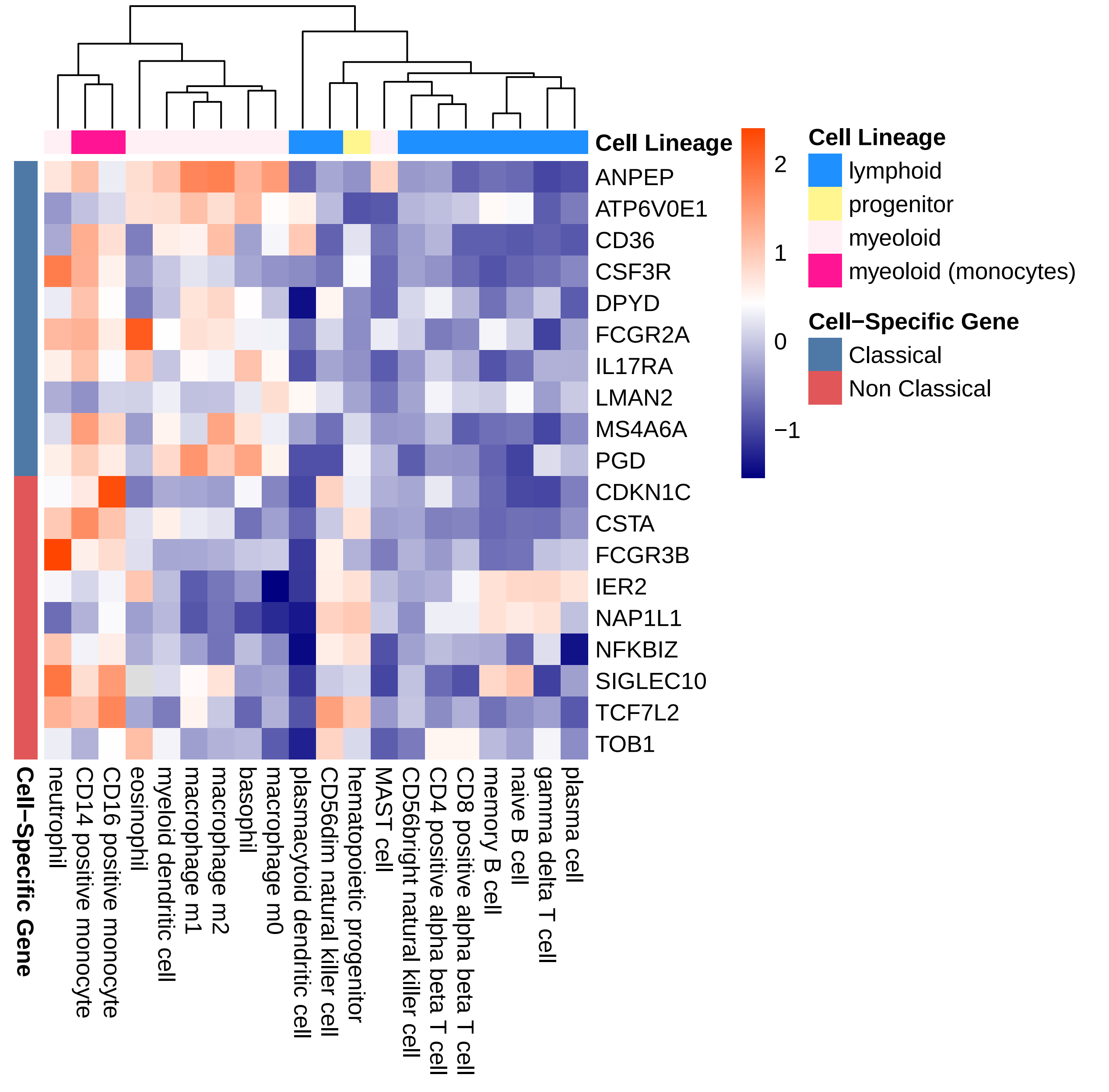

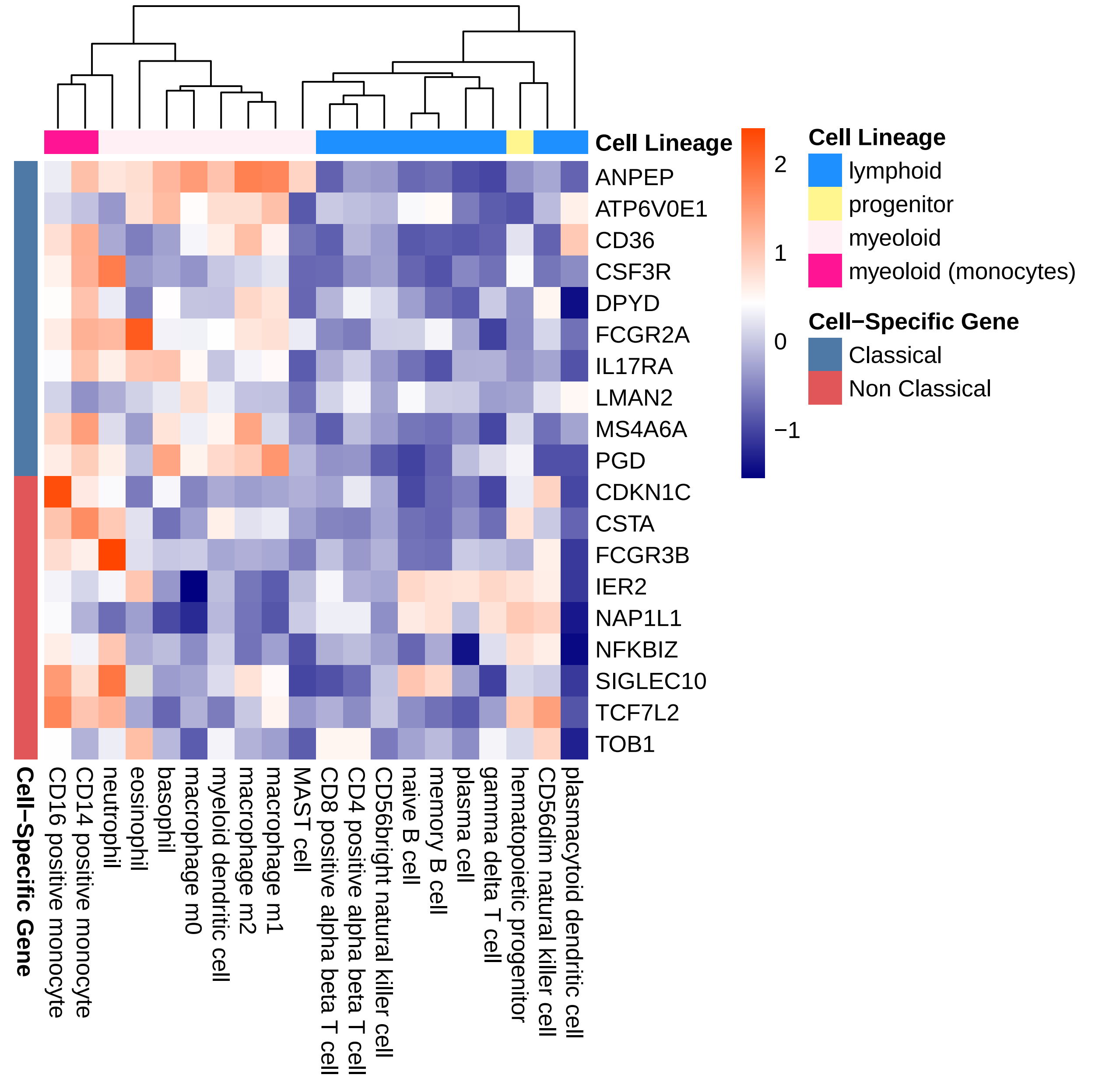

### 20201201_supplementary_figure4.pdf

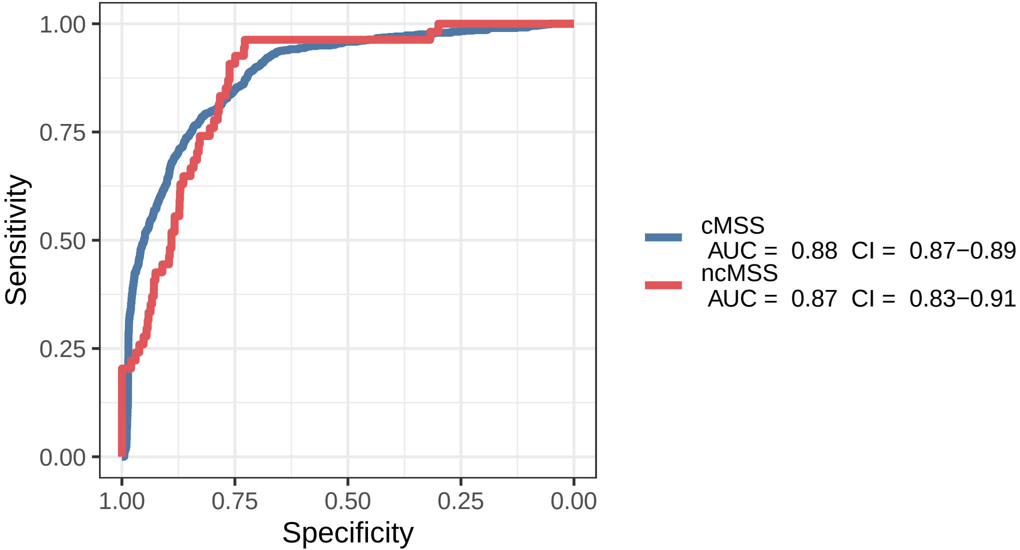

### 20201201_supplementary_figure6.pdf

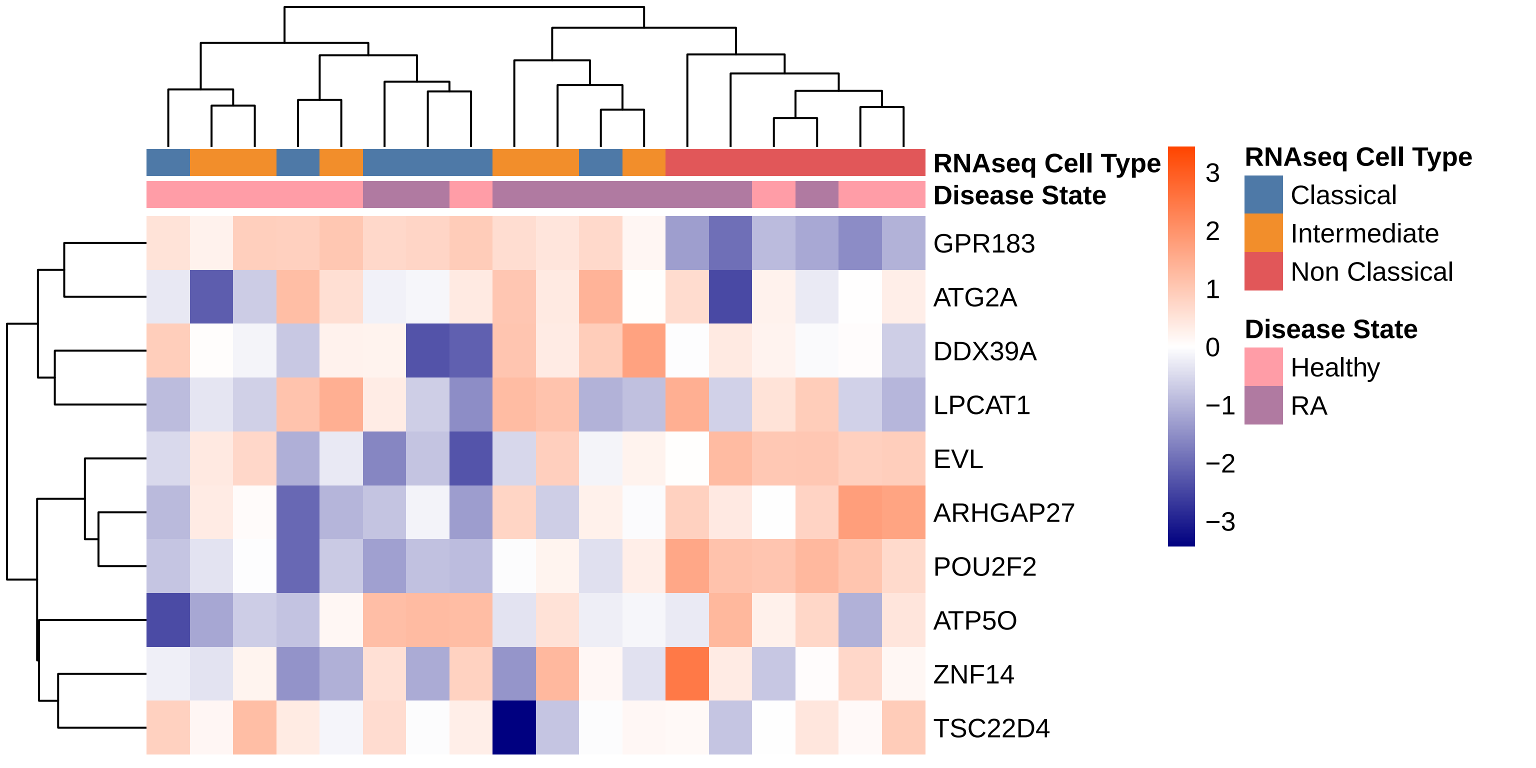
